# Synapse-like Specializations at Dopamine Release Sites Orchestrate Efficient and Precise Neuromodulatory Signaling

**DOI:** 10.1101/2024.09.16.613338

**Authors:** Chandima Bulumulla, Deng Zhang, Deepika Walpita, Nirmala Iyer, Mark Eddison, David Ackerman, Hideo Otsuna, Xianling Zhao, Shuqin Zhang, Shihong M. Gao, Nan Wang, Abraham G. Beyene

## Abstract

Neuromodulators are generally understood to act through expansive projections that broadcast signals relatively indiscriminately, in stark contrast to the precise, synapse-bound organization of fast neurotransmitters like glutamate and GABA. This dichotomy has left the local architecture of neuromodulatory release and receptor engagement largely obscure. Here, using dopamine as a model system, we reveal that neuromodulatory signaling can be guided by precise molecular specificity. We show that dopamine axons exhibit a fundamental tropism for neurons expressing D1 or D2 receptors in the striatum and amygdala, a targeting principle that is conserved in dissociated co-cultures. Importantly, single-bouton-resolved optical imaging of release shows that dopamine is not broadcast broadly but is highly localized to varicosities that form direct, synapse-like appositions with receptor-expressing somata and dendrites. In contrast, varicosities lacking such appositions, or contacting receptor-negative neurons, are largely quiescent, indicating that local interactions with receptor-expressing targets help specify dopamine release competence.

To directly visualize receptor organization relative to release sites at endogenous expression levels, we generated ALFA-tag knock-in mice (ALFADoR mice) in which the N-termini of D1 and D2 dopamine receptors are epitope-tagged. Combining these mice with immunolabeling and expansion microscopy in dissociated culture and intact tissue, we show that both D1 and D2 receptors are organized into discrete puncta rather than diffuse membrane distributions. Across major terminal fields of midbrain dopamine neurons, including striatum, amygdala, and prefrontal cortex, these receptor nanoclusters are preferentially positioned near dopamine varicosities at distances far smaller than expected from spatial shuffling controls.

Moreover, receptor clusters apposed to dopamine varicosities were observed to be larger than non-apposed clusters, consistent with specialized receptor microdomains that may enhance signaling efficacy.

Together, our data demonstrate that dopaminergic signaling is spatially organized by receptor-specific targeting and subcellular microdomain specialization. This structured framework challenges the canonical dichotomy between precise fast transmission and diffuse neuromodulation. It reveals that principles of synaptic organization extend to and define a previously hidden layer of specificity in neuromodulatory systems, opening new frontiers in understanding cell-type specific neuromodulation, axon guidance, and subcellular GPCR organization.

## Introduction

Two types of chemical signaling facilitate interneuronal communication in the brain. The first type, fast synaptic transmission, occurs between precisely coupled synaptic partners, where the activation of ligand-gated ion channels elicits depolarizing or hyperpolarizing electrical responses. The molecular and cellular mechanisms underlying fast synaptic transmission have been extensively studied (*1, 2*). The second type of communication involves neuromodulators, which are thought to be released volumetrically and act diffusively on their targets in a paracrine fashion (*3*). Although volumetrically released neuromodulators control the efficacy of fast synaptic transmission, and therefore play a critical role in neuronal computation that underlies cognition and behavior (*4, 5*), the precise nature of their release sites and signaling is less well understood and remains an active area of research (*6, 7*).

Historically, dopamine has served as the model system for neuromodulatory signaling.

The volume transmission framework, supported by early histochemical and ultrastructural studies, proposed that dopamine is released from axonal varicosities lacking synaptic specializations, allowing it to diffuse freely through extracellular space to activate distant receptors (*8–11*). Yet this interpretation has remained controversial. While some electron microscopy studies reported that most dopamine axons lack classical synapses (*12–14*), others described synapse-like appositions between dopamine terminals and postsynaptic processes (*15–18*). Moreover, physiological and behavioral studies have shown that dopamine neuron firing can influence behavior with millisecond-to-second precision (*19–23*). These temporal dynamics are difficult to reconcile with purely diffuse signaling. These discrepancies raise a fundamental question: does dopamine signaling employ synaptic mechanisms to achieve spatial and temporal specificity, and if so, how are these organized at the cellular level?

A key limitation underlying these discrepancies is the lack of functional data correlating dopamine release dynamics with synaptic ultrastructure and receptor localization. Prior work either mapped dopamine varicosities histologically without functional activity data (*12, 24–26*), or measured extracellular dopamine dynamics at low spatial resolution, obscuring single-varicosity mechanisms (*27, 28*). Consequently, it remains unclear whether dopamine release is stochastic or targeted, and whether synaptic architecture enhances neuromodulatory precision. Here, we combine single-bouton optical recording of dopamine release with correlated immunofluorescence, transcriptomics, and expansion and electron microscopy, together with dopamine receptor profiling in newly generated transgenic knock-in mice, to address these gaps. We show that dopaminergic axons preferentially form synapse-like contacts with neurons expressing dopamine receptors, and that release competence strictly depends on anatomical apposition. Varicosities lacking such appositions, or contacting receptor-negative processes, remain quiescent. Functionally active varicosities contain canonical presynaptic specializations (Bassoon, RIM, Munc13-1 & others), are juxtaposed against dopamine receptor clusters visualized in our ALFA-tag knock in (ALFADoR) mice, and evoke localized cAMP responses in their targets. These findings suggest that dopamine neuromodulation employs a hybrid signaling modality, integrating synaptic architecture for precision and efficiency while maintaining the biochemical complexity of GPCR signaling.

## Results

### Dopamine is Released from Varicosities that are Apposed to Neuronal Dendrites and Somata

Our study employs two complementary nanomaterial-based fluorescent dopamine sensors: DopaFilm and nIRCat. DopaFilm consists of optical dopamine nanosensors drop-casted onto 2-dimensional glass substrates (*29*), while nIRCat is a solution-phase version used in intact tissue slices (*30*). Both nanosensor modalities exhibit rapid, intensiometric fluorescence modulation in response to dopamine. For experiments on DopaFilm, we generated primary dopaminergic (DA) neurons from P1–P2 pups of *Slc6a3*-Cre (DAT-Cre) mice crossed with Ai9 reporter mice. DA neurons were co-cultured with either cortical (DA:C) or striatal (DA:S) neurons. By culturing a sparse monolayer of primary neuron co-cultures on DopaFilm, we imaged evoked and spontaneous dopamine release from dopamine neurons with single bouton resolution (Fig. 1A – B). DA hotspots on DopaFilm had a mean size of 2.43 ± 0.91 µm, latency to peak (rise time) of 0.66 ± 0.42 s, a peak ΔF/F of 14.8 ± 3.2%, and cleared with a first order time constant (τ) of 10.57 ± 4.85 s (all Mean ± SD). We corroborated this hotspot-like organization of dopamine release using nIRCat by imaging release from dopamine axons in the dorsomedial striatum (DMS) and basolateral amygdala (BLA) in acute slice preparations (Fig. 1C). While hotspots in the DMS and BLA were comparable in size to those in culture (2.12 ± 0.72 µm and 2.35 ± 0.81 µm respectively), we observed differences in peak ΔF/F (DMS: 5.02 ± 2.02%, BLA: 7.58 ± 4.11%), clearance kinetics (DMS: 7.87 ± 2.42 s, BLA: 19.61 ± 12.41 s), and latency to peak (DMS: 1.19 ± 0.51 s, BLA: 2.69 ± 1.31 s) (all Mean ± SD) (Fig. 1D – E). These data demonstrate spatially confined, hot-spot like organization is a key hallmark of extracellular dopamine dynamics, and that dopamine nanosensors enable the visualization of these dynamics in both culture and intact tissue, capturing subtle spatial and temporal differences that likely reflect distinct local molecular and ultrastructural environments. Additionally, DA:S co-cultures exhibited significantly greater axonal arborization, measured by total axonal length, branching points, and other morphometric parameters compared to DA:C co-cultures (Fig. S1). This morphological distinction is consistent with the known in vivo characteristics of their putative anatomical representations. In contrast to mesocortical projections, the mesostriatal pathway, primarily comprised of substantia nigra pars compacta (SNc) neurons, is characterized by fine axonal fibers that form dense and highly branched arbors (*31*). Taken together, these data show that DopaFilm recapitulates key characteristics of extracellular neurochemical dynamics and axonal morphology while reporting activity with subcellular resolution.

**Fig. 1.**
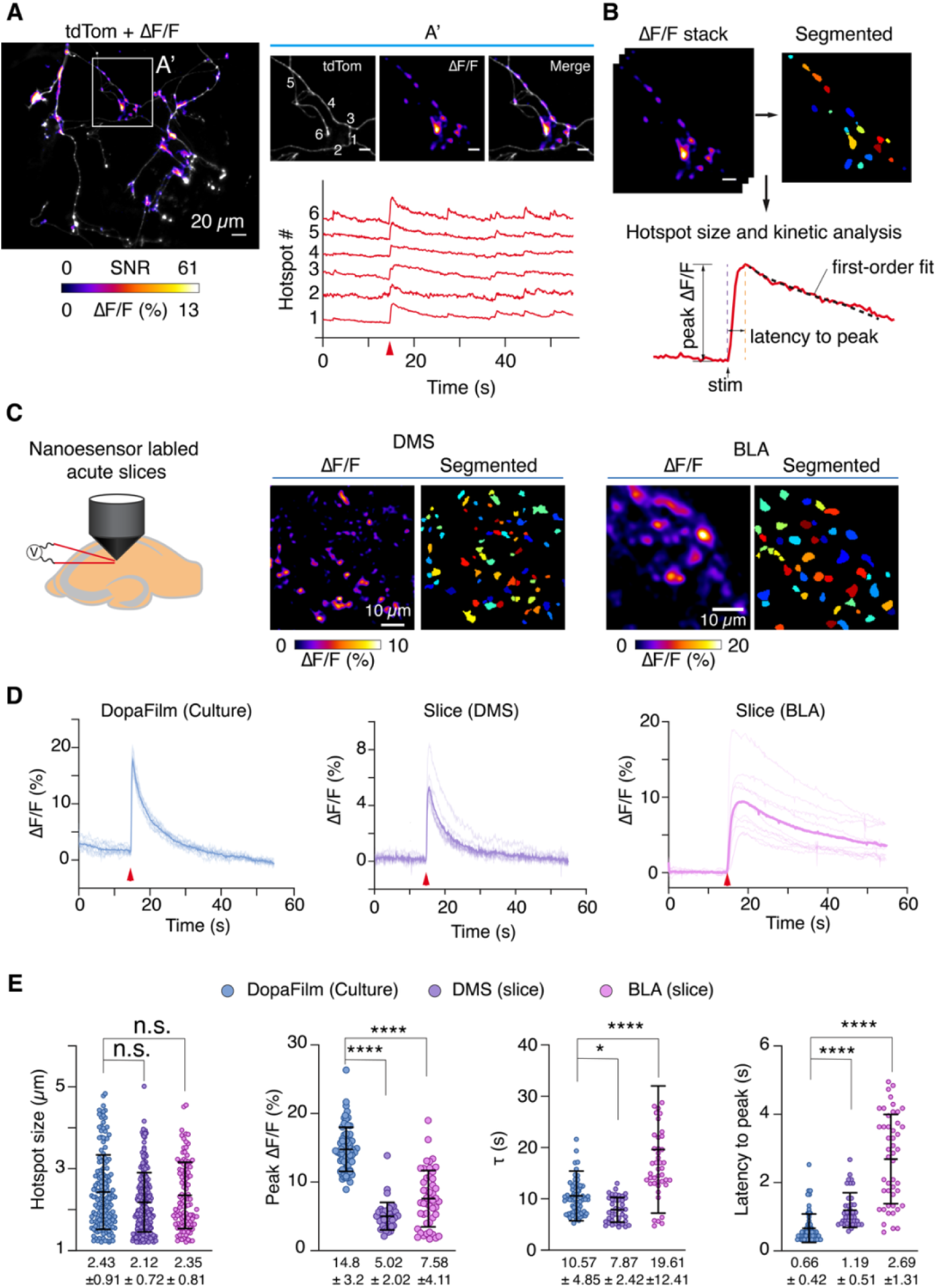
Imaging dopamine release hotspots in primary cultures and acute slices. (A) tdTomato (tdTom) expressing dopaminergic axons overlaid with dopamine ΔF/F hotspots. The ΔF/F frame is taken ∼300 ms after application of electrical field stimulation. Right: Close up of A’ depicted in (A), scale bar is 10 µm. Bottom: activity traces corresponding to varicosities 1– 6 showing a mix of electrically-evoked (red arrow) and non-evoked (i.e., spontaneous) release events. (B) Analyzing extracellular DA dynamics: hotspots are segmented from ΔF/F stack, and temporal trace corresponding to each hotspot is extracted for further analysis; peak ΔF/F, τ from a first order decay and rise time calculated as ‘latency to peak’ are evaluated. (C) Solution phase dopamine nanosensors (nIRCat) enable visualization of dopamine release hotspots in the dorsomedial striatum (DMS) and basolateral amygdala (BLA), which receive axonal projections from the substantia pars compacta (SNc) and ventral tegmental area (VTA), respectively. Activity is electrically evoked (1 pulse with bipolar electrodes) in 300 µm thick slices. (D) Representative temporal traces from DopaFilm, DMS and BLA. Each panel shows mean trace (bold) and individual traces from 10 randomly selected hotspots in the imaging field of view. Data from one imaging field is shown as an example. (E) Comparison of hotspot sizes, peak ΔF/F (evoked), clearance kinetics (τ) and latency to peak. Each data point represents hotspot pooled from n = 10 biological replicates. Mean ± SD values are shown at the bottom of each chart. Test of statistical significance: non-parametric Kolmogorov-Smirnov (KS) test. n.s. = not significant, p-values: * < 0.05, **** < 10^-4^.

Following live-cell imaging on DopaFilm, we conducted post hoc multimodal analyses of dopamine axonal varicosities and their putative targets using immunofluorescence microscopy, spatial transcriptomics, and serial-section transmission electron microscopy (Fig. 2A). With this approach, we were able to correlate functional activity data (i.e., DA hotspots) with ultrastructural and molecular features with single-bouton precision. Our analyses of DopaFilm data revealed *three* principal findings regarding dopamine release organization in axonal arbors.

**Fig. 2.**
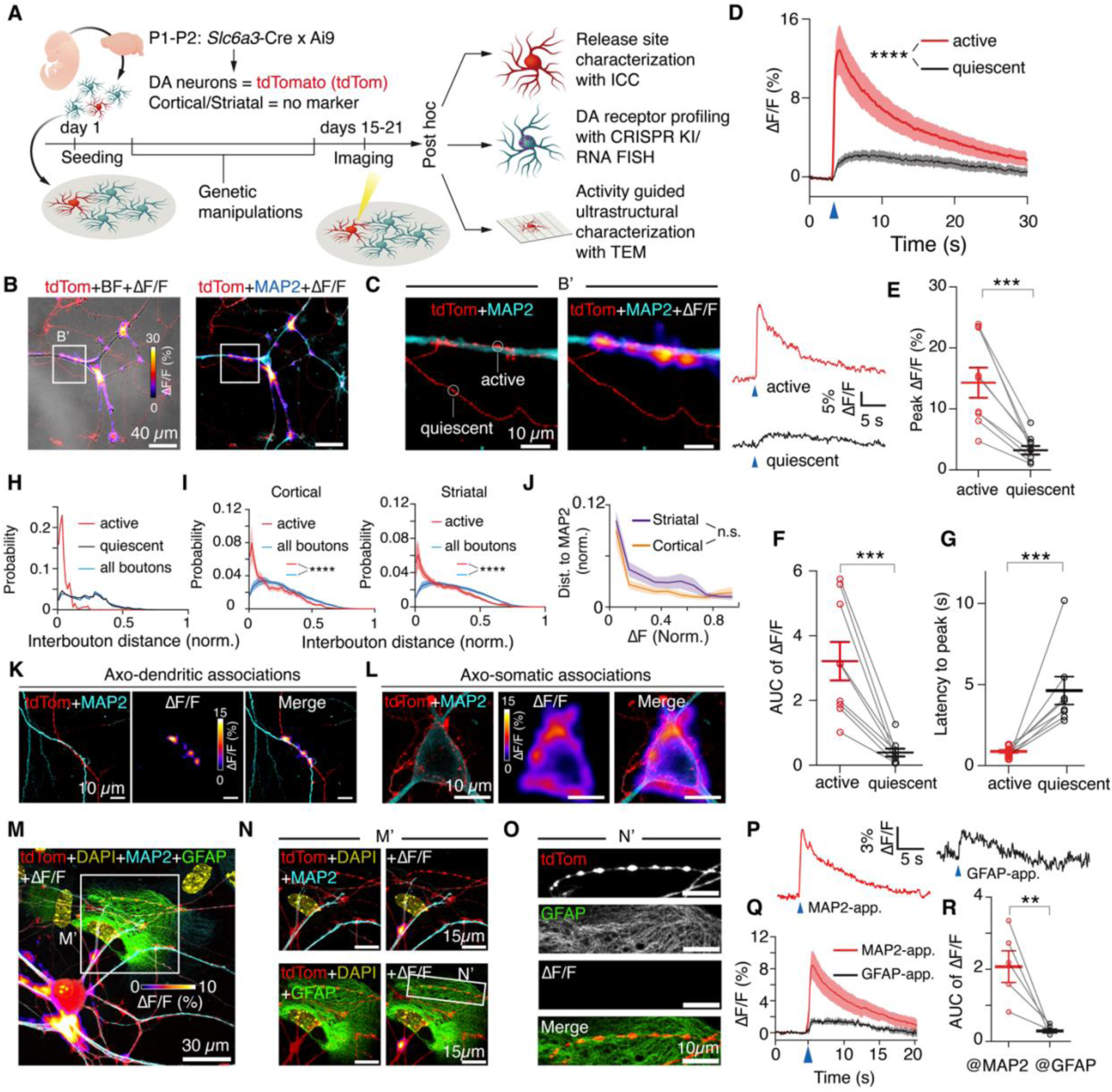
Experimental schematics and DopaFilm activity imaging in dopamine neuron axonal arbors. (A) Experimental workflow. Dopamine neurons and striatal or cortical neurons are extracted from transgenic mice pups at P1–P2 and co-cultured on DopaFilm, and are processed for post hoc analysis after live-cell imaging. (B) An axonal arbor of a dopamine neuron expressing tdTomato (red, tdTom) in co-culture with a cortical neuron that is seen in bright field (BF) image during live-cell imaging. Evoked dopamine activity was captured by DopaFilm and overlaid with the BF and tdTom images (heat map). Right: confocal images of the same field of view (FOV) after post-hoc immunocytochemistry (ICC) against MAP2 (cyan). (C) Magnified view of region of interest (ROI) (B’, white box) depicted in (B). Note that the active varicosities are apposed to and clustered near the dendrite (MAP2) and non-apposed varicosities do not release dopamine (quiescent). Right: activity traces corresponding to the representative active and quiescent varicosities depicted on the left panel. Blue arrow heads: time of field stimulation to evoke release. (D) Average activity traces from active and quiescent varicosities pooled from n = 9 biological replicates. Each replicate contributes a mean trace from n = 10 active and n = 10 quiescent varicosities. Statistical test of significance was performed using non-parametric, two-sample Kolmogorov-Smirnov (KS) test; p-value: **** < 10^-4^. (E, F, G) Peak ΔF/F, area under the curve (AUC) and latency to peak for active and quiescent varicosities from n = 9 biological replicates. Each data point represents a mean value from n = 10 active and n = 10 quiescent varicosities and Mean ± SEM bars are shown. AUC is computed over a 3-second window in the post-stim epoch. Statistical test of significance is performed using KS test; p-value: *** < 10^-3^. (H) Clustering analysis: distribution function of pair-wise distances between release-active boutons (red), quiescent boutons (black) and all boutons (blue). This data corresponds to the FOV shown in Fig. 2B. (I) Pooled clustering analysis. Distribution function of interbouton distances at active boutons (red) vs. all boutons (blue) for DA-cortical and DA-striatal culture. Each trace represents average form n = 18 (DA:C) and n = 25 (DA:S) biological replicates. Distances are normalized to enable direct comparison across FOVs of different sizes. Mean traces and SEM bands are shown. Statistical test of significance is performed using KS test, p-value: **** < 10^-4^. (J) Proximity analysis: plot of normalized ΔF value at a dopaminergic bouton as a function of its distance from a MAP2+ process. Higher ΔF regions occur close to MAP2+ processes. Each trace represents average of from n = 18 (DA:Cortical) and n = 25 (DA:Striatal) samples. DA:Cortical and DA:Striatal cultures exhibit a similar trend. Mean traces and SEM bands are shown. Statistical test of significance is performed using KS test, n.s.= not significant. (K) FOV consisting of dopamine axons (red, tdTom) innervating a dendrite of a cortical neuron (MAP2). Note that only dendritically-apposed varicosities release dopamine. (L) Axons of a dopamine neuron (red, tdTom) innervating a cortical neuron soma (MAP2), visualized with Airyscan imaging. Note that the soma-apposed varicosities participate in dopamine release whereas non-apposed ones do not. (M) FOV containing a tdTom positive dopamine neuron (soma and dendrites are in view) immunostained against MAP2 (cyan) and GFAP (green) and co-stained with nuclear marker DAPI (yellow). (N) Enlarged view of ROI depicted in M (M’, white box) with overlaid images of tdTom, DAPI, MAP2 and ΔF/F heatmap (top), and overlaid images of tdTom, DAPI, GFAP and ΔF/F heatmap (bottom). Note that dopamine release is evident at MAP2+-apposed varicosities but not at varicosities interacting with GFAP+ glial processes. (O) Magnified view of ROI depicted in N (N’, white box). tdTom, GFAP, ΔF/F and merged images are presented. The tdTom varicosities do not participate in release despite their proximity to glial processes. (P) Representative activity traces at MAP2-apposed and GFAP-apposed varicosities. Blue arrow heads: time of field stimulation to evoke release. (Q) Activity traces at MAP2-apposed and GFAP-apposed varicosities averaged over n=5 biological replicates. (R) Area under the curve of activity traces at MAP2-apposed and GFAP-apposed varicosities. Each point represents a mean value from activity at n = 10 varicosities, averaged from n = 5 samples, and Mean ± SEM bars are shown. Statistical test of significance is performed using KS test: p-value: ** < 10^-2^.

First, we observed that dopamine release originates from a subset of varicosities, with considerable segments of dopamine axonal processes remaining release-incompetent (quiescent) (Fig. 2B – 2C). The release-competent (active) varicosities, which constituted a smaller proportion of varicosities within an imaging field of view (34 ± 27 % were active, Mean ± SD), were distinguished from quiescent varicosities because of their higher ΔF/F (14.7 ± 2.5% vs. 3.65 ± 0.01%, Mean ± SEM, Fig. 2D – E), larger area under the ΔF/F curve (3.25 ± 0.59 vs. 0.42 ± 0.12, Mean ± SEM, Fig. 2F), and sharper, stimulation-locked responses (Fig. 2C, right panel). In contrast, quiescent varicosities exhibited smaller ΔF/F that often fell below the limit of detection, and when dopamine signals were measured, they were characterized by long latency to peak (0.81 ± 0.1 s vs. 4.55 ± 0.86 s, Mean ± SEM, Fig. 2G), suggesting detected signals likely reflect diffusive spill over from neighboring release sites rather than local exocytosis. These characteristic differences were also observed during spontaneous (cell-autonomous) release events (Fig. S2), and persisted even under vigorous stimulation conditions (Fig. S2B – C), indicating that the observed differences reflect intrinsic release competency differences between active and quiescent processes.

Second, active varicosities were spatially clustered, resulting in a distinctly non-uniform dopamine release profile across imaging fields (Fig. 2B, Fig S2A). To quantify this organization, we analyzed the distribution of pairwise distances between dopaminergic (i.e., tdTom+) varicosities (boutons). The distribution of pairwise-distances of active boutons displayed a prominent peak at small interbouton distances (10.5 ± 7.9 µm, Mean ± SD) contrasting with the broad, featureless distribution of quiescent boutons and all dopaminergic varicosities (Fig. 2H). This clustering was consistent across both DA:C and DA:S co-cultures (Fig. 2I). If release competence in dopamine axons is stochastically distributed, as is currently understood, the pair-wise distribution of active boutons would be expected to resemble the overall distribution of inter-bouton distances. However, this is not observed in our data. Thus, this analysis suggested a non-arbitrary and structured arrangement of release competence in axonal varicosities of dopamine neurons.

Third, release competence strongly correlated with anatomical apposition to neuronal processes. Immunostaining for the somatodendritic marker MAP2 revealed that active boutons were preferentially localized near dendrites or soma of co-cultured non-DA neurons (Fig. 2K and 2L, respectively). Normalized ΔF/F amplitudes decayed with increasing distance from MAP2+ processes, with the highest release amplitudes occurring at sites of direct juxtaposition between varicosities and dendrites/soma in both DA:C and DA:S cultures (Fig. 2J).

We identified two predominant configurations of release competent axons: those that were dendritically apposed (Fig. 2K), and those that are soma-apposed (Fig. 2L). Axo-somatic appositions were more frequently observed in DA:C co-cultures (72%, 36 out of 50 target cortical neurons had DA axons terminating on cell bodies and proximal dendrites) and we found that these innervations were often high-efficacy dopamine release sites (Fig. 2L, Animation 1) whereas in DA:S co-cultures, the innervations were predominantly dendritic (70%, 35 out of 50 striatal cells preferentially received dendritic DA input). Importantly, inspection of dopamine secretion along the contours of individual axonal processes revealed that release competence is highly sensitive to proximity to and appositional contact with neuronal processes (Fig. S3A – B). Notably, this spatial coupling was specific to neuronal processes; proximity to GFAP+ glia did not enhance release competence (Fig. 2M – O, Fig. S3C), and MAP2-apposed boutons exhibited significantly stronger release than GFAP-associated sites (Fig. 2P – R). Collectively, these data demonstrate that dopamine release is tightly coupled to structural apposition with neuronal targets, exhibiting an all-or-nothing propensity in which active varicosities display high release probability while non-apposed quiescent sites remain refractory.

### Dopamine Axons Selectively Target Dopamine Receptor Expressing Neurons to form Release-Competent Appositions

Neurons in culture are highly synaptogenic, readily forming synapses with protein-coated beads or non-neuronal cells heterologously expressing postsynaptic molecules (*32–35*). We therefore asked whether the appositional contacts made by active dopamine axons occur stochastically with all neurons or are preferentially targeted toward processes expressing dopamine receptors. Unlike receptors for fast ionotropic ligands, which are broadly expressed, dopamine receptors are expressed in specific neuronal subtypes. Thus, the stereotyped and sparsely distributed axo-somatic associations (e.g., Fig. 2L) observed between release-competent dopaminergic axons and some cortical neurons in our DA:C co-cultures provided a suitable substrate to probe the specificity of appositional contacts formed by active DA varicosities.

To assess specificity, we first performed post hoc RNA fluorescence in situ hybridization (FISH) to profile dopamine receptor expression in DA:C co-cultures. Dopamine signaling is primarily mediated by D1 or D2-type receptors, encoded by Drd1 and Drd2 genes, respectively. Using a modified RNAScope protocol (see Methods), we probed Drd1/2 transcripts and found that dopamine axons preferentially associate with soma containing Drd1 or Drd2 mRNA (Fig. 3A–3C and Fig. S4). Specifically, we observed that dopamine axons exhibit a conspicuous tropism towards nuclei that were positive for Drd1 mRNA, and to a lesser extent, Drd2 mRNA (Fig. S4A – C). To show that this innervation pattern was not random, we segmented all DAPI-stained nuclei that were co-located in imaging fields containing well arborized dopamine axons, and measured distances between nuclei (approximating soma positions, see *Methods*) and the nearest dopaminergic axons. Distribution analysis of these distances revealed that Drd1^+^ nuclei were significantly closer to dopamine axons than is observed in the overall population (Fig. 3D). Nearly all of the Drd1^+^ DAPI signal (97%) were found within 2 µm of DA axonal processes vs. 56% for the overall population (Fig. 3D). Shuffling analysis confirmed this correlation was unlikely to arise randomly (Fig. 3D). Furthermore, quantifying tdTom intensity relative to distance from DAPI-stained nuclei showed that Drd1^+^ cells exhibited stronger axonal innervation than the overall population (Fig. 3E). Notably, axo-somatic innervations were more frequently observed at Drd1^+^ soma (16/27) compared with Drd2^+^-only soma (5/27), and Drd1^+^ nuclei received more intense axonal innervation than Drd2^+^-only nuclei. Together, these results indicate a clear bias in appositional contacts, with active varicosities targeting receptor-expressing cells, and further showed subtle distinctions in innervation pattern between Drd1-and Drd2-expressing cell populations.

**Fig. 3.**
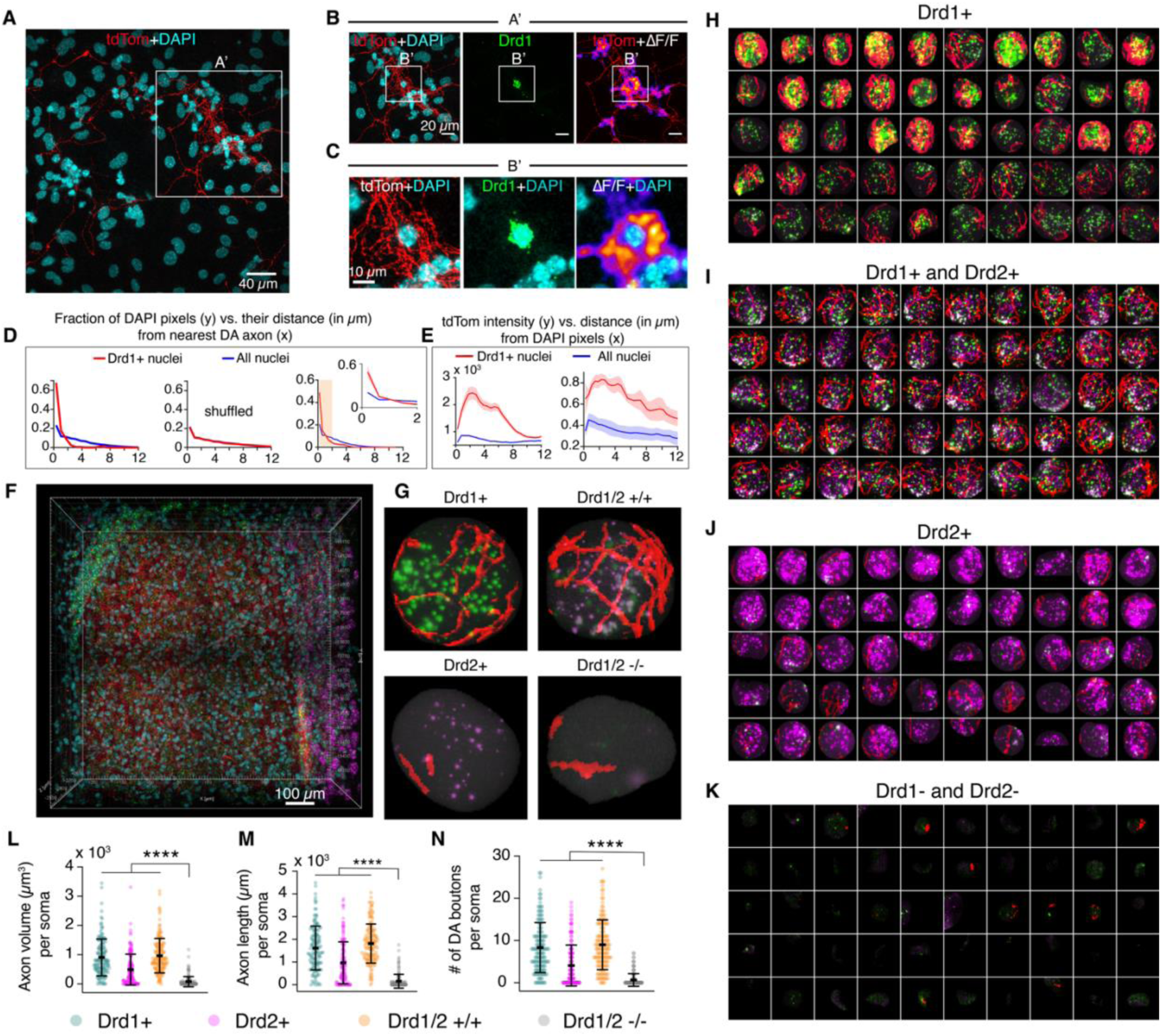
Dopamine axons target dopamine receptor expressing neurons to form release-competent appositional contacts. (A) A FOV showing dopaminergic axonal processes (red, tdTom) and DAPI stain (cyan). Note the convergence of dopamine axons towards the cell at the center of the ROI depicted in the solid white box (A’). (B) The ROI (A’) depicted in panel (A) is expanded and shown as a three-panel figure, including tdTom+DAPI, Drd1 signal from RNAScope, and tdTom+ΔF/F showing dopamine release. Note the tropism of tdTom processes (i.e. DA axons) towards the Drd1+ soma, and the robust dopamine secretion around the soma. (C) A close-up corresponding to the ROI (B’) depicted in (B), highlighting a “corona” of extracellular dopamine around the Drd1+ soma. (D) Left: Distance analysis shows that convergence of dopamine axons onto Drd1+ soma is unlikely to be due to chance. Fraction (i.e., percent of total) of DAPI+ pixels are plotted as a function of their distance from nearest tdTom pixel. DAPI pixels from Drd1+ cells (red) have a markedly different distribution compared to the overall distribution (blue). 97% of the Drd1+ DAPI signal is found within 2 µm of a tdTom process vs. 56% for the overall population (blue); p-value < 10^-4^ using two sample KS test. Middle: shuffling analysis shows the observed differences are not due to chance. tdTom pixels were randomly translated while DAPI pixels remained static, and the analysis was repeated over 100 such shuffling trials. The mean of the distributions from 100 trials shows no difference in the profile between Drd1+ cells and all cells. Right: proximity analysis averaged across n = 6 independent fields of view. Inset: first 2 µm on the x-axis. (E) tdTom intensity analysis shows that Drd1+ nuclei receive stronger innervation by dopamine axons. Left: data corresponding to FOV shown in (A). Right: Normalized data averaged from n = 6 independent FOVs. (F) A representative EASI-FISH volume in amygdala, showing dopamine axons (red), DAPI (blue), and Drd1 and Drd2 mRNA puncta (green and magenta, respectively). (G) DA axonal innervation of representative nuclei from four classes of neurons extracted from the EASI-FISH volume. (H–K) Machine learning-based classification of all nuclei in the EASI-FISH volume shown in (F) into Drd1+, Drd2+, Drd1/2 +/+, and Drd1/2-/-classes. Nuclei (n = 150) belonging to each class were identified, of which n = 50 are shown here. (L–N) Quantification of dopamine axon innervation of the four classes of nuclei (n = 150) is shown. Each point is a nucleus, which is dilated to approximate soma location (see Methods). Total volume and length of axonal innervation, and number of DA boutons were calculated for each nucleus. Statistical test of significance was performed using KS-test; p-value: ****<10^-4^, from pairwise comparison of Drd1/2-/-vs. each of the rest. Mean ± SD bars are shown.

We next asked whether dopaminergic axons can release transmitter when they contact somata lacking dopamine receptor transcripts. In fields where dopamine axons formed discrete axo-somatic “coronas” around individual neurons, RNAscope showed that dopamine release was restricted to somata expressing Drd1 and Drd2 transcripts (Fig. S4D – E). Within the same fields of view, nearby axons that contacted receptor-negative soma or processes showed no detectable release and remained quiescent (Fig. S4F). Thus, physical proximity or co-mingling of dopamine axons with neuronal membranes is not sufficient to confer release competence; instead, release competence emerges selectively at contacts onto receptor-expressing neuronal processes.

While transcript-level mapping defines the molecular identity of target neurons, it does not establish receptor functionality or membrane localization. To verify that dopamine receptors are not only transcribed but also present and active at the membrane, we directly measured receptor-mediated signaling in ‘postsynaptic’ partners. Fast synapses are defined by rapid ionic currents that confirm their functional coupling; by analogy, we asked whether appositionally-engaged active dopamine varicosities evoke a biochemical counterpart (that is, a receptor-dependent cAMP response) in their targets.

To obtain spatiotemporal coupling between dopamine release and cAMP modulation, we performed dual-color imaging of dopamine and cAMP using viral transduction of the intensity-based cAMP sensor G-Flamp1 (*36*). We then examined the degree of colocalization between dopamine release events and cAMP dynamics. We found that cAMP activity in ‘postsynaptic’ partners was spatiotemporally correlated with dopamine release from active varicosities (Fig. S5). We observed clear dopamine receptor–mediated increases in cAMP, with cAMP signals lagging dopamine release by 4.3 ± 2.3 s (mean ± SD) in Drd1+ neurons (Fig. S5C). In Drd2-expressing cortical neurons receiving axo-somatic dopamine input, we detected a temporally correlated decrease in cAMP across soma and proximal dendrites (Fig. S5D–G). Receptor identity determined by post hoc RNAScope was consistent with the direction of cAMP modulation (negative) measured by G-Flamp1 (Fig. S5E). While cAMP activity in soma and primary dendrites was largely synchronized, distal dendritic branches displayed localized hotspots of cAMP activity that aligned spatially with dopamine release sites visualized on DopaFilm (Fig. S5H–I).

We next asked whether receptor-specific targeting of DA axons is also observed in intact brain tissue. Because axo-somatic innervations were observed more frequently in DA:C co-cultures than in DA:S (72% vs 30%, respectively), we first examined mesocorticolimbic axons from VTA dopamine neurons projecting to the amygdala (including basolateral amygdala (BLA) and central amygdala (CeA)). We combined transcriptomic profiling (HCR) of dopamine receptor expression with expansion microscopy for high-resolution imaging (EASI-FISH, ref. (*37*)). EASI-FISH couples tissue expansion with multiplexed in situ hybridization to map gene-expression–defined cell types and their spatial relationships to axons at near-synaptic resolution in intact tissue, making it well suited to relate receptor transcript identity to local dopaminergic innervation (Fig. 3F). The amygdala provided robust VTA-derived dopaminergic innervation and a balanced representation of Drd1/2-positive and-negative neurons, enabling a sensitive test of receptor-associated wiring specificity in relatively sparse projections. In parallel, we extended this approach to the dorsal striatum, where dense, fine dopaminergic fibers arising largely from SNc neurons course through a neuropil rich in Drd1/2 expressing neurons. Together, these two anatomical regions, with moderately innervated amygdala with mixed receptor expression and densely innervated striatum with high Drd1/2 cell density, allowed us to assess receptor-specific targeting across distinct dopaminergic projection systems.

Using EASI-FISH, we segmented all DAPI-stained nuclei in amygdala and used machine learning to classify them into Drd1+, Drd2+, Drd1+ and Drd2+ double positive (Drd1/2 +/+), or receptor-negative (Drd1/2-/-) subgroups (Fig. 3G). Consistent with previous studies, a substantial fraction (73%, 1840 neurons out of 2521) of amygdala neurons expressed Drd1 and/or Drd2 (*38*). Soma expressing Drd1 alone (12%) or both Drd1 and Drd2 (13%) received the strongest axonal innervation, with dopamine axons forming pericellular basket innervations around such soma (Fig. 3H–I). These innervations were less intense in Drd2 neurons (Fig. 3J) and, critically, were not observed in neurons lacking Drd1 and Drd2 (27%, Fig. 3K). We quantified these observations by measuring total axonal volume per soma, contour length of soma-associated axons, and ramification (number of boutons). Across all metrics, Drd1 and Drd2-expressing cells were more heavily innervated than receptor-negative neurons (Fig. 3L–N). Specifically, total axonal volume was highest around Drd1+ soma (905 ± 51 µm³) and Drd1/2 (+/+) soma (964 ± 48 µm³), remained robust but reduced around Drd2+ soma (490 ± 43 µm³), and was sharply lower around Drd1/2 (−/−) soma (78 ± 14 µm³) (all mean ± SEM). Similarly, total length of soma-associated axons followed the same pattern, with 1616 ± 79 µm around Drd1+ soma, 1814 ± 70 µm around Drd1/2 (+/+) soma, 964 ± 75 µm around Drd2+ soma, and 154 ± 25 µm around Drd1/2 (−/−) soma (all mean ± SEM). We also segmented axonal processes to identify boutons: Drd1+ and Drd1/2 (+/+) soma exhibited more boutons (8.3 ± 0.48 and 9.0 ± 0.48, respectively), compared to 4.1 ± 0.40 around Drd2+ soma and 0.7 ± 0.12 around Drd1/2 (−/−) soma (all mean ± SEM). Consistent with our observations in culture, the density of innervation was markedly lower in Drd2+ soma compared to Drd1+ soma (Fig. 3H–I vs. 3J, Fig. 3L–N), confirming the differences in innervation between dopamine receptor subtypes observed in vitro.

Extending this analysis to the dorsal striatum, we applied EASI-FISH to segment all DAPI-stained nuclei and similarly classifed neurons into Drd1+, Drd2+, Drd1/2 (+/+), or receptor-negative (Drd1/2 −/−) subgroups (Fig. S6). In this region, dopamine axons formed pervasive fine-caliber arborizations throughout the neuropil, yet receptor specificity was still preserved: soma classified as Drd1+ or Drd2+ showed the strongest soma-associated innervation, whereas receptor-negative neurons exhibited markedly reduced soma-associated axonal coverage (Fig. S6). Quantification of perisomatic axonal volume, contour length, and bouton number confirmed significantly higher innervation of Drd1/2-expressing soma relative to Drd1/2 (−/−) cells (Fig. S6), indicating that receptor-driven tropism of dopaminergic axons generalizes from the sparser VTA-to-amygdala projection to the dense SNc-to-striatal projection system. Notably, when comparing EASI-FISH datasets across the two regions, two contrasts stood out. First, unlike the amygdala, Drd1/2 double-positive soma were rarely observed in the striatum (Fig. 3L–3N vs. Fig. S6). Second, whereas Drd2+ cells in the amygdala were less robustly innervated than Drd1+ cells, Drd2+ cells in the striatum received innervation comparable to Drd1+ cells. In summary, dopamine axons exhibit strong tropism toward receptor-expressing neurons with which they form appositional contacts. This receptor-dependent innervation is evident in both dissociated cultures and intact tissue, and is tightly linked to release competence in vitro. These findings suggest that dopaminergic varicosities align their output with target neurons whose transcriptomic profiles support dopamine receptor expression.

### Visualizing Receptor Nanoscale Organization with ALFADoR Mice

While transcriptomic profiling has revealed axonal tropism and wiring specificity in dopamine axonal outgrowth, and cAMP activity imaging suggest the presence of compartmentalized signaling at the subcellular level, it does not directly address the spatial relationship between dopamine receptors and dopamine release sites. To resolve this, it is necessary to map dopamine receptor distribution with subcellular resolution. However, the subcellular distribution of dopamine receptors, which are G-protein coupled, is poorly characterized, partly due to the lack of reliable reagents compatible with immunolabeling, a known and vexing issue for the field (*39*). To overcome this limitation, we focused on the two predominant dopamine receptors, D1 and D2, and generated transgenic mice in which one copy of the ALFA-tag (*40*) was knocked into the N-terminus of either receptor using CRISPR-mediated gene editing (Fig. 4A) (Methods). We confirmed precise ALFA-tag insertion at the target gene loci by PCR genotyping and Sanger sequencing. Knock-in animals showed the expected 5′ and 3′ junction amplicon sizes (wild-type versus knock-in fragments), confirming correct homology-directed integration (Fig. S7 and Fig. S8). Sanger chromatograms spanning both junctions showed single, high-quality reads with good peak resolution and complete continuity across the genome-to-ALFA and ALFA-to-genome boundaries on both ends (Fig. S7 and Fig. S8). Alignment of the reads to the reference sequence showed an in-frame fusion without insertions, deletions, or other sequence deviations, verifying an error-free ALFA-tag knock-in at the intended locus. In vitro, mRNA for each gene corresponded with ALFA-tag positive cells, confirming specificity (Fig. S7 and Fig. S8). We refer to these animals as ALFADoR mice (<u>ALFA</u>-tagged <u>do</u>pamine receptor mice; ALFADoR1 for Drd1 KI and ALFADoR2 for Drd2 KI).

**Fig. 4.**
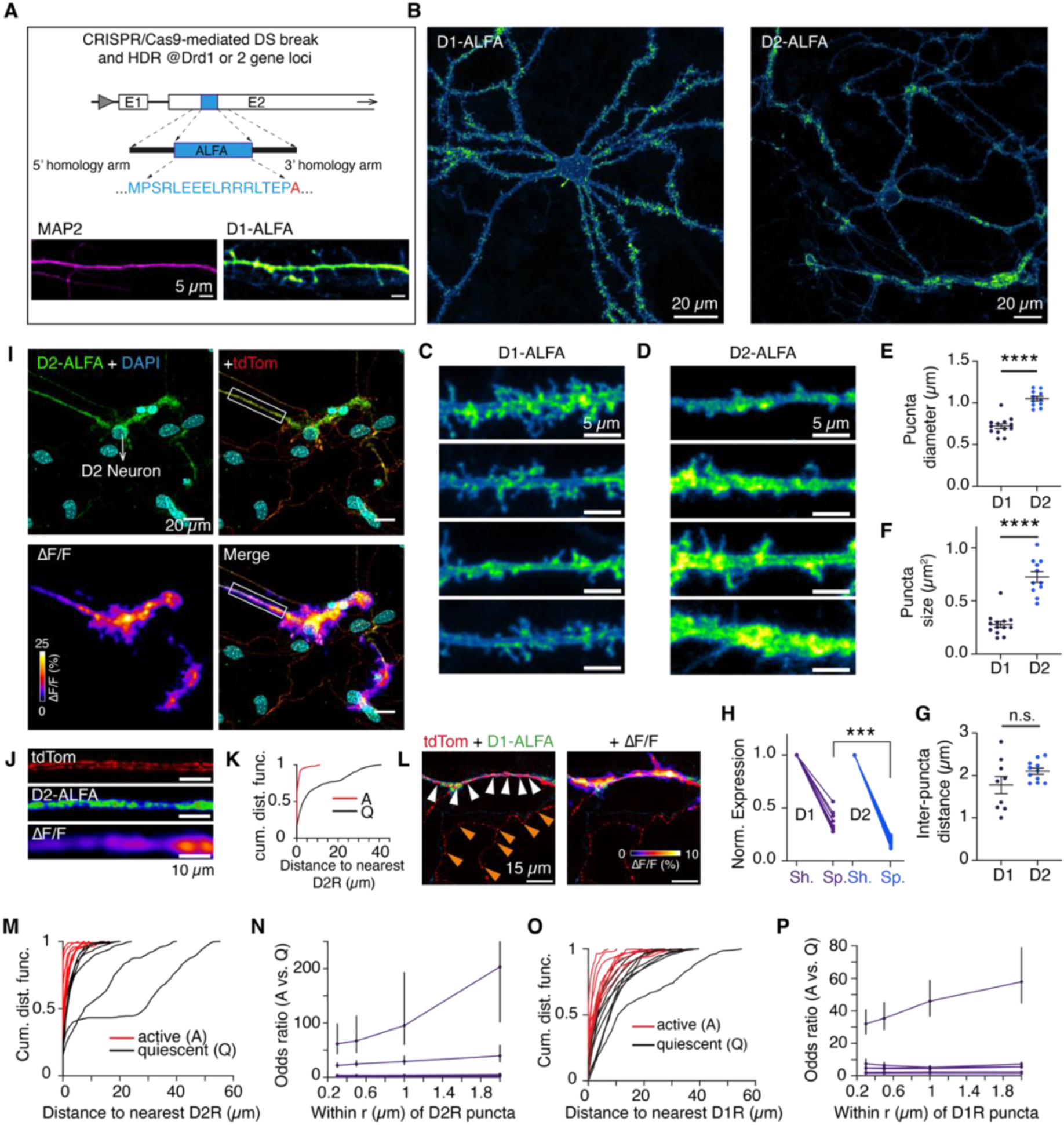
The subcellular distribution of D1 and D2 dopamine receptors, visualized with ALFADoR KI mice. (A) Design of ALFADoR mice. Bottom panel: MAP2+ dendritic process of a Drd1+ neuron, and its corresponding D1-ALFA readout, showing receptor enrichment in shaft and spines. (B) Pan-neuronal live cell visualization of D1-ALFA and D2-ALFA expression with subcellular resolution using anti-ALFA-tag nanobodies conjugated to Atto643. (C–D) Representative expression of D1-ALFA and D2-ALFA in four dendritic segments. (E–F) D1-ALFA and D2-ALFA puncta size (area, characteristic linear scale reported as ‘diameter’). Each point represents mean puncta size averaged over one neuron, and mean ± SEM bars are shown. Significance was evaluated with a nonparametric two-sample KS test; p-value: **** < 10^-4^ (G) Interpuncta distance of D1-and D2-ALFA signals. Each point represents mean value from one neuron, and mean ± SEM bars are shown. Significance was evaluated with a nonparametric two-sample KS test; not significant (n.s.). (H) Comparison of expression intensity in shafts (Sh.) vs. spines (Sp.), normalized against values in shaft. Each point represents average of puncta over a dendritic segment, and spine averages are normalized against shaft average. Note spine intensity of D2-ALFA expression is low relative to D1-ALFA (38% of shaft intensity vs. 18.5% of shaft intensity, p-value: *** < 10^-3^) on two-sample KS test). (I) DA axons (tdTom) arborizing in a FOV containing a D2-ALFA-positive neuron. Dopamine ΔF/F closely tracks D2-ALFA expression pattern, with only receptor-apposed axons releasing dopamine. (J) Close-up of the ROI (white box) depicted in (I), showing apposition of active varicosities and D2-ALFA puncta. Scale bar is 10µm. (K) Analysis of nearest-receptor distance distribution of active (A) and quiescent (Q) varicosities corresponding to (I), showing, as a population, active varicosities are found much closer to D2-ALFA puncta than quiescent ones. (L) Dopamine release in D1-ALFA apposed axons (tdTom, red) shows D1-puncta lying in close apposition to active varicosities (white arrow heads) vs. quiescent varicosities (orange arrow heads). (M–P) Analysis of distances between active (A) and quiescent (Q) varicosities and receptor puncta. Cumulative distribution of nearest-puncta to A and Q varicosities is shown in (M) and (O). There are n = 7 and n = 9 paired traces (A&Q) from biological replicates for D2 and D1, respectivey. (N, P) odds ratio: calculated as the odds of finding A or Q varicosities withing ‘r’ µm radius of a D2 or D1 receptor puncta; y-values >1 indicate enrichment of active varicosities closer to receptor puncta; all traces lie above the y = 1 line; vertical bar represents a 95% bootstrap confidence interval for the odds ratio at that radius (see Methods for more).

Using ALFADoR mice, we visualized the subcellular distribution of dopamine receptors in cortical and striatal cultures. In both cultures, dopamine receptors exhibited a punctate distribution (Fig. 4B–4D). D1 receptors formed puncta with mean characteristic length scales of 0.72 ± 0.03 µm and average area of 0.28 ± 0.03 µm^2^, whereas D2 receptor puncta were larger, averaging 1.05 ± 0.03 µm in linear scale and 0.72 ± 0.05 µm^2^ in area (mean ± SEM for all). (Fig. 4E – F). The mean inter-puncta distance was 1.78 ± 0.2 µm for D1, and 2.1 ± 0.07 µm for D2 (Fig. 4G). Dopamine receptor puncta were observed in both spiny neurons (predominantly striatal) and aspiny neuron types (predominantly cortical) (Fig. S9A–B). D1 puncta were observed in both dendritic shafts and dendritic spines, with puncta intensity in spines being approximately 38±3% (mean ± SEM) of those in shafts (Fig. 4H). In contrast, D2 puncta were primarily localized in dendritic shafts, with intensity of D2 expression in spines being approximately 18.5± 1.1% (mean ± SEM) of those in shafts, markedly lower compared to D1 intensity in spines (Fig. 4H). Importantly, we were able to visualize dopamine receptors in live-cell imaging using fluorescent anti-ALFA-tag nanobodies, confirming the N-terminus locus and extracellular accessibility of the tag (Fig. S9C–D) and performed cAMP imaging to confirm receptor functionality (Fig. S9E–F). Collectively, these data indicate that dopamine receptors exhibit a structured and heterogenous subcellular distribution rather than uniform membrane dispersion. This patterning suggests that GPCR-mediated neuromodulation may be organized at a finer subcellular scale than previously recognized.

With this context, we examined how release competent (active) varicosities are organized relative to dopamine receptor puncta. Here, DA neurons were cultured from the standard *Slc6a3*-Cre x Ai9 line and the striatal and cortical co-cultures were produced from ALFADoR1 or ALFADoR2 mice. In mixed co-cultures, and consistent with the tropism observed with RNA FISH data, we found that dopamine axons preferentially innervate dopamine receptor expressing processes, forming close appositional contact between active varicosities and receptor puncta (Fig. 4I–4L, Fig. S10).

To quantify these associations, we calculated the distance between all dopamine axonal varicosities (active and quiescent) and the nearest D1 or D2 receptor puncta. We performed two analyses on these distances: (1) the cumulative distribution function (CDF) of the distances between varicosities and receptor puncta, and (2) the odds ratio (OR) of finding an active or quiescent varicosity within the vicinity of dopamine receptor puncta (an OR > 1 indicates enrichment of active varicosities near receptor puncta relative to quiescent ones). Using the first metric, we found that active varicosities, as a population, lie closer to dopamine receptor puncta than inactive processes (Fig. 4M, 4O). For example, the CDF showed that 75±6% of all active varicosities were within 1 µm of a D2 receptor punctum, compared with 41±4% of quiescent varicosities (Kolmogorov–Smirnov test p-value = 0.0006; mean±SEM, Fig. 4M). Similarly, 45±6% of all active varicosites were within 1 µm of a D1 receptor punctum, compared with 20±4% of quiescent varicosities (Kolmogorov–Smirnov test p-value = 0.034; mean± SEM, Fig. 4O).

Using the second metric, we similarly found that the odds ratio of finding an active varicosity within ‘x’ (x = 0.2, 0.3, 1, 2 µm) radius of a receptor puncta was well above 1 (Fig. 4N, 4P); (@ x=1 µm, OR = 7.85±4.8 for D1-ALFA and 15.7±10.4; mean ± SEM for both). These evaluations demonstrate a spatial alignment between dopamine release sites and receptor-rich domains that cannot be accounted for by chance alone. Integrating these proximity data with RNA-FISH mapping of receptor expression, and EASI-FISH analyses in the amygdala and striatum, our results indicate that dopamine axons are biased toward receptor-expressing targets, and release sites are anatomically apposed to receptor clusters.

### Dopamine Releasing Varicosities are Enriched with Specializations that Facilitate Temporally Precise Neurotransmitter Release

Given the proximity between dopamine release sites and receptor clusters, we next asked whether release-competent varicosities possess molecular specializations characteristic of classical presynaptic terminals. Using immunofluorescence microscopy, we compared active and quiescent varicosities to determine whether active sites exhibit a molecular architecture consistent with synaptic organization. Synaptophysin (SYP) staining revealed that appositional contacts formed by active varicosities were enriched in SYP, consistent with vesicle clustering at the interface and suggesting the presence of a synapse-like presynaptic specialization at these sites (Fig. 5A–B). Encouraged by this observation, and by previous reports showing that dopamine release sites contain cytomatrix proteins found in fast synapses (*27, 41*), we examined additional presynaptic proteins.

**Fig. 5.**
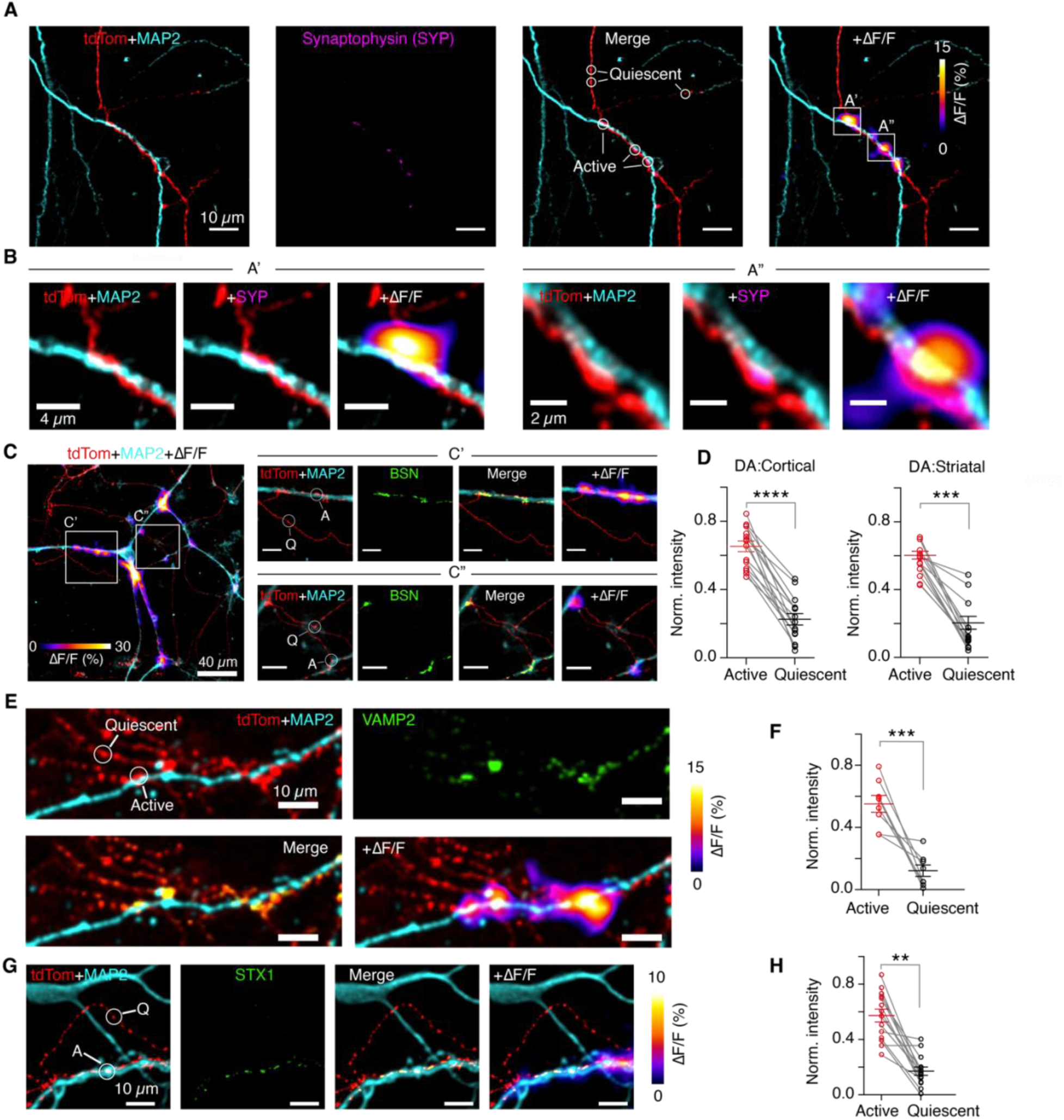
Assessment of synaptic release machinery at active vs. quiescent varicosities. (A) Dopamine axonal process (tdTom) innervating a dendrite (MAP2), synaptophysin (SYP) readout from the same field of view (SYP), overlaid with dopamine release activity measured on DopaFilm (+ΔF/F). (B) Close-up of ROI’s A’ and A” showing enrichment in synaptophysin (SYP, magenta) at the interface between the red (tdTom, DA axon) and cyan (MAP2, dendrite) channels. (C) Dopamine neuron axonal processes (red, tdTom) innervating a MAP2 immunostained cortical neuron (cyan), overlaid with dopamine release activity measured on DopaFilm (ΔF/F). Right: the ROIs shown in white boxes (C’ and C”) are expanded, bassoon (Bsn) expression in shown in green, and overlaid with dopamine release (+ΔF/F). Representative active (A) and quiescent (Q) varicosities are depicted. (D) Bsn expression assayed at active and quiescent varicosities. Each point is mean from a biological replicate, averaging Bsn expression at 10 randomly selected active and quiescent varicosities from the same sample. Data is normalized against the highest intensity varicosity, and averaged; mean ± SEM bars are shown. Significance was evaluated with a nonparametric two-sample KS test; p-values: **** <10^-4^, *** <10^-3^. (E) Dopamine release from an axonal arbor innervating a dendritic process (cyan, MAP2), and synaptobrevin 2 immunofluorescence readout (Vamp2, green). Representative active and quiescent varicosities are depicted. (F) Syaptobrevin 2 (Vamp2) expression at active vs. quiescent varicosities. Each point is mean from a biological replicate, averaging Vamp2 expression at 10 randomly selected active and quiescent varicosities from the same sample. Data is normalized against the highest intensity varicosity, and averaged; mean ± SEM bars are shown. Significance was evaluated with a nonparametric two-sample KS test; p-values: *** < 10^-3^. (G) Dopamine release from an axonal process innervating a dendritic processes (MAP2), and syntaxin 1 immunofluorescence readout (Stx1, green). Representative active (A) and quiescent (Q) varicosities are depicted. (H) Stx1 expression at active vs. quiescent varicosities. Each point is mean from a biological replicate, averaging Stx1 expression at 10 randomly selected active and quiescent varicosities from the same sample. Significance was evaluated with a nonparametric two-sample KS test; p-value: ** < 10^-2^.

At conventional synapses, the scaffolding protein bassoon (Bsn) is concentrated at the active zone and is among the first molecules recruited during presynaptic assembly (*42, 43*). Using immunofluorescence microscopy, we found that Bsn expression strongly correlated with release competency in dopamine varicosities (Fig. 5C). Bsn levels were significantly higher in active compared to quiescent varicosities across both DA:C (0.65±0.03 vs. 0.22±0.03, mean± SEM on normalized data) and DA:S co-cultures (0.57±0.02 vs. 0.17 ±0.04, mean± SEM on normalized data) (Fig. 5D), and Bsn expression correlated with dopamine release amplitude (ΔF) (Fig. S11A–B). The correspondence between DopaFilm hotspots and Bsn enriched varicosities was precise, especially at non-branching axons (Fig. 5C). At branch points or where dopamine axons innervate dendritic shafts, varicosities were ramified and produced merged hotspots in DopaFilm recordings (compare Fig. S11C&D vs Fig. S11C&E). Although individual hotspots were not visually resolvable in such clusters, non-negative matrix factorization (NNMF) analysis (see Methods) successfully separated overlapping signals, and the computationally resolved hotspots colocalized with individual or clustered Bsn puncta in immunofluorescence images (Fig. S11E).

Other presynaptic proteins essential for exocytosis were similarly enriched at active varicosities. The v-SNARE protein synaptobrevin 2 (Vamp2) and the t-SNARE Syntaxin 1 (Stx1) were both enriched relative to quiescent varicosities 0.55±0.05 vs. 0.12± 0.04, mean± SEM on normalized data) (Fig. 5E–H). The vesicle-priming protein Munc13-1 showed preferential enrichment in active varicosities (Fig. S11F–G). Rims1/2 (RIM), a scaffolding protein previously shown to be critical for dopamine release (*27, 44*), was likewise enriched at active sites (Fig. S11H–I).

To test whether these molecular specializations are required for dopamine release, we performed RNA interference-mediated knockdown (KD) of Bsn, RIM, Stx1, Syt1, Vamp2, and Munc13-1 in our co-culture assays (Fig. S12A–B). Using quantitative proteomics, we confirmed efficient depletion of each protein target (Fig S12 C–D). Munc13-1 and RIM KD efficiencies were the highest, showing near complete depletion, whereas the remaining targets were depleted with efficiencies ranging from 68% (Bsn) to 98% (Vamp2) (Figure S12C–D). In each KD preparation, we measured evoked dopamine release, and showed that it was nearly completely abolished for Munc13-1, RIM, Stx1 and Syt1 knock down (Fig. S12E–F). Bsn KD reduced the extent of dopamine release but did not completely eliminate it (Fig. S12F). In contrast, we did not observe any diminution in evoked dopamine release in cells transduced with scrambled control shRNA ( Fig. S12F).

Taken together, these findings show that release-competent dopamine varicosities harbor a canonical presynaptic molecular architecture. Their enrichment in Bassoon, Munc13-1, RIM, Vamp2, Stx1, and Syt1 mirrors that of fast chemical synapses, and targeted disruption of these proteins functionally silences dopamine release. Together with receptor organization and FISH-based axonal tropism, these results reveal that dopamine axons possess synapse-like structural and molecular specializations and wiring specificity that confer precision and efficiency to dopamine signaling.

### Ultrastructural Assessment of Dopamine Releasing Varicosities in Contact with Dendrites and Soma

To further define the structural basis of dopamine release, we examined the ultrastructure of release-competent (active) varicosities that formed appositional contacts with neuronal targets. Because dopamine neurons express Cre recombinase in our co-culture system (*Slc6a3*-Cre x Ai9), we used a cytosol-localized pAAV–EF1a-DIO-APEX2.NES construct (*45*) to label dopaminergic varicosities and correlate their ultrastructure with ΔF/F activity (Fig. 6A). Viral expression levels were titrated such that dopaminergic processes exhibited enhanced electron density under transmission electron microscopy (TEM) while preserving ultrastructural detail (Fig. S13A). Co-cultures were grown on gridded DopaFilm substrates, enabling us to identify the same imaging fields in live fluorescence and electron microscopy sessions (Fig. 6A). Following live cell imaging, cultures were fixed and processed for serial-section TEM (section thickness = 50 nm). Low-magnification electron micrographs were first used to align light and TEM datasets, and DAB reaction product provided positive identification of dopaminergic (DAB+) varicosities.

**Fig. 6.**
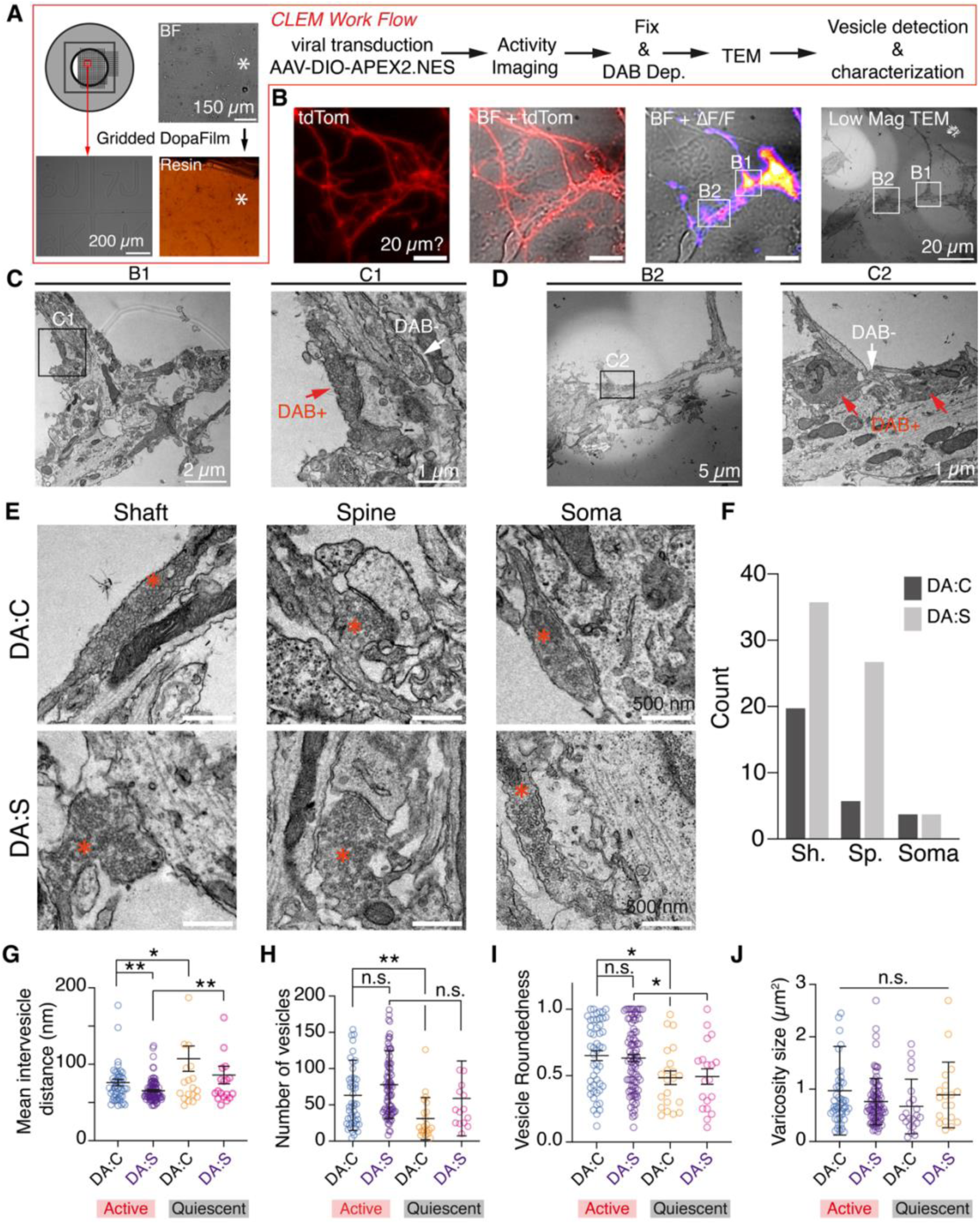
Activity-guided ultrastructural characterization of dopamine release sites near dendrites and soma of target neurons. (A) Schematic of a gridded (alphanumerically patterned) cell culture dish where neurons are sparsely co-cultured on DopaFilm. After dopamine release sites are recorded in live cell imaging, the specimen is fixed and processed for EM (Resin) (see Methods). The ROI is identified in the resin and trimmed. Thin serial sections collected from the resin blocks are imaged using TEM. To identify DA varicosities, virally transduced APEX2-based DAB deposition is employed. (B) Dopaminergic axonal processes innervating a dendrite of a cortical neuron shown in brightfield (BF). Dopamine ΔF/F is from live-cell imaging on DopaFilm, and the corresponding low magnification (low mag) TEM of the same FOV is shown in the right-most panel. (C) Left: Close-up of ROI B1 depicted in (B). Right: Further zoomed into ROI C1 depicted on the left panel, showing DAB+ varicosity lying in apposition against a dendritic process. DAB-varicosity is seen in the same field. (D) Left: Close-up of ROI B2 depicted in (B). Right: Further zoomed into ROI C2 depicted on the left panel, showing two DAB+ DA varicosities lying in apposition against a major dendrite. DAB-varicosity is seen in the same field. (E) Representative ultrastructures of DAB+ varicosities in DA:C and DA:S co-cultures, forming appositional contact with dendritic shafts, spines, and soma. (F) Number of appositional contacts of each category visualized in survey of TEM thin sections. (G–J) Comparison of active and quiescent varicosities (both DAB+). Each point represents average statistic over a varicosity. Mean ± SEM bars are shown in G, H, I and mean ± SD bars are shown in J. Significance was evaluated with a nonparametric two-sample KS test; p-values: * < 0.05, ** < 10^-2^, n.s. = not significant.

We first surveyed dopaminergic and non-dopaminergic boutons within the same imaging fields and performed morphometric analysis using a previously published method (Fig. S13B) (*46*). Dopaminergic varicosities contained numerous small, clear vesicles averaging 0.14 µm² per varicosity, whereas vesicles in non-dopaminergic boutons averaged 0.07 µm². Dopamine varicosities harbored more vesicles compared to non-dopaminergic boutons (90 vs. 48 per bouton, n=15) while retaining similar inter-vesicle distances (56 nm vs. 56 nm per bouton, n =15), indicative of similar vesicle clustering (Fig. S13C). Consistent with this observation, vesicles occupied 20% of total varicosity volume in dopaminergic boutons compared with 17% in non-dopaminergic boutons (Fig. S13C). Classical synaptic ultrastructure defined by parallel pre-and postsynaptic membranes separated by a uniform cleft were observed in 60% of non-dopaminergic varicosities but only 40% of dopamine varicosities. Instead, most dopamine varicosities formed tight appositional contacts with their targets, characterized by narrow, undulating interfaces lacking a well-defined cleft (Fig. S13B). Using correlated light and electron microscopy (CLEM), we confirmed that DAB+ varicosities co-localized with ΔF/F release hotspots, linking this morphology directly to functional activity (Fig. 6B–D). These observations indicate that dopamine varicosities form structural appositions that somewhat deviate from canonical synapses but retain key synaptic hallmarks, including vesicle clustering, membrane apposition, and release competence.

We next characterized the appositional partners of dopaminergic varicosities across DA:C and DA:S co-cultures (Fig. 6E). Survey of 58 serial sections from DA:C and 68 sections from DA:S co-cultures revealed that spine appositions were more frequent in DA:S than in DA:C (Fig. 6F). Notably, active dopaminergic boutons contacting dendritic spines exhibited classical synaptic ultrastructure in 56% of cases, compared with 47% for shaft and somatic contacts, suggesting compartment-specific organization of dopamine signaling may be present. Finally, comparison of active and quiescent varicosities revealed ultrastructural differences: quiescent boutons contained fewer, more loosely packed vesicles and showed greater vesicle pleomorphism (Fig. 6G–J), although their overall varicosity size was similar. Taken together, these findings demonstrate that dopamine varicosities possess ultrastructural organization that closely parallels but does not fully resemble classical synapses.

### Spatial organization of dopamine receptor microdomains in intact brain tissue

Based on the evidence that dopamine axons contain presynaptic specializations consistent with synapse-like release sites, and that dopamine axons exhibit a tropism for receptor expressing processes, we next asked whether endogenous dopamine receptors are organized in intact tissue in a manner that enables spatially restricted, local neuromodulation. To do so, we focused on three predominant terminal fields of midbrain dopamine neurons, namely, the striatum (Str), amygdala (Amg), and prefrontal cortex (PFC; abbreviated here as Ctx), which together span dense and sparse innervation regimes and distinct dopamine neuronal subtypes (*47–49*). At the macroscopic level, conventional, non-expanded immunohistochemistry against the AFLA-tag confirms that both D1 and D2 receptors are strongly enriched in the striatum (Fig. 7A), consistent with the high density of of D1 and D2 expressing neurons in Str. However, this spatial scale does not address how receptor signaling machinery is spatially organized relative to dopamine release sites.

**Fig. 7.**
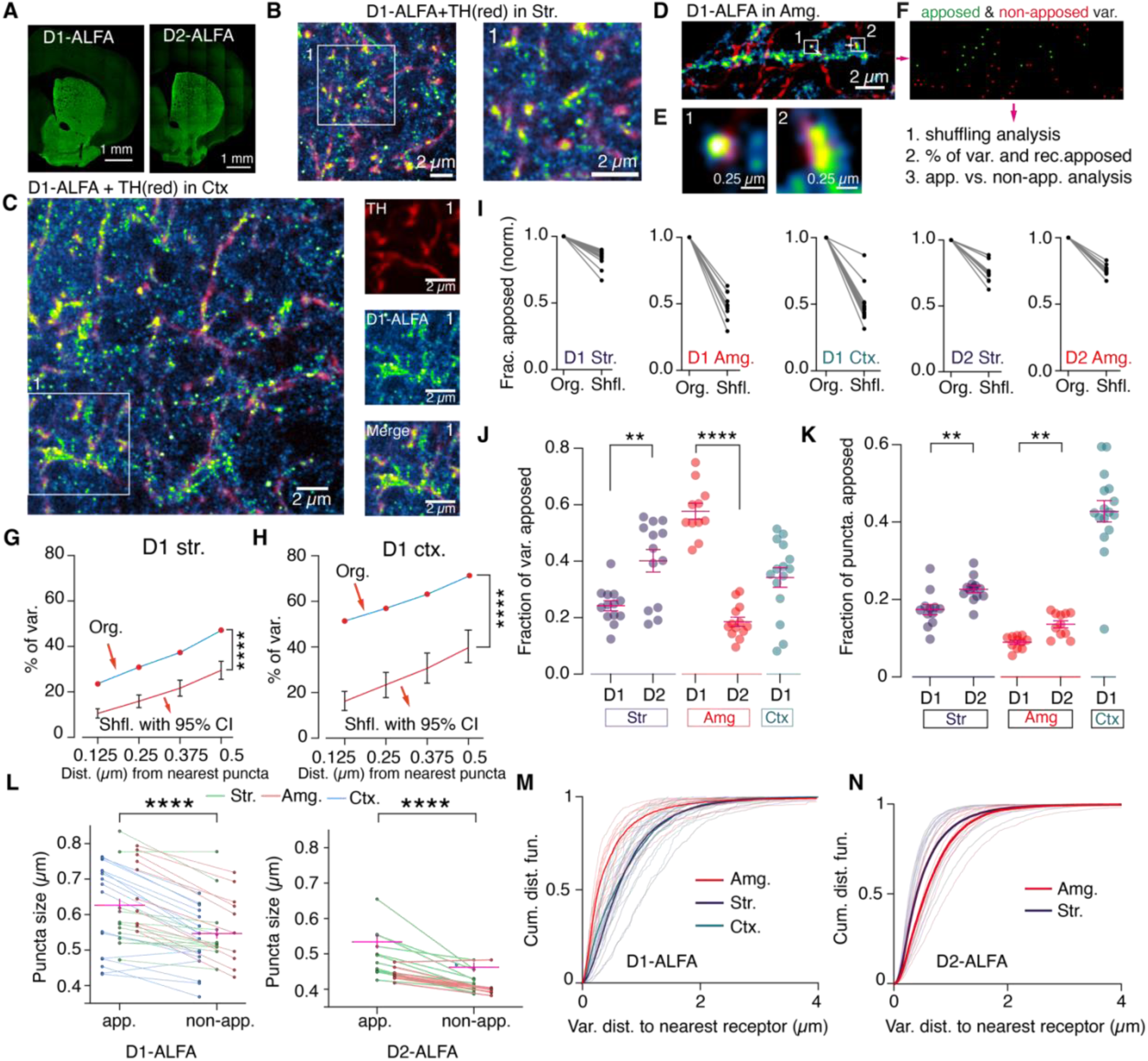
Visualizing the spatial organization of dopamine receptor microdomains in intact brain tissue using expansion microscopy. (A) Images of immunostained 50µm coronal tissue slices from ALFADoR mice, showing the expected enrichment of D1 and D2 receptors in the striatum. (B) Two-color composite image of D1-ALFA and dopamine axons (TH, red) in the striatum in 4x expanded tissue. Right panel: close-up of the ROI-1 (white box) depicted in the left panel. Scale bars represents real biological scale. (C) ExM images (4x) of D1-ALFA and tyrosine hydroxylase (TH) expressing axons in the pre-frontal cortex. Right panels: close up of ROI-1 (white box) depicted on the left. (D) Expression in the basolateral amygdala (ExM, 4x), with TH (red) representing dopaminergic varicosities. (E) Close up of dopaminergic varicosities (red) in tight appositional juxtaposition with D1-ALFA clusters, corresponding to ROIs-1 and-2 depicted in (D). (F) Varicosities are classified as apposed or non-apposed based using proximity analysis (see Methods). Subsequently, toroidal shuffling null models and nearest distance analysis are used to quantify ExM volumes. (G–H) Toroidal shuffling of ExM data in D1-ALFA in striatum (str.) and prefrontal cortex (ctx.). First, the percentage of varicosities that lie within 0.125, 0.25, 0.375, and 0.5 µm are evaluated from the original data (Org., blue trace). The analysis was then repeated after shuffling receptor puncta randomly (but uniformly) in the x-y plane n = 200 times, from which mean and 95% confidence interval (CI) ranges are evaluated (red trace); p-values reflect a one-sided permutation test against 200 shuffle runs. (I) Shuffling analysis performed as described in G–H for three anatomical regions and the two receptor subtypes. Data points corresponding to the 0.125 µm appositional contact are shown where each shuffled point (Shfl.) is normalized against the experimental data (Org.). (J–K) Fraction of apposed varicosities and apposed puncta are evaluated, and mean ± SEM bars are shown. Pair-wise comparisons were made to test significance of observed differences; significance was evaluated with a nonparametric two-sample KS test; p-values: ** < 10^-2^, **** < 10^-4^. (L) Comparison of apposed (app.) and non-apposed (non-app.) receptor puncta size show significant differences (KS-test, p-value: **** <10^-4^, unpaired t-test on pooled data). Magenta bars represent mean ± SEM of pooled data, and individual data points corresponding to the three anatomical regions are broken out for better visualization of anatomical differences. Each point represents mean puncta size averaged over a biological replicate. (M–N) Cumulative distribution functions of varicosity-to-nearest receptor puncta in three anatomical regions and two receptor subtypes.

To analyze receptor organization at the scale relevant to putative release–receptor coupling implied by our earlier findings, we used ALFADoR mice and performed expansion microscopy (ExM) in Str, Ctx, and Amg (*50*). In expanded tissue (expansion factor = 4), ALFA-tagged receptors appear as discrete puncta rather than diffuse membrane labeling, while dopaminergic axons visualized with a TH-stain (red) formed an arbor with typical varicosities (Fig. 7B–7D). Qualitatively, these images frequently reveal dopamine varicosities positioned immediately adjacent to receptor puncta, including examples in the striatum (right panel in Fig. 7C) and amygdala (Fig. 7D–7E) where close apposition is particularly clear in magnified views. This suggested that the punctate receptor architecture and appositional coupling may provide a structural substrate for receptor microdomains that could support locally constrained (i.e., synapse-like) dopamine signaling.

To quantify our ExM data, we implemented an analysis framework (schematized in Fig. 7F) that systematically extracts varicosities and receptor puncta from single or multi-plane ExM images and quantified their spatial relationships. Briefly, TH axons were segmented and varicosities were identified using an ExM-optimized, watershed-based approach (Methods); and receptor puncta were segmented as discrete clusters, again using watershed-based algorithms.

We then quantified spatial coupling by evaluating (i) minimum distance from each varicosity to the nearest receptor cluster, (ii) cumulative distribution functions (CDFs) of varicosity-to-receptor, and receptor-to-varicosity distances, and (iii) analysis of receptor puncta stratified by whether they are classified as apposed versus non-apposed. Critically, we compared the observed spatial coupling against a conservative null model using toroidal shuffling in which receptor masks are shifted randomly in (x,y) with wrap-around, preserving receptor number, size, and global density structure while disrupting their putative alignment relative to dopamine axons (see Methods for more). This provided anatomical region-specific “chance apposition” baseline without artificially redistributing puncta or changing their shapes.

Across all three regions, our data show that varicosity to receptor proximity cannot be explained by receptor density/chance-contact alone. In Str and Ctx (D1-ALFA), the fraction of varicosities within each of four increasingly permissive distance thresholds is substantially higher in the experimental data than in shuffled controls (Fig. 7G–H), with highly significant differences (p < 10⁻⁴ across thresholds, comparing mean of experimental data vs. Mean +/-95% CI evaluated from n = 200 shuffling trials). A complementary ‘normalized’ view (Fig. 7I) shows that in all three anatomical regions, shuffling consistently reduces apparent coupling across all tested conditions (D1 Str/Amg/Ctx and D2 Str/Amg, D2 in Ctx was not included in our analysis because of low D2 expression), with particularly dramatic reductions for D1 coupling in Amg and Ctx, indicating strong spatially-coupled organization in those regions. Specifically, shuffling reduced appositional contact to 49.9±0.02% of that observed in non-shuffled data (Org.) in D1-Amg and to 48.2±0.03% in D1-Ctx, but only reduced it to 83.6±0.02% in D1-Str (mean±SEM for all).

When we summarized coupling by receptor subtype and region (Fig. 7J–K), we observed two complementary trends that help clarify the architecture of dopamine signaling microdomains. First, from a varicosity-centric perspective (Fig. 7J), coupling was observed to be both region and receptor-dependent: in Str for example, D2-apposed varicosities exceed D1 (40.1±4% vs. 24± 1.8%, mean±SEM, p-value < 10^-2^), whereas in Amg the pattern reverses, with D1-apposed varicosities being strikingly higher than D2 (57.7±2.8% vs.18.6±1.6%, mean±SEM, p-value <10^-4^). In Ctx, where only D1 expression was analyzed, we observed an intermediate level of coupling compared with that observed in Str. and Amg (34.2±3.4%, mean±SEM). Second, from a receptor-centric perspective (Fig. 7K), the pattern differs in a way that is informative. Str and Amg show modest but significantly higher fractions of receptor puncta that are apposed for D2 relative to D1 (22.6 ±0.9% vs. 17.4±1.3% in Str., and 13.6±0.9% vs. 8.9±0.5% in Amg, mean±SEM), whereas in Ctx, D1 exhibits a relatively high fraction of puncta apposed (42.8±2.8%, mean±SEM). Considering the relatively low percentage of receptor puncta that are appositionally engaged (Fig. 7K), we asked whether apposed receptor clusters differ structurally from non-apposed clusters. Across all three regions, receptor puncta classified as apposed were systematically larger than non-apposed puncta, for both D1-ALFA and D2-ALFA (p-value < 10^-4^) (Fig. 7L). This result suggested that apposition is not simply a geometric coincidence, but is associated with distinct receptor microdomain properties and is consistent with the idea that apposed clusters may represent functionally specialized sites with enhanced receptor content that elicit local signaling.

Finally, we performed varicosity-to-receptor and receptor-to-varicosity distance distribution analysis. Cumulative distribution function (CDF) analyses of varicosity-to-nearest-receptor distance (Fig. 7M) showed clear regional structure: for D1-ALFA, Amg distances are most left-shifted (closest coupling) relative to Str and Ctx, consistent with a gradient of nanoscale receptor proximity across all three terminal fields. For D2-ALFA, Str exhibits a notably left-shifted distribution relative to Amg, suggesting stronger D2 coupling in striatum (Fig. 7N). The presence of distance distributions that vary based on receptor type and anatomical region, combined with shuffle sensitivity, supports the premise that dopamine receptors are not merely stochastically distributed in axonal terminal fields, but are organized into spatially biased nanoclusters relative to dopamine varicosities.

Together, these findings provide convergent evidence that endogenous dopamine receptors form punctate nanoclusters that are non-randomly positioned relative to dopamine varicosities across major dopaminergic axonal terminal fields. The combination of strong shuffle sensitivity, region-and receptor-specific coupling patterns, puncta size differences at apposed vs. non-apposed receptors, and distance-CDF structure supports a model in which dopamine neuromodulation engages in synapse-like, spatially restricted microdomains, with the balance between targeted appositional signaling and broader extrasynaptic modulation varying by anatomical context and receptor subtype.

## Discussion

A prevailing framework for neuromodulatory communication is volume transmission, in which signaling molecules released from axons diffuse through extracellular space to engage receptors at variable distances while competing with uptake, enzymatic degradation, and bulk clearance. This model captures the fact that neuromodulators can influence many targets over extended spatial scales. However, the model also leaves unresolved how such a comparatively imprecise, one-to-many mode of communication supports the highly specific computations attributed to dopamine circuits (*19, 20*). In particular, key mechanistic questions remain unanswered, namely (1) how neuromodulatory axons identify and stabilize within their terminal fields, (2) whether they preferentially engage particular cellular targets in their terminal fields, and (3) what spatiotemporal constraints govern receptor activation in densely innervated neuropil where receptors are broadly distributed. Even with a mature understanding of GPCR signaling cascades, the rules by which neuromodulatory axons find “where to signal” and how release events are translated into organized receptor engagement remain incompletely defined.

In this study, we converge on a model in which dopamine transmission frequently exhibits synapse-like organization, providing an anchoring principle for spatial precision that may be broadly applicable to monoaminergic neuromodulatory signaling. This interpretation was motivated initially by functional observations that are difficult to reconcile with purely stochastic diffusion. In acute slices and cultured preparations, dopamine release is not spatially uniform but instead forms reproducible hotspots, suggesting that dopamine output is organized into discrete release-competent sites rather than being an interchangeable property of all varicosities.

Extending this to a mechanistic test, single-bouton optical recordings in co-cultures indicated that release competence is tightly linked to appositional coupling: dopamine boutons become release competent specifically when they contact dopamine receptor–positive soma and dendrites, implying that the local postsynaptic environment is not merely a recipient of diffusing transmitter but participates in specifying where release occurs. These functional data helped us establish a key premise that dopamine signaling can be spatially and structurally constrained, and motivated a search for the anatomical substrates that support such constraints.

Our anatomical and molecular analyses in intact tissue provided convergent evidence for those substrates. Using EASI-FISH in amygdala and the striatum, and in co-culture, we observed that dopamine axons exhibit pronounced tropism for receptor-expressing processes, consistent with a model in which neuromodulatory projections may be guided by molecular features of their targets. This addresses a longstanding conceptual gap in volume transmission models: if synaptic coupling is dispensable for release of neuromodulators, it is unclear what rules govern how neuromodulatory axons navigate to and stabilize within appropriate terminal fields, often millimeters away from their midbrain origins. Tropism for receptor-positive cellular microenvironments provides a plausible mechanism for both targeting and local stabilization, and it predicts that synapse-like contact-dependent mechanisms could influence bouton maturation and release competency.

Consistent with these observations, the ExM analyses presented here provide nanoscale resolution of how dopamine axons and receptors are arranged once axons reach their targets.

Across striatum, amygdala, and prefrontal cortex, endogenous D1 and D2 receptors are organized into discrete nanoclusters and are preferentially positioned near dopamine varicosities at distances significantly smaller than expected from receptor density/chance encounter alone.

Conservative shuffle controls that preserve receptor geometry and spatial heterogeneity while disrupting alignment markedly reduce apparent coupling, indicating that the observed proximity reflects structured organization rather than random overlap. Moreover, receptor puncta that are apposed to dopamine varicosities are larger than non-apposed puncta, suggesting that apposition is associated with a clustering similar to those observed in classical synapses. Together, these results place sub-cellular receptor clustering and varicosity apposition at the center of a synapse-like model in which local microdomain structure helps define where dopamine signaling is most efficient and potentially most functionally consequential.

This spatially restricted interpretation also helps reconcile why only a subset of dopamine varicosities appear release competent. Under a purely volumetric model, heterogeneity is often treated as stochastic sampling within dense arbors or as an intrinsic presynaptic property. In contrast, our combined functional and anatomical results support a scenario in which contact-dependent release-maturation is a key determinant of functional release sites. In this model, appositional coupling may serve as the anatomical “selection” step that stabilizes and concentrates release machinery at particular varicosities while leaving others comparatively silent. Consistent with this, we find that presynaptic components classically associated with rapid neurotransmitter release are enriched at dopamine-releasing varicosities (Bsn, RIM, Stx1 and others) and that ultrastructural analysis by TEM reveals synapse-like contact at a subset of dopamine boutons. Importantly, the ultrastructure is not simply a copy of canonical fast synapses; instead, it exhibits nuanced differences consistent with a specialized form of coupling optimized for neuromodulatory signaling. These molecular and ultrastructural data provide a mechanistic bridge between bouton-level release hotspots and the receptor-varicosity architecture revealed by ExM.

Finally, our findings do not argue that diffusion is irrelevant per se; rather, they suggest that stochastic release and diffusion is unlikely to be the primary organizing principle for dopamine signaling. Dopamine released at synapse-like microdomains may still spill over to influence peri-synaptic or extrasynaptic receptors, particularly under high activity or in microenvironments with reduced clearance. However, anchoring release to receptor-proximal sites provides an efficient strategy for achieving the high local concentrations and rapid kinetics needed to reliably engage GPCR pathways under physiological constraints. This is consistent with prior slice-based observations that certain forms of dopamine-dependent GPCR responses, including D2-mediated GIRK currents, are preferentially driven by rapidly rising, high-amplitude dopamine transients rather than by slow, low-level tone (*51, 52*). In our model, synapse-like release–receptor appositions represent a fundamental unit of neuromodulatory organization that create spatial precision and biochemical efficiency while still permitting broader modulation through spillover and network-level integration. Thus, we propose that dopamine transmission is best understood as a hybrid system in which synapse-like microdomains provide the core substrate for precise signaling, embedded within a context where diffusion can extend influence beyond the immediate site subject to activity vigor and local molecular landscape.

## Acknowledgments

We would like to thank Phuong Nguyen for helping with primary culture preparations; Morgan Clarke, Monique Copeland and Benjamin Foster for assistance with histology work, FISH and antibody screening tests. We would like to thank Janelia Viral Tools (Hyun Ah Yi) with packaging viruses and Janelia Molecular Biology team for help with plasmid preparations. We would like to thank Wei Wu on assistance with quantitative proteomics. We would like to thank Eric Schreiter, Alison Tebo and Tim Brown for discussions related to cAMP imaging. We are grateful for the helpful guidance from Paul Tillberg and Mojtaba Tavakoli on expansion microscopy experiments. We would like to thank Marla Feller and Bernardo Sabatini for helpful feedback on the manuscript. **Funding:** This work was funded in full by the Howard Hughes Medical Institute. **Author Contributions:** C.B and A.G.B. designed the study and performed most imaging experiments. C.B. and A.G.B. carried out most of the data analysis, produced the figures and wrote the manuscript with assistance from the rest of the authors. D.W. carried out the primary neuronal preparations with assistance from C.B. N.I. performed most of the electron microscopy experiments with assistance from C.B. RNA FISH probe design and EASI-FISH experiments were carried out by M.E. and C.B. A.G.B developed most of the image analysis code with assistance from D.A. and C.B. Authors declare that they have no competing interests.

## Supplementary Materials

Materials and Methods

Tables S1 to S3

Figs. S1 to S13

Animation 1

References

## References and Notes

1. B. L. Sabatini, W. G. Regehr, Timing of synaptic transmission. Annu Rev Physiol 61, 521–542 (1999).

2. T. C. Südhof, Neurotransmitter release: the last millisecond in the life of a synaptic vesicle. Neuron 80, 675–690 (2013).

3. P. Greengard, The neurobiology of slow synaptic transmission. Science 294, 1024–1030 (2001).

4. D. A. McCormick, D. B. Nestvogel, B. J. He, Neuromodulation of Brain State and Behavior. Annu Rev Neurosci 43, 391–415 (2020).

5. C. D. Grossman, J. Y. Cohen, Neuromodulation and Neurophysiology on the Timescale of Learning and Decision-Making. Annu Rev Neurosci 45, 317–337 (2022).

6. C. Liu, P. Goel, P. S. Kaeser, Spatial and temporal scales of dopamine transmission. Nature Reviews Neuroscience 22, 345–358 (2021).

7. Ö. D. Özçete, A. Banerjee, P. S. Kaeser, Mechanisms of neuromodulatory volume transmission. Molecular Psychiatry 29, 3680–3693 (2024).

8. K. Fuxe et al., The discovery of central monoamine neurons gave volume transmission to the wired brain. Prog Neurobiol 90, 82–100 (2010).

9. K. Fuxe et al., From the Golgi-Cajal mapping to the transmitter-based characterization of the neuronal networks leading to two modes of brain communication: wiring and volume transmission. Brain Res Rev 55, 17–54 (2007).

10. E. S. Vizi, A. Fekete, R. Karoly, A. Mike, Non-synaptic receptors and transporters involved in brain functions and targets of drug treatment. Br J Pharmacol 160, 785–809 (2010).

11. M. E. Rice, S. J. Cragg, Dopamine spillover after quantal release: rethinking dopamine transmission in the nigrostriatal pathway. Brain Res Rev 58, 303–313 (2008).

12. G. Wildenberg et al., Partial connectomes of labeled dopaminergic circuits reveal non-synaptic communication and axonal remodeling after exposure to cocaine. eLife 10, e71981 (2021).

13. V. M. Pickel, S. C. Beckley, T. H. Joh, D. J. Reis, Ultrastructural immunocytochemical localization of tyrosine hydroxylase in the neostriatum. Brain Res 225, 373–385 (1981).

14. J. J. Bouyer, T. H. Joh, V. M. Pickel, Ultrastructural localization of tyrosine hydroxylase in rat nucleus accumbens. J Comp Neurol 227, 92–103 (1984).

15. V. Paget-Blanc et al., A synaptomic analysis reveals dopamine hub synapses in the mouse striatum. Nature Communications 13, 3102 (2022).

16. T. F. Freund, J. F. Powell, A. D. Smith, Tyrosine hydroxylase-immunoreactive boutons in synaptic contact with identified striatonigral neurons, with particular reference to dendritic spines. Neuroscience 13, 1189–1215 (1984).

17. M. Uchigashima, T. Ohtsuka, K. Kobayashi, M. Watanabe, Dopamine synapse is a neuroligin-2–mediated contact between dopaminergic presynaptic and GABAergic postsynaptic structures. Proceedings of the National Academy of Sciences 113, 4206–4211 (2016).

18. K. K. L. Yung et al., Immunocytochemical localization of D1 and D2 dopamine receptors in the basal ganglia of the rat: Light and electron microscopy. Neuroscience 65, 709–730 (1995).

19. M. W. Howe, D. A. Dombeck, Rapid signalling in distinct dopaminergic axons during locomotion and reward. Nature 535, 505–510 (2016).

20. J. A. da Silva, F. Tecuapetla, V. Paixão, R. M. Costa, Dopamine neuron activity before action initiation gates and invigorates future movements. Nature 554, 244–248 (2018).

21. N. F. Parker et al., Reward and choice encoding in terminals of midbrain dopamine neurons depends on striatal target. Nature Neuroscience 19, 845–854 (2016).

22. L. T. Coddington, J. T. Dudman, The timing of action determines reward prediction signals in identified midbrain dopamine neurons. Nature Neuroscience 21, 1563–1573 (2018).

23. J. E. Markowitz et al., Spontaneous behaviour is structured by reinforcement without explicit reward. Nature 614, 108–117 (2023).

24. C. Ducrot et al., Dopaminergic neurons establish a distinctive axonal arbor with a majority of non-synaptic terminals. Faseb j 35, e21791 (2021).

25. L. Descarries, K. C. Watkins, S. Garcia, O. Bosler, G. Doucet, Dual character, asynaptic and synaptic, of the dopamine innervation in adult rat neostriatum: a quantitative autoradiographic and immunocytochemical analysis. J Comp Neurol 375, 167–186 (1996).

26. L. Descarries, N. Mechawar, in Progress in Brain Research. (Elsevier, 2000), vol. 125, pp. 27–47.

27. C. Liu, L. Kershberg, J. Wang, S. Schneeberger, P. S. Kaeser, Dopamine Secretion Is Mediated by Sparse Active Zone-like Release Sites. Cell 172, 706–718.e715 (2018).

28. A. G. Salinas et al., Distinct sub-second dopamine signaling in dorsolateral striatum measured by a genetically-encoded fluorescent sensor. Nature Communications 14, 5915 (2023).

29. C. Bulumulla et al., Visualizing synaptic dopamine efflux with a 2D composite nanofilm. eLife 11, e78773 (2022).

30. A. G. Beyene et al., Imaging striatal dopamine release using a nongenetically encoded near infrared fluorescent catecholamine nanosensor. Science Advances 5, eaaw3108 (2019).

31. R. Y. Moore, F. E. Bloom, Central catecholamine neuron systems: anatomy and physiology of the dopamine systems. Annu Rev Neurosci 1, 129–169 (1978).

32. X. Jiang, R. Sando, T. C. Südhof, Multiple signaling pathways are essential for synapse formation induced by synaptic adhesion molecules. Proc Natl Acad Sci U S A 118, (2021).

33. P. Scheiffele, J. Fan, J. Choih, R. Fetter, T. Serafini, Neuroligin Expressed in Nonneuronal Cells Triggers Presynaptic Development in Contacting Axons. Cell 101, 657–669 (2000).

34. G. Gopalakrishnan et al., Label-free visualization of ultrastructural features of artificial synapses via cryo-EM. ACS Chem Neurosci 2, 700–704 (2011).

35. K. Suzuki et al., A synthetic synaptic organizer protein restores glutamatergic neuronal circuits. Science 369, eabb4853 (2020).

36. L. Wang et al., A high-performance genetically encoded fluorescent indicator for in vivo cAMP imaging. Nature Communications 13, 5363 (2022).

37. Y. Wang et al., EASI-FISH for thick tissue defines lateral hypothalamus spatio-molecular organization. Cell 184, 6361–6377.e6324 (2021).

38. Ł. Bijoch et al., Opposing effects of rewarding and aversive stimuli on D1 and D2 types of dopamine-sensitive neurons in the central amygdala. bioRxiv, 2024.2011.2007.622413 (2024).

39. P. Kirkpatrick, Specificity concerns with antibodies for receptor mapping. Nature Reviews Drug Discovery 8, 278–278 (2009).

40. H. Götzke et al., The ALFA-tag is a highly versatile tool for nanobody-based bioscience applications. Nat Commun 10, 4403 (2019).

41. L. Kershberg, A. Banerjee, P. S. Kaeser, Protein composition of axonal dopamine release sites in the striatum. Elife 11, (2022).

42. H. V. Friedman, T. Bresler, C. C. Garner, N. E. Ziv, Assembly of New Individual Excitatory Synapses: Time Course and Temporal Order of Synaptic Molecule Recruitment. Neuron 27, 57–69 (2000).

43. M. Shapira et al., Unitary Assembly of Presynaptic Active Zones from Piccolo-Bassoon Transport Vesicles. Neuron 38, 237–252 (2003).

44. B. G. Robinson et al., RIM is essential for stimulated but not spontaneous somatodendritic dopamine release in the midbrain. eLife 8, e47972 (2019).

45. S. S. Lam et al., Directed evolution of APEX2 for electron microscopy and proximity labeling. Nature Methods 12, 51–54 (2015).

46. B. Imbrosci, D. Schmitz, M. Orlando, Automated Detection and Localization of Synaptic Vesicles in Electron Microscopy Images. eneuro 9, ENEURO.0400-0420.2021 (2022).

47. A. Björklund, S. B. Dunnett, Dopamine neuron systems in the brain: an update. Trends in Neurosciences 30, 194–202 (2007).

48. J.-F. Poulin et al., Mapping projections of molecularly defined dopamine neuron subtypes using intersectional genetic approaches. Nature Neuroscience 21, 1260–1271 (2018).

49. M. Azcorra et al., Unique functional responses differentially map onto genetic subtypes of dopamine neurons. Nature Neuroscience 26, 1762–1774 (2023).

50. M. R. Tavakoli et al., Light-microscopy-based connectomic reconstruction of mammalian brain tissue. Nature 642, 398–410 (2025).

51. A. G. Yee, Y. Liao, B. S. Muntean, C. P. Ford, Discrete spatiotemporal encoding of striatal dopamine transmission. Science 389, 200–206 (2025).

52. C. P. Ford, P. E. Phillips, J. T. Williams, The time course of dopamine transmission in the ventral tegmental area. J Neurosci 29, 13344–13352 (2009).

