## Supplementary Information for "Synapse-like Specializations at Dopamine Release Sites Orchestrate Efficient and Precise Neuromodulatory Signaling"

### Materials and Methods

All chemical reagents and antibodies are listed in Supplementary Tables 1-3. Synthesis of the nanosensors (nIRCat) and fabrication of DopaFilm have been published previously (*1, 2*). Plasmid for hSyn-G-Flamp1 (Addgene#188569) was purchased from Addgene and pAAV–EF1a-DIO-APEX2.NES construct was subcloned into a lenti backbone by VectorBuilder. Lentiviruses containing shRNA knock down sequences were sourced from VectorBuilder. Virus packaging was performed at HHMI Janelia research campus. All data analysis was performed using MATLAB_R2024a unless otherwise noted.

#### Supplementary Table 1. List of chemical reagents.

| Chemical | Abbreviation | Vendor | Identifier |
|---|---|---|---|
| Single-walled carbon nanotubes | SWCNT | NanoIntegris | HR35-141 |
| 5'-GTG TGT GTG TGT-3' | $(GT)_6$ | IDT | N/A |
| Dopamine hydrochloride | DA | Sigma-Aldrich | H8502 |
| (3-Aminopropyl) triethoxysilane | APTES | Sigma-Aldrich | 440140 |
| Ethanol | EtOH | Fisher Scientific | A405P |
| 32% Paraformaldehyde | PFA | EMS | 15714 |
| Glycine | N/A | Sigma-Aldrich | G7126 |
| Triton X-100 | N/A | Fisher Scientific | BP151 |
| Tween-20 | N/A | Sigma-Aldrich | P7949 |
| NbActiv4 | N/A | Transnet/ Brain Bits | NC1477957 |
| Acrylamide | AA | Sigma-Aldrich | A9099 |
| Sodium acrylate | SA | AK Scientific | R624 |
| N,N'-Methylenebisacrylamide | Bis | Sigma-Aldrich | M7279 |
| Ammonium persulfate | APS | Bio Rad | 1610700 |
| Sodium bicarbonate | $NaHCO_3$ | Oakwood Chemicals | 099274 |
| N,N,N,N'-Tetramethylethylenediamine | TEMED | Bio Rad | 161-0800 |
| Sodium azide | N/A | Sigma-Aldrich | 71289 |
| Glycidyl glycerol ether | TGE | Polysciences Europe GmbH | 09221 |
| Glycidyl methacrylate | GMA | TCI | G0497 |
| Sodium chloride | NaCl | Fisher Scientific | 190383 |

| | | | |
|---|---|---|---|
| 4,6’-diamino-2-phenylindole dihydrocholride | DAPI | Sigma-Aldrich | D9542 |
| 4-(2-Hydroxyethyl)piperazine-1-ethanesulfonic acid | HEPES | Sigma-Aldrich | H3375 |
| 20X Sodium-saline citrate buffer | 20X SSC | Invitrogen | AM9763 |
| (Tris(hydroxymethyl)aminomethane) | 1M Tris pH 8 | Invitrogen | AM9856 |
| D-Glucose | N/A | Acros Organics | 41095 |
| Sodium phosphate monobasic | $NaH_2PO_4$ | Sigma-Aldrich | P9791 |
| Sodium cacodylate trihydrate | SC | Sigma-Aldrich | C0250 |
| Osmium tetraoxide | $OsO_4$ | Electron microscopy Sciences | 19150 |
| Potassium ferrocyanide | $K_4Fe(CN)_6$ | Macron | 6932-04 |
| Uranyl acetate | N/A | Ibi Labs | U73493-25-ACS |
| Eponate 12$^{TM}$ kit | | Ted Pella | 18012 |
| (3-(N-Morpholino)propane sulfonic acid) | MOPS | Fisher Scientific | BP308 |
| Melphalan | N/A | Cayman Chemicals | 16665 |
| (6-((acryloyl)amino)hexanoic acid, succinimidyl ester) | AcX | ThermoFisher Sci. | A20770 |
| Isoflurane | N/A | Fisher Scientific | 50-304-2993 |
| poly-L-lysine | N/A | Pelco | 18026 |
| Poly-D-lysine | N/A | Sigma-Aldrich | P6407 |
| Bovine serum albumin | BSA | Sigma-Aldrich | A9418 |
| Calcium chloride | $CaCl_2$ | Acros Organics | 349615000 |
| Magnesium sulfate | $MgSO_4$ | Fisher Scientific | M65 |
| Potassium chloride | KCl | Fisher Scientific | P217 |
| Glutamine | N/A | Fisher Scientific | 25-030-149 |
| Heat inactivated fetal bovine serum | FBS | Fisher Scientific | SH30071.03 |
| MEM media | N/A | Invitrogen | 51200-038 |
| Transferrin | N/A | Sigma-Aldrich | 616420 |
| Insulin | N/A | Sigma-Aldrich | I-6634 |
| Papain enzyme | N/A | Worthington Biochemical | PAP2 |
| Drd1 and Drd2 HCR Probes | N/A | Molecular Instruments | N/A |
| RNAScope probes (Drd1/2) and reagents | N/A | Advanced Cell Diagnostics | N/A |
| Opal 520 and Opal 690 | N/A | Akoya Biosciences | N/A |
| Water | N/A | Fisher Scientific | W6-4 |
| Acetonitrile | ACN | Fisher Scientific | A9554 |
| Methanol | N/A | Fisher Scientific | A454SK-4 |
| Formic acid | N/A | Pierce | PI28905 |
| Acetone | N/A | Fisher Scientific | A949-1 |
| RapiGest SF surfactant | N/A | Waters | 186001861 |
| Dithiothritol | DTT | Thermo Scientific | R0861 |
| Trifluoroacetic acid | TFA | Sigma-Aldrich | 302031 |
| Iodoacetamide | IAA | Sigma-Aldrich | 407710 |
| Ammonium bicarbonate | N/A | Sigma-Aldrich | 40867 |
| Trypsin | N/A | Promega | V5113 |

### Supplementary Table 2. List of primary antibodies

| Target | Vendor | Identifier | Host species | Dilution factor |
|---|---|---|---|---|
| Tyrosine hydroxylase (TH) | Aves Labs | TYH 67677979 | Chicken | 1:1000 |
| Tyrosine hydroxylase (TH) | Sigma-Aldrich | AB152 | Rabbit | 1:1000 |
| Bassoon (BSN) | Abcam | Ab82958 | Mouse | 1:1000 |

| | | | | |
|---|---|---|---|---|
| RIM1/2 | Synaptic Systems | SySy 140 205 | Guinea pig | 1:1000 |
| Munc13-1 | Synaptic Systems | SySy 126 103 | Rabbit | 1:1000 |
| Synaptophysin1 (SYP) | Synaptic Systems | SySy 101 308 | Guinea pig | 1:1000 |
| Synaptobrevin2 (VAMP2) | Synaptic Systems | SySy 104 318 | Guinea pig | 1:1000 |
| ALFA antibody | NanoTag Biotech | N1586 | Human | 1:300 |
| ALFA nanobody Atto643 | NanoTag Biotech | N1502-At643-L | Alpaca | 1:200 |
| RFP | ThermoFisher | 600-401-379 | Rabbit | 1:1000 |
| MAP2 | Novus biologics | NB300-213 | Chicken | 1:1000 |
| GFAP | Sigma-Aldrich | G3893 | Mouse | 1:1000 |

**Supplementary Table 3. List of secondary antibodies**

| Target | Host | vendor | Identifier | Fluorophore | Dilution factor |
|---|---|---|---|---|---|
| Chicken IgY (H+L) | Donkey | Jackson Immuno. | 703-675-155 | BV 421 | 1:1000 |
| Chicken IgY (H+L) | Goat | ThermoFisher Sci. | A78948 | AF Plus 488 | 1:2000 |
| Chicken IgY (H+L) | Goat | ThermoFisher Sci. | A32933TR | AF Plus 647 | 1:2000 |
| Mouse IgG/IgM (H+L) | Goat | ThermoFisher Sci. | A10680 | AF 488 | 1:2000 |
| Mouse IgG (H+L) | Goat | ThermoFisher Sci. | A32728TR | AF 647 | 1:2000 |
| Rabbit IgG (H+L) | Goat | ThermoFisher Sci. | A11034 | AF 488 | 1:2000 |
| Rabbit IgG (H+L) | Goat | ThermoFisher Sci. | A11037 | AF 594 | 1:2000 |
| Rabbit IgG (H+L) | Goat | Abberior | STRed-1002 | STAR Red | 1:2000 |
| Rabbit IgG (H+L) | Donkey | Biotium | #20125 | CF 633 | 1:2000 |
| Guinea pig IgG (H+L) | Goat | ThermoFisher Sci. | A11073 | AF 488 | 1:2000 |
| Guinea pig IgG (H+L) | Goat | Abberior | STRed-1006 | STAR Red | 1:2000 |
| Human IgG (H+L) | Goat | ThermoFisher Sci. | A21089 | AF 546 | 1:300 |

### Ethical considerations and mouse strains

All experiments were conducted according to the Institutional Animal Care and Use Committee (IACUC) guidelines of HHMI Janelia Research Campus. For culture and tissue experiments, we used *Slc6a3*-cre(*3, 4*)x Ai9, ALFADoR1 mice (this work), ALFADoR2 mice (this work) transgenic P1–P2 pups and adult (P50 – P60) mice, respectively.

### ALFADoR1/2 knock-in mouse generation and validation

Both transgenic mouse lines were generated by CRISPR-cas9 pronuclear microinjection by the transgenics facility at HHMI Janelia research campus. Male mice older than 8 weeks were mated with 24-27 days old superovulated C57Bl/6J females (from Jackson laboratory, JAX) to obtain zygotes for pronuclear injection. CD-1 Elite females (Charles River laboratories, CRL) were used as embryo transfer recipients. Pronuclear-stage embryos were collected in M2 (Tribioscience; TBS8070-50mL), washed three times in KSOM (Tribioscience; TBS8071-50mL) to remove cumulus cells, and incubated in KSOM at 37 °C, 5% $CO_2$. The mix of Cas9 ribonucleoprotein (RNP) and donor double-stranded DNA was injected into the pronuclei of the zygotes. Following injection, embryos were incubated in KSOM until all injections were complete. Approximately 10–15 embryos were transferred into the oviducts of pseudo-pregnant CD-1 Elite recipient females on the same day. Cas9 protein (TRUECUT CAS9 PROTEIN V2, 100UG) was purchased from Life Tech Corp (A36498). The sgRNA (5’-GTTAGGAGCCATCTTCCAGAAGG -3’) was generated by in vitro transcription using

MEGAshortscript T7 kit (LIFE TECH CORP; AM1354) and purified with MEGAclear™ Transcription Clean-Up Kit (LIFE TECH CORP; AM1908). The following donor DNA (500bp double-stranded fragment) was synthesized by Integrated DNA Technologies (IDT).

***Drd1 ALFA-tag donor sequence:***

GGTGGAAAGCAAGCGGCTCTTCTTCCTGGTATGGCTTGGATTGCTATGGAGATGCTCCTGATGGAACA
CCATTGTGCTTTTGTCCAGACAGCAACTGGGGCTGGAGAAGGGGCTGGGTGGTGAGTGATTGGGGGAA
GTCTGGCTAAGCCTGGCCAAGAACGTGAGGGCTAAGCCACCGGAAGTGCTTTCCTTCTGGAAGGCCAC
CATGccgagccgcctggaagaagaactgcgccgccgcctgaccgaaccgGCTCCTAACACTTCTACCATGGATGAGACTGGCCT
GCCAGTGGAGAGGGACTTCTCCTTTCGCATCCTCACAGCCTGTTTTCTGTCCCTGCTTATCCTGTCCACT
CTCTTAGGGAATACCCTTGTCTGTGCCGCTGTCATCAGGTTTCGACACCTGCGGTCCAAGGTGACCAAC
TTCTTTGTCATCTCTTTAGCTGTGTCAGATCTCTTGGTGGCTGTCTTGGTCATGCCCTGGAAAGCTGTGG
CTG

***Drd2 ALFA-tag donor sequence:***

GCTAGAAGAACAGAAGCTTGTCCTAACATTATAAGATCGCTAGAACCAGGAATAAAAAAAACATAGT
TTGGGGAATTCTCAGCTCTGCTAGCTCTTGGTTTTTCTGCAGGGAATCCTCTTTAGGAGGAAGCATGCC
TTGAAAACACTCCTGCTCACTCCTTGTATTCATTTCTCCCGGCCAGAGCCGTGCCACCCAGTGGCCCCA
CTGCCCCAGCCACCATGccgagccgcctggaagaagaactgcgccgccgcctgaccgaaccgGATCCACTGAACCTGTCCTGGT
ACGATGATGATCTGGAGAGGCAGAACTGGAGCCGGCCCTTCAATGGGTCCGAAGGGAAGCCAGACAG
GCCCCACTACAACTACTATGCCATGCTGCTCACCCTCCTCATCTTTATCATCGTCTTTGGCAATGTGCTG
GTGTGCATGGCTGTATCCAGAGAGAAGGCTTTGCAGACCACCACCAACTACCTGATAGTCAGCCTCGC
TGTGGC

The ALFA tag sequence (PSRLEEELRRRLTEP) was inserted into the second exon (E2) of either Drd1 and Drd1 genes, resulting in a fusion with the endogenous gene. For Drd1 and Drd2, the inserts were flanked by left and right homology arms of 199 bp/247 bp and 213 bp/233 bp, respectively. For microinjections, the Cas9 protein/sgRNA/double-stranded DNA donor mix was suspended in 0.22 µm-filtered 10mM Tris-Cl buffer (PH7.5) at 30 ng/ 30 ng/ 8ng for Drd1 and 5 ng for Drd2, respectively. $F_0$ mice were genotyped using PCR with primers indicated below.

| Gene | Target | Forward | Reverse |
|---|---|---|---|
| Drd1 | 5'-junction | TGGAGCACTGAACCCAGAAG | CCAGTCTCATCCATGGTAGA |
| | 3'-junction | CTAAGCCACCGGAAGTGCTT | CGTGGAGCACATGATGTCAA |
| Drd2 | 5'-junction | GAAAGAATGGATGAGTTAGCTC | GATCATCATCGTACCAGGAC |
| | 3'-junction | GAAAGAATGGATGAGTTAGCTC | AGTGTGGCCACCAGAAGATC |

Targeted insertion was further confirmed by Sanger sequencing using the following primers:

Drd1 5F: 5'-TGGAGCACTGAACCCAGAAG-3', 3R: 5'-CGTGGAGCACATGATGTCAA-3' (WT 543 bp; mut 594 bp).

Drd2 5F: 5'-GAAAGAATGGATGAGTTAGCTC-3', 3R: 5'- AGTGTGGCCACCAGAAGAT C-3' (WT 549 bp; mut 600 bp).

**Preparation of Primary Mouse Neuronal Cultures**

Primary mouse neuronal culture work was carried out as previously published (*2*) with some modifications. Briefly, *Slc6a3* x Ai9 transgenic neonatal pups (P1 – P2) were euthanized and tissue sections from the cortex, striatum, and substantia nigra pars compacta (SNc) and ventral tegmental Area (VTA) from midbrain region were dissected. We used a fluorescence guided dissection microscope (Zeiss SteREO Discovery.V20) to identify and maximize the extraction of tdTomato (tdTom) expressing cells in SNC and VTA regions. Mixed co-cultures were generated by extracting dopamine neurons in the SNC + VTA regions from *Slc6a3* x Ai9 transgenic mice and cortical or striatal regions from ALFADoR1/2 mice. SNC + VTA cells were seeded as co-cultures with either cortical cells or striatal cells at a ratio of 3:7, and density of 30k cells per 35mm MatTek dish (14mm insert), in attachment media (1:1 v/v plating media to NbActiv4) on top of DopaFilm substrates. Cultures were initially maintained in attachment media for 3h in a 5% $CO_2$ humid incubator at 37 °C. Next, attachment media was replaced with growing media (plating media: NbActiv4 1:20 v/v) and cells were allowed to grow for one week. After one week, NbActiv media (500 µl) was added on top of growing media and cultures were fed twice a week. Viral transductions with hSyn-G-Flamp1 (AAV2), pAAV–EF1a-DIO-APEX2.NES and knock-down (KD) constructs (lentis) for presynaptic proteins were carried out around 5-7 days in vitro (DIV 5-7). shRNA KD constructs were designed using the Genetic Perturbation Platform (GPP, Broad Institute). We chose the top three-scoring KD constructs per gene, based on KD scores reported on the web portal. Each gene perturbation experiment used equal titers of the three constructs simultaneously.

**Data acquisition and analysis for neuronal activity imaging**

We carried out activity imaging in neuronal co-cultures (DIV14 – 21) using a custom-built broad spectrum (400 –1400 nm) epifluorescence microscope with a 25x water immersion objective (Olympus) and two cameras (SWIR/NIR on NiNox 640 II, Raptor photonics, and visible on Hamamatsu ORCA FusionBT (C15440)) built on a Thorlabs Bergamo microscope body. Prior to activity imaging experiments, we aspirated the NbActiv4 (growth media) and washed cells 2X with fresh ACSF and incubated them with 5ppm nanosensors in ACSF for 1h inside an incubator (5% $CO_2$, 37°C). The ACSF composition was as follows in mM: 124 NaCl, 2.5 KCl, 1.25 $NaH_2PO_4$, 24 $NaHCO_3$, 12.5 Glucose, 5 HEPES, 2 $CaCl_2$, 2 $MgSO_4$. We then washed cells 2X and mounted the dish on the microscope stage for imaging. Before imaging sessions, we first saved the tdTom, BF images and switched to the NIR channel for DopaFilm activity measurements. GFP based sensors and tdTom positive axons were excited using a LED light source (Thorlabs LED4D067) and DopaFilm/NIRCats were excited using a 785 nm laser (Thorlabs S4FC785) with 100mW power at the source, giving ~30 mW at the sample. For dopamine activity imaging experiments, 500 images were acquired with µManager (*5*), at ~10 frames per second at room temperature (RT). To reliably evoke dopamine release, field stimulation (5 pulses at 25Hz, 10 ms pulse width) was applied to cultured neurons using custom-made platinum electrodes connected to a stimulus isolator (A385, World Precision Instruments) after 15 seconds of baseline activity were collected.

We performed analysis of dopamine imaging data as previously reported using a custom-built MATLAB code (*2*), available at: https://github.com/JaneliaSciComp/nnmf/commits/main/.

For analysis, raw image files were first convolved with a 2D gaussian (σ = 0.5 pixels). To obtain baseline intensity values at each pixel (i.e., $F_0$), we used a leaky cumulative minimum and applied repeated low pass filtering to converge on a smooth $F_0$. We then computed ΔF as $F-F_0$. Next, we applied nonnegative matrix factorization (NNMF) to ΔF with sparsity and contiguity constraints to partition the image into correlated components. NNMF is a dimensionality reduction method that segments the image into correlated regions (independent components) that best approximate the underlying image. This effectively groups pixels with similar activity into components, and these components spatially colocalize with the fluorescence hotspots detected on DopaFilm. NNMF masks were then used to compute ΔF/F traces corresponding to each hotspot. For clustering analysis related to Figure 2 data (and presented as a summary in Figure 2H-J), we first binarize the tdTom channel and classify active and quiescent varicosities based on their overlap with ΔF channel/ We then compute the pairwise distances between tdTom pixels at ΔF hotspots (active), non-hotspots (quiescent) or over all tdTom pixels (i.e., all boutons). Proximity analysis for an individual region was performed by taking the distance transform of a binarized MAP2 image. The distance from MAP2 as a function of normalized ΔF is plotted, with error bars showing 95% confidence interval.

**Proximity and Odds Ratio analysis in mixed-culture datasets**

We used three co-registered 2D TIFF images (single field per analysis), including (i) tdTomato (tdTom) labeling dopamine axons, (ii) D1 or D2 dopamine receptor immunofluorescence, and (iii) ΔF/F images reporting dopamine release. Images were converted to grayscale and intensity-normalized. Background was reduced with a morphological top-hat filter. Axons were segmented from the tdTom channel using adaptive thresholding, followed by small-object removal and hole filling. To minimize bias from axon thickness, we skeletonized the axon mask. Receptor puncta were detected from a Laplacian-of-Gaussian (LoG) response image ($\sigma \approx 0.8$–2.0 px) thresholded on z-scored intensity (typically $Z>2$–4), constrained by local maxima, then cleaned by area filtering (to remove small and large objects). For robustness to the point-spread function and minor misregistration, we optionally dilated the receptor mask by 2 px before distance computation. Release regions (ΔF/F) were segmented by adaptive thresholding minimum area filtering (remove large and small objects).

We computed a Euclidean distance map to the nearest receptor pixel. Distances were sampled over axon pixels and partitioned into release+ (axon ∩ ΔF/F, active) and release− (axon ∩ not-ΔF/F, quiescent). We summarized the distributions with empirical cumulative distribution functions (ECDFs). For odds ratio calculations, for radii r spanning 0.3–2.0 µm, we defined proximity as distance $\leq r$. For each r, we tabulated axon pixels within vs. outside r for release+ and release− groups and computed the odds ratio (OR) as fraction (or percentage) of active varicosities that are at $\leq r$ divided by fraction (or percentage) of quiescent varicosities that are at $\leq r$. OR values close to 1 indicate no difference in enrichment. We estimated 95% confidence intervals for OR by bootstrapping log(OR) with resampling within each group (4,000 iterations) and exponentiating the 2.5th–97.5th percentiles. We visualized OR versus r with corresponding bootstrap CIs; values above 1 indicate enrichment of D1R proximity among release+ axon pixels.

**Acute slice imaging**

Acute coronal brain slices containing dorsomedial striatum or amygdala were prepared as described previously, and extracellular dopamine dynamics was recorded using nIRCats (*1*). Briefly, acute slices were labeled with dopamine nanosensors in solution phase via passive incubation at room temperature (15 minutes) in ACSF containing 2 µg/mL of the nanosensors. Slices were allowed to equilibrate in a submerged recording chamber under continuous perfusion of oxygenated ACSF (95% $O_2$ / 5% $CO_2$) and transferred to an imaging chamber under perfusion for activity recording. Dopamine release was monitored optically by imaging dopamine-evoked fluorescence transients in the field of view during local electrical stimulation. A bipolar stimulating electrode was positioned on the slice surface within the imaging region under low magnification (4x). Then, a high-magnification objective (25x, Olympus) was used to select a field of view at a distance of ~100 µm from the electrode for activity recording. Time-lapse imaging was acquired at 10 frames per second, with baseline collected for 15 seconds before stimulation; each acquisition lasted 500 frames. Single-pulse electrical stimuli (0.3 mA, 1 ms) were delivered to evoke dopamine release using WPI A365 Stimulus Isolator.

**Post-hoc Immunocytochemistry, Confocal and Airyscan Imaging**

Following activity recording, the imaging buffer (ACSF) was aspirated, and the specimens were fixed with 4% PFA at RT for 10 min. Fixative was then aspirated, and cells were washed 3X with DPBS. After blocking with blocking buffer (2% BSA in 0.1% Triton X-100 in PBS) for 1h at RT, tissues were incubated with primary antibodies diluted in antibody dilution buffer (1% BSA in 0.1% Triton X-100 in PBS) overnight at 4 °C. The samples were then rinsed multiple times with PBS and incubated with secondary antibodies diluted in antibody dilution buffer for 3h at RT. To probe the ALFA-tag in fixed cultured cells, we performed an extra heat induced epitope retrieval step using 5% urea in 100mM Tris buffer (pH 9.5) for 10mins at 80 °C. Standard confocal and Airyscan images were acquired on a Zeiss LSM980 inverted microscope equipped with a Plan-Apochromat 40X or a 63X (1.2 NA water or 1.4 NA oil, respectively) objective.

**Immunohistochemistry on brain slices**

Transgenic adult mice were anesthetized with isoflurane and perfused transcardially, sequentially with cold PBS and 4% PFA (50mL each). After perfusions, isolated brains were sectioned using a vibratome (Leica VT1000 S) to get 50µm thick coronal brain slices. After blocking with blocking buffer for 1h at RT, tissues were incubated with primary antibodies (rabbit anti-TH; 1:500 and human anti-ALFA; 1:250) diluted in antibody dilution buffer overnight at 4 °C. The samples were then rinsed multiple times with PBS and incubated with secondary antibodies diluted in antibody dilution buffer for 3h at RT. Tissue sections were mounted onto glass slides (Fisher Scientific, cat. #.12-550-15) and #1.5 glass coverslips (EMS, cat # 63793-01) were applied using VECTASHIELD mounting medium (Vector laboratories, part # H-1400-10). Confocal imaging was carried out using a spinning disk confocal microscope (Nikon CFI Plan Apo Lamda 60X oil/1.4NA). Images were denoised, deconvoluted, and stitched using the in-built NIS elements software.

### Data acquisition and analysis of EASI-FISH volume

Experimental procedure for EASI-FISH experiments was adapted from a previously published protocol (*6*). Briefly, *Slc6a3* x Ai9 mice were anesthetized with isoflurane and perfused transcardially with 15mL of RNase-free PBS followed by 50mL of ice-cold 4% PFA. After overnight post-fixation in the same fixative, brains were sectioned coronally into 300µm slices. Slices containing amygdala or the striatum were selected for the study. We used TREx-1000 gel for the study and the gel recipe is published elsewhere (*7*). Hybridization probes for Drd1 and Drd2 were used at a concentration of 1 µM and amplified using hairpins conjugated to AF488 and JF669 at a final concentration of 3µM. The endogenous tdTom signal (i.e., dopamine axons) was amplified by IHC using the anti-RFP primary antibody (1:500) followed by an anti-rabbit secondary antibody conjugated AF594 (1:200). The gels were imaged with a Zeiss Light Sheet 7 microscope.

EASI-FISH image analysis was performed in 3D using a cell-indexed workflow that enabled cell-wise extraction of receptor signals and dopamine axon-associated fiber/bouton features. Nuclei were segmented from the DAPI channel using Cellpose to generate an indexed 3D label mask. To approximate cell-associated volumes around each nucleus, the indexed mask was then expanded by 8 µm in 3D using an anisotropic, label-preserving distance-transform dilation method. During dilation, background voxels within the dilation radius were assigned the label of the nearest nucleus, preventing higher-index labels from overwriting lower-index labels during expansion.

Using the resulting expanded cell index ID mask, the Drd1, Drd2, and axon channels were clipped in 3D on a per-cell basis. For each cell ID, maximum-intensity projections (MIPs) were generated at an image size of 180 × 180 pixels. Two MIP versions were produced: a two-channel MIP containing only Drd1 and Drd2 signals, and a three-channel MIP containing Drd1, Drd2, and the axon channel. Only the two-channel MIP version was used for machine-learning training, and the trained classifier was applied for four-class separation.

To train the classifier, we manually selected 25 cell-ID MIPs for each of four classes: Drd1-only positive, Drd2-only positive, Drd1 and Drd2 both positive, and both negative. For machine-learning classification of these two-channel MIP datasets into the four experimental classes, we used a lightweight convolutional neural network (CNN) implemented in PyTorch. Each input image was loaded in RGB (not including the axon channel), converted to a tensor, and intensity-normalized. The CNN ("SmallCNN") consisted of three convolutional blocks with 3×3 kernels (padding = 1) and increasing channel depth (16 → 32 → 64). Each convolution was followed by a ReLU nonlinearity and 2×2 max pooling. Features were flattened and passed through a fully connected layer (128 units, ReLU) followed by a final linear layer producing 4 output logits corresponding to the classes: Drd1, Drd2, Both_Negative, and Both_Positive. For inference, model weights were loaded from a saved PyTorch state dictionary (.pth), and classification was performed in evaluation mode with gradient computation disabled. For each image, class probabilities were computed using a softmax operation over the four logits, and the predicted class was assigned as the argmax of the probability vector. The associated confidence score was defined as the maximum softmax probability for the predicted class. Model performance was evaluated using an 80/20 stratified train/validation split. On the held-out validation set (n = 20), the model achieved 95% accuracy (macro-average precision = 0.96, recall = 0.95, F1 = 0.95). Class-wise F1-scores were: Both negative = 1.00, Both positive = 0.89, Drd1 = 0.91, and Drd2 = 1.00. To facilitate downstream analysis and manual review, images were automatically organized into class-specific output folders, and each file was renamed to embed the prediction confidence and predicted class.

Axon fiber and bouton analyses were then performed within each cell ID's 3D index mask (using the 8 μm dilated mask). Entire axon fiber light-microscopy data were segmented in 3D using a direction-selective local-thresholding method (DSLT). The segmented fiber volume was clipped by the 8 μm dilated cell index mask to assign fiber structures to individual cells. For axon fiber quantification, 3D skeletons were created and short terminal spurs were pruned while preserving the main trunk; these short terminal spurs were treated as artificial branches arising from reduced z optical resolution. Fiber length was measured based on skeleton length. For branch point counting, we counted three-branch intermediate points on the 3D skeleton.

Bouton structures were quantified from the axon channel within each cell-ID mask as follows: a maximum-intensity projection of the axon stack was generated and an automatic intensity threshold was estimated on the MIP using ImageJ ("Default dark", 16-bit). The resulting lower threshold was tightened for striatum data (final cutoff = 1.4× lower) and applied to the full 3D synapse volume to generate a binary bouton candidate mask. The mask was denoised using outlier removal and converted into an indexed 3D object map by connected-component labeling (minimum size 3 voxels), yielding bouton candidate volumes.

To estimate fiber geometry, a 3D Euclidean distance transform (EDT) was computed and a diameter threshold was set to 0.1874 μm for amygdala and 0.002 μm in striatum. The bouton mask was skeletonized in 3D; skeleton endpoints and branchpoints were annotated and short terminal spurs were pruned (truncation = 1). Bouton objects were further classified in amygdala using a skeleton-thickness profile matching method. Specifically, a 3D EDT volume was used as a local radius/thickness estimate, and skeleton endpoints and branch points were identified in the skeleton mask. From each endpoint, the algorithm traced the skeleton voxel-by-voxel in 3D (26-neighborhood), propagating until reaching a branch point, while recording along the traced path (i) the EDT-derived thickness values and (ii) the corresponding connected-component bouton index value at each voxel (with a 3×3×3 neighborhood mode fallback when the center voxel index was 0). For each bouton candidate (connected-component ID), the collected thickness sequence was summarized into a compact "shape" by averaging thickness over three equal-length bins, generating a low-dimensional thickness-profile vector. This vector was compared to predefined reference templates representing bouton-like versus neurite-like profiles using cosine similarity. Final bouton calls were made by combining the average thickness and the profile similarity scores (bouton templates versus neuron template).

In the striatum, a 3D EDT was computed to estimate local thickness, and per-object thickness statistics (mean and maximum diameter) were aggregated for each connected-component ID). In preliminary filtering, objects with mean diameter ≈ maximum diameter were treated as trunk-like structures (uniform thickness, consistent with neurite shafts) and were removed first. However, in the striatum many true bouton-like structures were very thin and could still exhibit near-uniform thickness, causing them to be mistakenly classified as trunk-like by diameter-based rules. Because diameter-based thresholding could not reliably separate striatal boutons from thin non-bouton structures without discarding valid boutons, we did not apply any diameter/thickness thresholding in striatum and instead retained all candidates that passed the upstream intensity thresholding. Objects classified as boutons were retained in a filtered bouton-only index volume for downstream quantification of bouton number, volume, and skeleton length. For each cell ID, we reported bouton number, axon volume (scaled by voxel volume), and axon skeleton length (scaled by pixel size).

**Modified RNAScope protocol for cultured neurons**

RNA transcripts for Drd1 and Drd2 genes were probed according to the manufacturer's recommended protocol with some modifications. These modifications include (1) milder target retrieval step (80 °C for 5 min instead of 100 °C for 15 min) and (2) reduced time for Protease III digestion (5 min instead of 30 min). During the RNAScope protocol, tdTom signal becomes heavily attenuated and we included a post ICC step with rabbit anti-RFP antibody (1:1000) and amplified using an anti-rabbit secondary antibody conjugated to AF594 (1:2000).

**LC-MS/MS based synaptic protein quantification and data analysis**

Cortical neurons were cultured in 6-well plates at a density of 1 x $10^6$ cells per well**.** Following shRNA mediated gene silencing using lentiviral transduction at DIV5-6 (three wells per protein target), cells were lysed at DIV15-16 in ice-cold 1x RIPA buffer supplemented with protease inhibitor (200µL per three wells). Cells were detached using a cell scraper and transferred into prechilled 2mL Eppendorf tubes, gently vortexed, and incubated on ice for 20 mins. Lysates were centrifuged at 13000g for 20 min at 4 °C, the resulting supernatants were collected and stored at -20 °C. The whole cell lysates (supernatants) from each strain were precipitated with cold acetone at 1:4 (*v:v*) ratio, -20 °C for overnight. Protein pellets were centrifuged and washed with cold acetone. Residue acetone was then air dried, and the proteins were re-dissolved using 0.25% RapiGest according to manufacturer's protocol. Afterwards, proteins were reduced with 5mM DTT (65°C, 30 min) and alkylated using 11mM IAA at ambient temperature in dark for 30 minutes. Trypsin was added at a protein-enzyme ratio of 50:1 for overnight digestion at 37 °C. The digestion was quenched by adding 10% TFA to pH between 1 and 2 and further incubated at 37 °C for 45 min. The solutions were then centrifuged at 14 000 rpm for 10 min, and the supernatant peptides were collected. Desalting of the peptides was performed using C18 ZipTip (Millipore, ZTC18M096) and the eluents were dried using SpeedVac (Thermo Scientific). The samples were stored at –80 °C before being re-suspended in 0.1% formic acid for LC-MS/MS analysis.

LC separation was performed on a Vanquish Neo System (Thermo Scientific) with an IonOptik Aurora Ultimate C18 column (15cm length × 75 µm inner diameter × 1.7 µm particle size) at 35 °C and 300 nL/min flow rate. Mobile phase A consisted of 0.1% formic acid in water. Mobile phase B consisted of 0.1% formic acid in 80% ACN. Eluting peptides were ionized by electrospray ionization and then analyzed by an Orbitrap Ascend Tribrid mass spectrometer (tune version 4.2.4321, Thermo Scientific). Ion transfer tube temperature was set to 275 °C. Source positive ion voltage was set to 1950v. For single-shot proteomics with data independent acquisition (DIA), the MS1 scan resolution was set to 120,000 (at m/z 200). MS1 scan range was 400–980 m/z, AGC target was 225%, maximum injection time mode was set to auto. Precursors were isolated with an isolation width of 12 m/z covering 145–1450 m/z. Precursors were fragmented by HCD at an NCE of 26%. MS2 scans were acquired by Orbitrap with a resolution of 15,000. Maximum injection time was 59 ms for the Ascend.

For acquired DIA data, raw data was directly processed using DIA-NN (1.9.2) with Mus musculus (Mouse, UP000000589) proteome fasta file. A maximum of one missed cleavage was allowed and cysteine carbamidomethylation (+57.0215 Da) as the fixed modification. Oxidation (M) and Acetylation (N-term) were selected as variable modifications. And precursor FDR was set to 1%. The mass accuracy and MS1 accuracy tolerance window were optimized by DIA-NN during the processing. Match-Between-Runs (MBR) was selected and the quantification matrix was generated after processing. All statistical analyses were performed using R (version 4.3) with the following packages: *tidyverse* for data manipulation; *readxl* and *writexl* for Excel file handling; *limma* for differential expression analysis; *pheatmap* and *ComplexHeatmap* for heatmap visualization; *ggplot2* with *ggrepel* for publication-quality plots; *clusterProfiler* and *enrichplot* for GO enrichment; *fgsea* and *msigdbr* for GSEA; *UpSetR* for set intersection visualization; and *dendextend* for dendrogram customization. Differential expression magnitudes were visualized using $Log_2$ Fold Change metrics relative to wild-type controls. All analysis scripts are provided as supplementary materials.

Raw protein abundance data were processed using the R statistical computing environment (v4.3). To account for missing values characteristic of label-free proteomics—typically representing peptides below the limit of detection—we applied a column-wise minimum imputation strategy. Specifically, missing values were replaced with 10% of the minimum observed intensity value within each respective sample column:

$$Impute_{Val} = 0.1 \times Min_{\text{sample}}$$

Following imputation, intensities were $Log_2$-transformed with a pseudo-count of 1 ($\log_2(x + 1)$) to stabilize variance and normalize the distribution for downstream statistical analysis.

*Quality Control and Exploratory Analysis*s

Data integrity and sample reproducibility were evaluated using a comprehensive multi-faceted approach:

*Correlation Analysis*

Sample similarity was assessed using Pearson correlation coefficients calculated on $Log_2$-transformed data. Correlation matrices were visualized as heatmaps using the pheatmap R package. Within-condition correlations were calculated to evaluate technical reproducibility among biological replicates.

*Dimensionality Reduction*

Principal Component Analysis (PCA) was performed on centered and scaled data to visualize global sample separation and identify potential batch effects. Variance explained by each principal component was calculated and visualized using scree plots. Multi-Dimensional Scaling (MDS) was conducted using Euclidean distances as a complementary approach to verify group clustering patterns.

*Knockdown Efficiency and Specificity Analysis*

Knockdown efficiency for target synaptic proteins was quantified by calculating the ratio of mean intensity in knockdown (KO) samples relative to wild-type (WT) controls. Percentage knockdown was defined as:

$$\% \text{ Knockdown} = 100 - (\text{Mean}_{\text{KO}} / \text{Mean}_{\text{WT}} \times 100)$$

**TEM sample preparation and imaging**

After activity imaging, imaging media (ACSF) was aspirated, and cells were rinsed twice with 0.1M phosphate buffer at RT. Next, 1mL of warm EM fixative (2% glutaraldehyde, 2mM $CaCl_2$ in 0.1M SC buffer pH 7.2) was gently added, and cells were incubated on ice for 1h. Fixative was then aspirated and neutralized with ice-cold glycine (20mM) in 0.1M SC buffer. Cells were thoroughly rinsed with ice-cold 0.1M SC buffer, followed by addition of 0.5mg/mL DAB and 10mM $H_2O_2$ in ice-cold 0.1M SC buffer. Cells were incubated on ice for 25 min and washed thoroughly with fresh ice-cold 0.1M SC containing 2mM $CaCl_2$. EM processing began with post-fixation in 2% $OsO_4$, 1.25% $K_4Fe(CN)_6$ and 3mM $CaCl_2$ in 0.15M sodium cacodylate for 30 min on ice. After rinsing with 0.1 M SC buffer containing 2mM $CaCl_2$, cells were incubated with 1% uranyl acetate in water as the positive stain for 2 days at 4 °C. Uranyl acetate solution was gently aspirated, and cells were rinsed three times with ice-cold milliQ water. Serial ethanol dehydrations were carried out using ice-cold solutions (30%, 50%, 70%, 90% ethanol; 7 min each). Cells were brought to RT in 90% ethanol and further dehydrated with three exchanges of 100% ethanol (7 min each) at RT.

Cells were infiltrated with Epon resin in ethanol at 1:2, 1:1 and 2:1 (v/v) ratios for 90 min each at RT. Cells were then incubated overnight in 100% Epon resin containing a reduced amount of accelerator. The next day, cells underwent three exchanges of 100% Epon resin with the manufacturer recommended accelerator amount, at 2h intervals. Excess Epon was removed from the dish, and coverslips were gently detached and placed cells-up onto an Aclar sheet (EMS, Catalog# 50426-10) supported by a heat-resistant base. A labeled BEEM capsule (Ted Pella Product #130) filled with Epon resin was very carefully inverted over the region of interest (ROI) on the coverslip. Polymerization was proceeded at 60 °C for 48h. Post polymerization, coverslips were separated from the BEEM capsule by alternating brief submersion in liquid nitrogen and hot water. Grid markings etched onto the block face facilitated locating the region and neurons of interest. Blocks were trimmed and serially sectioned (50nm) using a Leica UC6 ultramicrotome, with thin sections collected on formvar coated slot grids. The ROIs were mapped to the activity patterns via low-magnification imaging (80keV) on a Tecnai G2 Spirit Biotwin transmission electron microscope using a Gatan Oneview camera (4k x 4k resolution). Higher-magnification TEM images were acquired for detailed analysis. Automatic detection and quantification of synaptic vesicle parameters were performed in Python 3.12 using a published algorithm available at: https://github.com/Imbrosci/synaptic-vesicles-detection. Small fields of views (FOVs) that include DAB+ and DAB- boutons (8-bit TIFF images) were analyzed, and statistical tests were performed in Prism10 (GraphPad software).

**Expansion microscopy (ExM) sample preparation and data analysis**

Tissue preparation and gel recipe for ExM were performed according to a previously published procedure (*8*). Briefly, ALFADoR1 and ALFADoR2 mice were anesthetized with isoflurane and perfused transcardially with a ice-cold solution of 4% PFA and 10% acrylamide in 1x PBS. After overnight post-fixation in the same fixative, brains were sectioned coronally into 100µm slices and excess PFA was quenched with 100mM glycine. Tissue sections containing prefrontal cortex, striatum and amygdala were selected for the study. After the first round of polymerization, tissue-gel pieces were probed using rabbit anti-TH and human anti-ALFA antibodies at 1:300 dilutions at 4 °C overnight with gentle shaking. Donkey anti-rabbit CF633 and goat anti-human AF546 secondary antibodies were used at 1:300 dilutions and probing were carried out overnight at 4 °C. ExM images were collected on Nikon spinning disk microscope, using a 40x water immersion objective (1.25NA).

Typical imaging volumes included fields of view that were approximately 375 μm by 375 μm in x-y plan, and 50–100 μm in the z-direction, taken at 0.5 μm z-stacks. Each field of view contained two channels: a dopamine axon channel (dopamine axons with varicosities) and an ALFA-tagged receptor channel (D1-ALFA *or* D2-ALFA). Images were analyzed in MATLAB using a custom pipeline designed to quantify nanoscale apposition between dopamine varicosities and receptor clusters in 2D. Images were converted to double precision and normalized. A light Gaussian smoothing step was applied to stabilize segmentation, using 0.5 sigma in pixels. The dopamine and receptor channels were processed independently up to the point of apposition analysis.

Dopamine axons were first segmented to define an axonal domain that constrained subsequent varicosity identification. The dopamine channel was enhanced to emphasize elongated axon-like structures and then thresholded to generate an initial binary axon mask. Morphological cleanup steps were applied to close small gaps, fill holes, and remove small, isolated components. Varicosities were then segmented using a watershed-based approach, which improved granularity and avoided merged varicosity objects in densely labeled axons when compared to other segmentation methods we tried. Within the axon mask, the dopamine channel was smoothed and thresholded to identify bright candidate regions. To separate adjacent bright structures, watershed segmentation was applied to a topography derived from the intensity image such that local intensity peaks were resolved into distinct objects. The resulting objects were filtered by minimum size, producing a final varicosity mask. For each segmented varicosity, geometric descriptors including centroid position, area, and perimeter were extracted using connected-component labeling and region properties. These per-varicosity measurements were used for both spatial apposition quantification and downstream distributional analyses. Receptor clusters were segmented from the ALFA channel using a complementary watershed-based puncta pipeline. The receptor image was smoothed, a candidate bright mask was generated by thresholding, and watershed was applied to separate adjacent puncta. An h-minima suppression step was used to reduce over-segmentation by removing shallow intensity minima prior to watershed. Segmented puncta were then filtered by a minimum area threshold. To control stringency and suppress dim detections, puncta were optionally filtered by requiring that each object's peak intensity exceed a user-defined fraction of the maximum image intensity. For each punctum, centroid position and morphological features (including area, major axis length, minor axis length, eccentricity, and perimeter) were quantified.

Apposition between dopamine varicosities and receptor puncta was quantified using a perimeter-based minimum distance criterion designed to reflect near-contact between the boundaries of segmented structures. In our analysis, we defined apposition as proximity of less than 0.125 μm, which corresponded to 3 pixels in the expanded gel images, but additional appositional analysis was performed at larger threshold distances of 0.25, 0.35 and 0.5 μm, corresponding to camera pixels of 6, 9 and 13 in expanded gels, as shown in Fig. 7G. A distance transform was computed from the receptor puncta mask to the nearest receptor pixel. For each varicosity, the analysis extracted the varicosity perimeter and computed the minimum distance from any perimeter pixel to the receptor mask. Varicosities were classified as “apposed” at a given threshold if this minimum distance was less than or equal to the threshold (in pixels). In addition to binary classification, a fractional contact metric was computed for each varicosity at each threshold, defined as the fraction of varicosity perimeter pixels lying within the threshold distance to the receptor mask. These per-object values were aggregated across varicosities to generate apposition fractions per imaging fields of view.

To determine whether the observed apposition exceeded what would be expected from receptor density and spatial heterogeneity alone (i.e., appositional encounters by chance-alone), we implemented a conservative null model based on toroidal shuffling. For each permutation, the receptor puncta mask was translated by a uniformly random (x, y) offset with wrap-around at image boundaries. Shifting (x,y) values were randomly chosen such that $0 < x_{shift} <$ ImagePixelSize in the x-direction, and the same approach was used for shifts in the y-direction. This procedure preserves the number, size, and internal geometry (for example, noise, autocorrelation) of puncta, as well as any broad spatial inhomogeneities in receptor labeling, while disrupting spatial alignment to dopamine axons and varicosities. For each shuffled run, the same varicosity-to-receptor perimeter distance analysis was repeated and the resulting apposition metrics computed and stored. Null means and confidence intervals were computed from the permutation results, and p-values were reported by comparing the experimental values to the shuffled distribution. These shuffled controls were used throughout to benchmark “apposition by chance” expectations and to support our hypothesis that proximity reflects structured coupling rather than random overlap.

Several complementary spatial summaries were computed from the same segmented objects. Nearest-neighbor distance distributions were evaluated by measuring, for each varicosity centroid, the distance to the nearest receptor punctum centroid, and reciprocally, for each receptor punctum centroid, the distance to the nearest varicosity centroid. These distances were used to generate cumulative distribution functions (CDFs) for each field of view and to compare distributions across brain regions and receptor types (as shown in Fig. 7M and 7N).

### Supplementary Figures

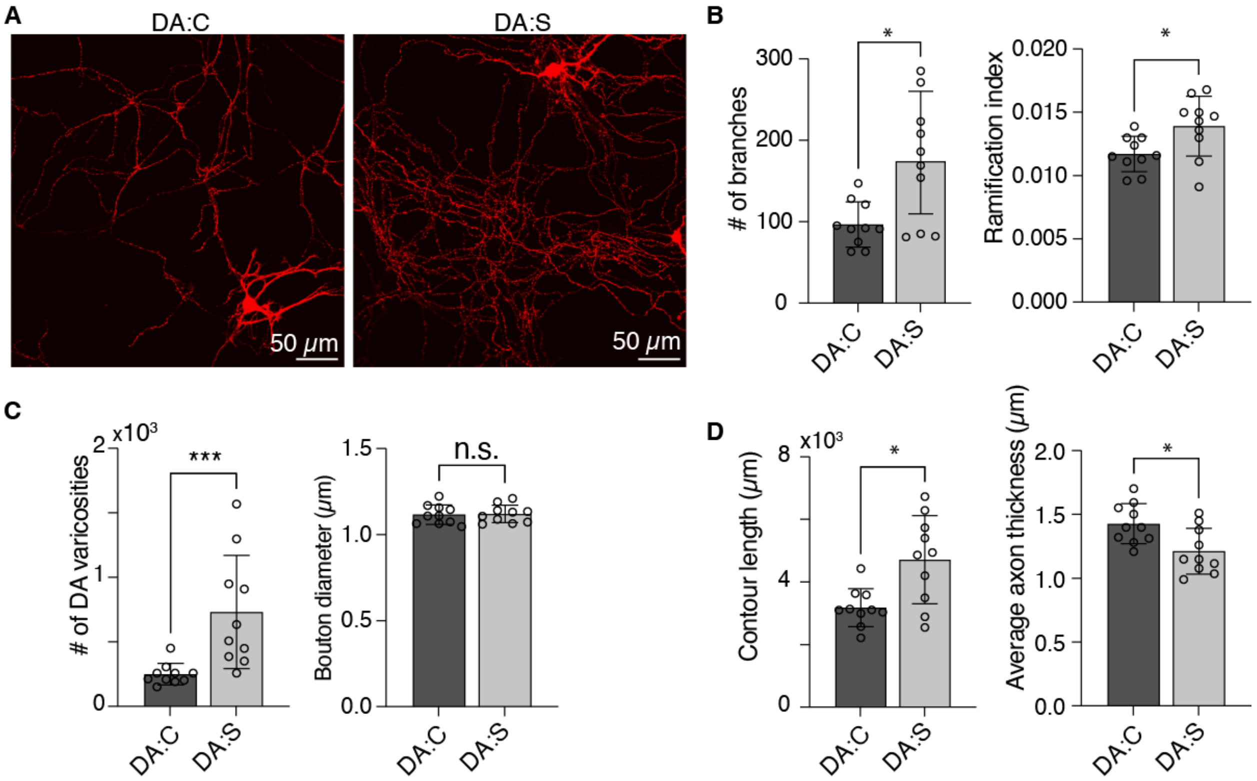


**Figure S1. DA:C and DA:S cultures exhibit morphological differences that mirror reported in vivo characteristics**

(A) Representative dopamine axonal arborization in an imaging field of view corresponding to DA:C and DA:S co-cultures. Note that DA:S cultures generate highly ramified dopaminergic axonal processes.

(B–D) Analysis of arborization (B), varicosity (C) and axonal morphometrics (D). Each data point represents average across an independent FOV belonging to one dopamine neuron. Mean ± SD bars are shown. Test of statistical significance: Mann-Whitney test. p-values: n.s. = not significant, * < 0.05, *** < $10^{-3}$. Ramification index = # of branches per unit axon length.

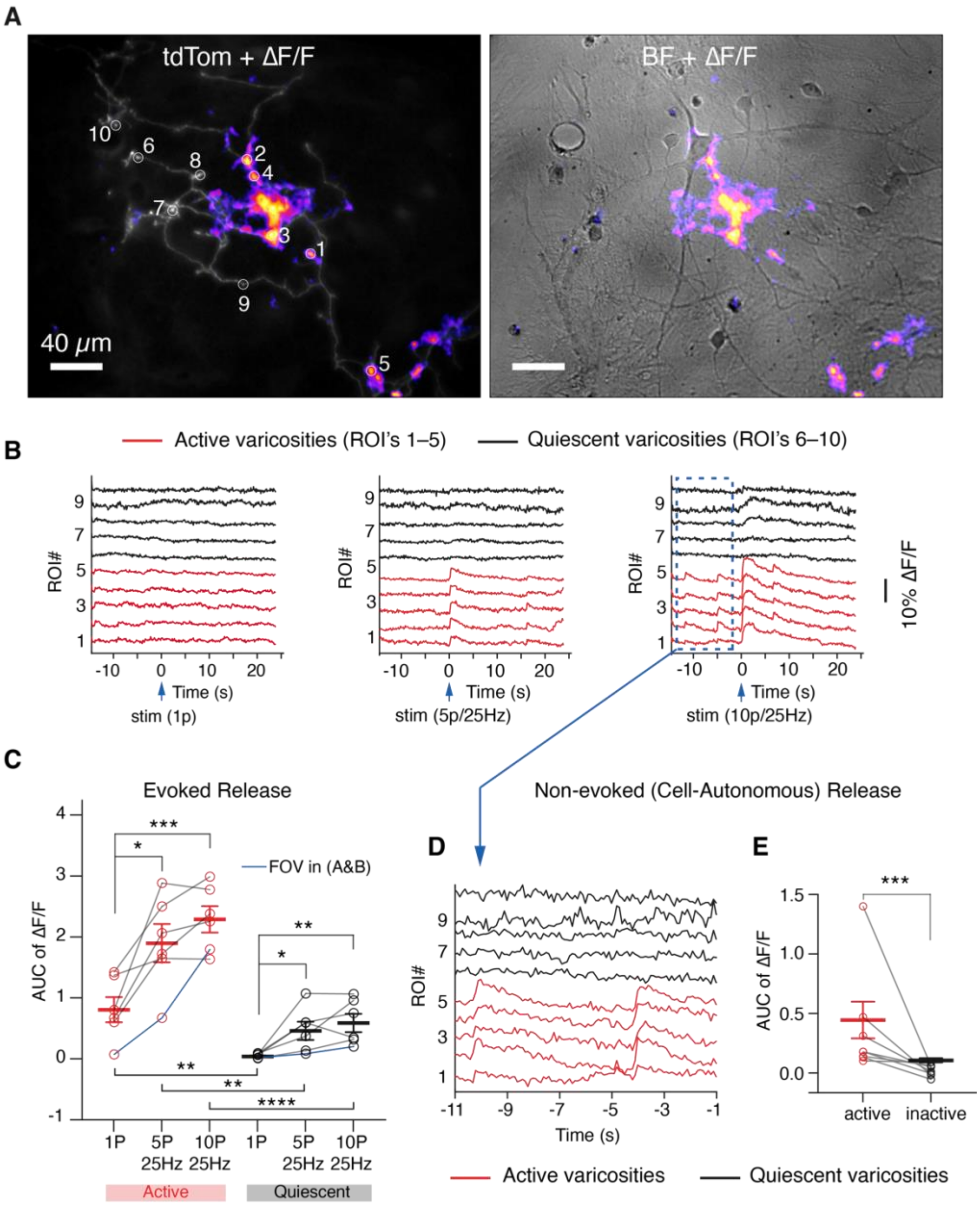


**Figure S2. Characterization of active and quiescent varicosities.**

(A) An imaging field of view showing dopamine axons (tdTom) overlaid with dopamine release activity heatmap (ΔF/F). Right: brightfield (BF) image of the same FOV shown on the left.

(B) Activity traces corresponding to the varicosities (ROI's 1–10) depicted in (A) corresponding to single pulse (left), 5-pulse (middle) and 10-pulse (right) electrical field stimulation.

(C) Analysis of area under the curve (AUC) of ΔF/F trace over a 3-second window after stimulation. Each data is an average across a field of view of imaging from n = 10 randomly sampled active and quiescent varicosities; mean±SEM bars are shown. The blue trace represents data from the field of view/data shown in (A&B). Test of statistical significance: Kolmogorov-Smirnov (KS) test. p-values: * < 0.05, ** < $10^{-2}$, *** < $10^{-3}$, **** < $10^{-4}$

(D) Close-up of non-evoked release window corresponding to the 10-pulse stimulation event shown in (B). Note that quiescent varicosities remain release-incompetent during non-evoked release events.

(E) Analysis of AUC of non-evoked release events such as that shown in (D). Each point is an average from an independent field of view, and mean ± SEM bars are shown. Test of statistical significance: KS test. p-value: *** < $10^{-3}$

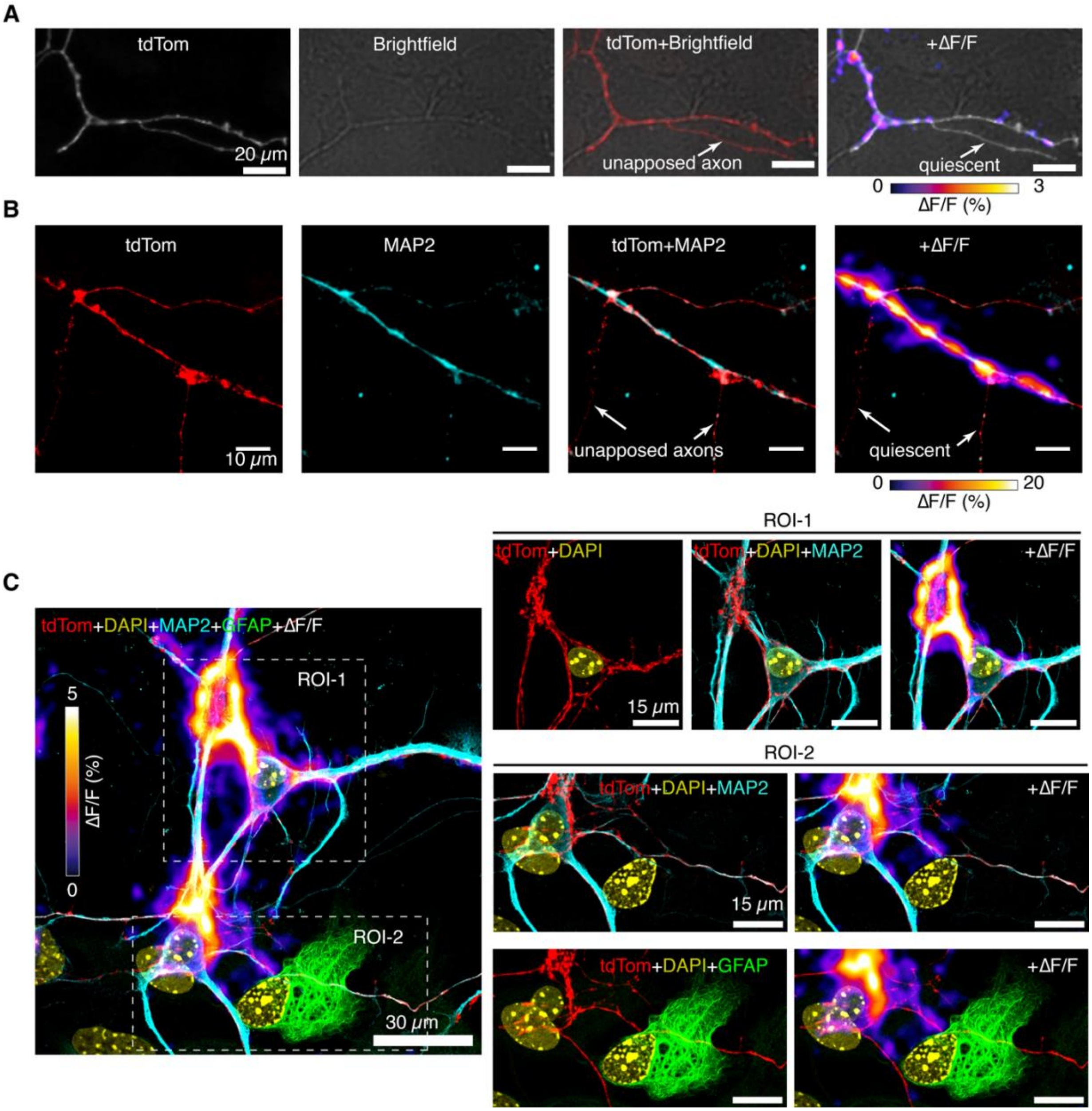


**Figure S3. Apposition to neuronal processes is necessary for release-competence.**

(A) Live-cell imaging: a dopamine axonal process (tdTom) and corresponding bright field image, overlay between tdTom and brightfield, and overlay with dopamine release (ΔF/F) are shown. Notice the process in brightfield with which the dopamine axon is associating/co-mingling. A non-apposed axonal branch does not participate in release.

(B) Immunocytochemistry staining for MAP2: only dopamine axons (tdTom) that are dendritically apposed participate in release. Notice that non-apposed axonal branches do not participate in release.

(C) Immunocytochemistry staining for MAP2 and GFAP: MAP2 apposed axonal processes participate in release whereas GFAP-associating axons remain quiescent.

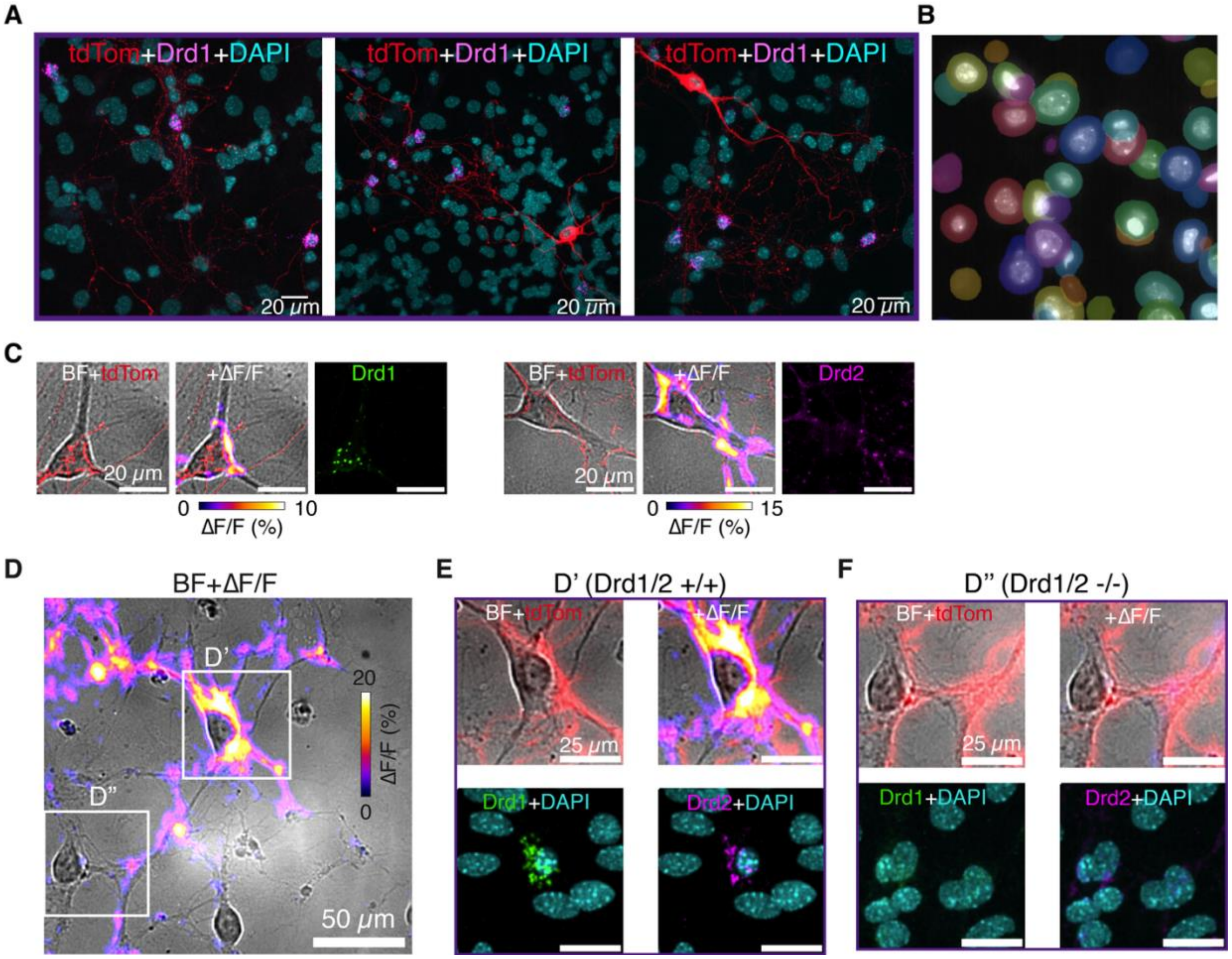


**Figure S4. Transcriptomic profiling of dopamine receptors in culture and tissue shows tropism of dopamine axons towards Drd1 or Drd2-positive nuclei.**

(A) Three example fields of view showing dopamine axons (red, tdTom) preferentially innervating DAPI nuclei (blue) that are positive for Drd1 (magenta).

(B) An example of a FOV containing Cellpose-segmented DAPI-nuclei used in analysis of tropism in dopamine axons.

(C) Example Drd1 and Drd2 soma in brightfield, overlayed with Drd1 (green puncta) and Drd2 (magenta puncta) readout from RNAScope. Dopamine release heatmap shows secretion of dopamine onto soma.

(D) A FOV of imaging showing brightfield image (BF) and dopamine release (ΔF/F).

(E, F) Close up of D' and D'' (from panel D) showing tdTom, ΔF/F heatmaps, DAPI stain and readout from RNAScope. Neuron in D' is positive for both Drd1 and Drd2 whereas the neuron in D'' is negative for both. Note the absence of dopamine release in F'' despite some innervation by dopaminergic axons.

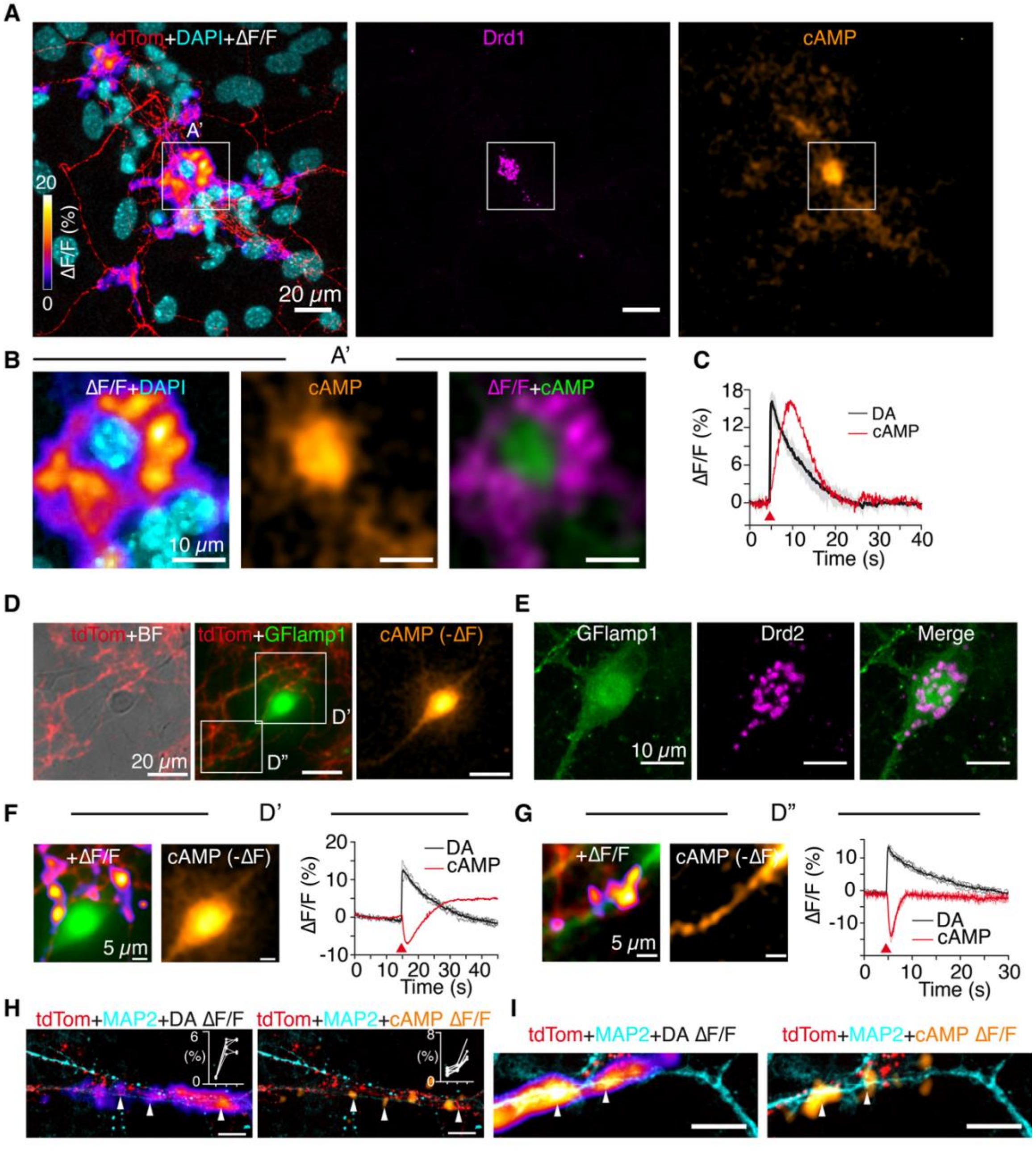


**Figure S5. Dopamine release rapidly modulates cAMP levels in target neurons.**

(A) A three-panel figure showing tdTom, DAPI and ΔF/F, Drd1 RNA transcripts probed by RNAScope, and cAMP ΔF/F frame. Dopamine and cAMP peak ΔF/F images are shown, which occur at different time points after application of stimulus (see panel C). Note that this field of view is identical to one used in Fig. 3A–C.

(B) Magnified view of A' depicted in the solid white box in (A). Overlaid images of ΔF/F and DAPI, and cAMP ΔF frame (orange), and a merge of dopamine ΔF/F (magenta) and cAMP ΔF (green) are shown. An extracellular dopamine “corona” elicits a vigorous intercellular cAMP activity.

(C) Temporal traces of dopamine (black) and cAMP (red) for images shown in (B) are shown. Dopamine is presented as an average trace of identified hotspots and cAMP activity is averaged over the soma.

(D–E) A representative example of a dopaminergic axonal innervation of a cortical neuron exhibiting negative cAMP modulation at the soma and dendritic shafts. The cAMP pseudo color is inverted for display purposes. The neuron was confirmed to be positive for Drd2 transcripts using post hoc RNA FISH as shown in (E).

(F–G) Zoomed-in images of white box ROIs depicted in (D), along with temporal traces showing dopamine and cAMP activity.

(H–I) cAMP activity hotpots generated by dopamine hotspots in distal dendritic processes in DA:C co-cultures. Note that dopamine activity and cAMP activity are spatially co-located. Insets in (H): DA and cAMP peak ΔF/F scale with stimulation (2 pulses, 5 pulses and 10 pulses from left to right, all at 25 Hz).

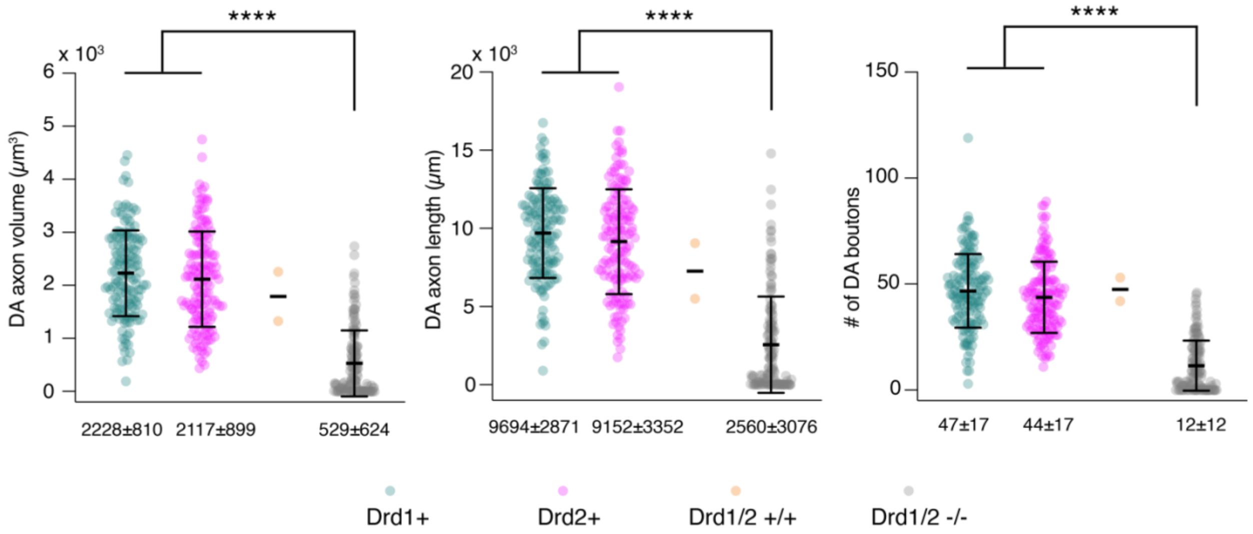


**Figure S6. Analysis of EASI-FISH volume in the dorsal striatum.**

Total axon volume, length and number of boutons are evaluated. Each point (n = 150 for three groups) is a DAPI+ nucleus and dopamine axons that associate with the soma (approximated by the nucleus with some dilation, see Methods) are segmented and analyzed. Mean ± SD bars and values are shown at the bottom. A two-sample KS-test was used to evaluate significance of differences between transcript-positive vs. negative groups (group-wise); p-value: **** < $10^{-4}$. See Methods for more on analysis of EASI-FISH volume.

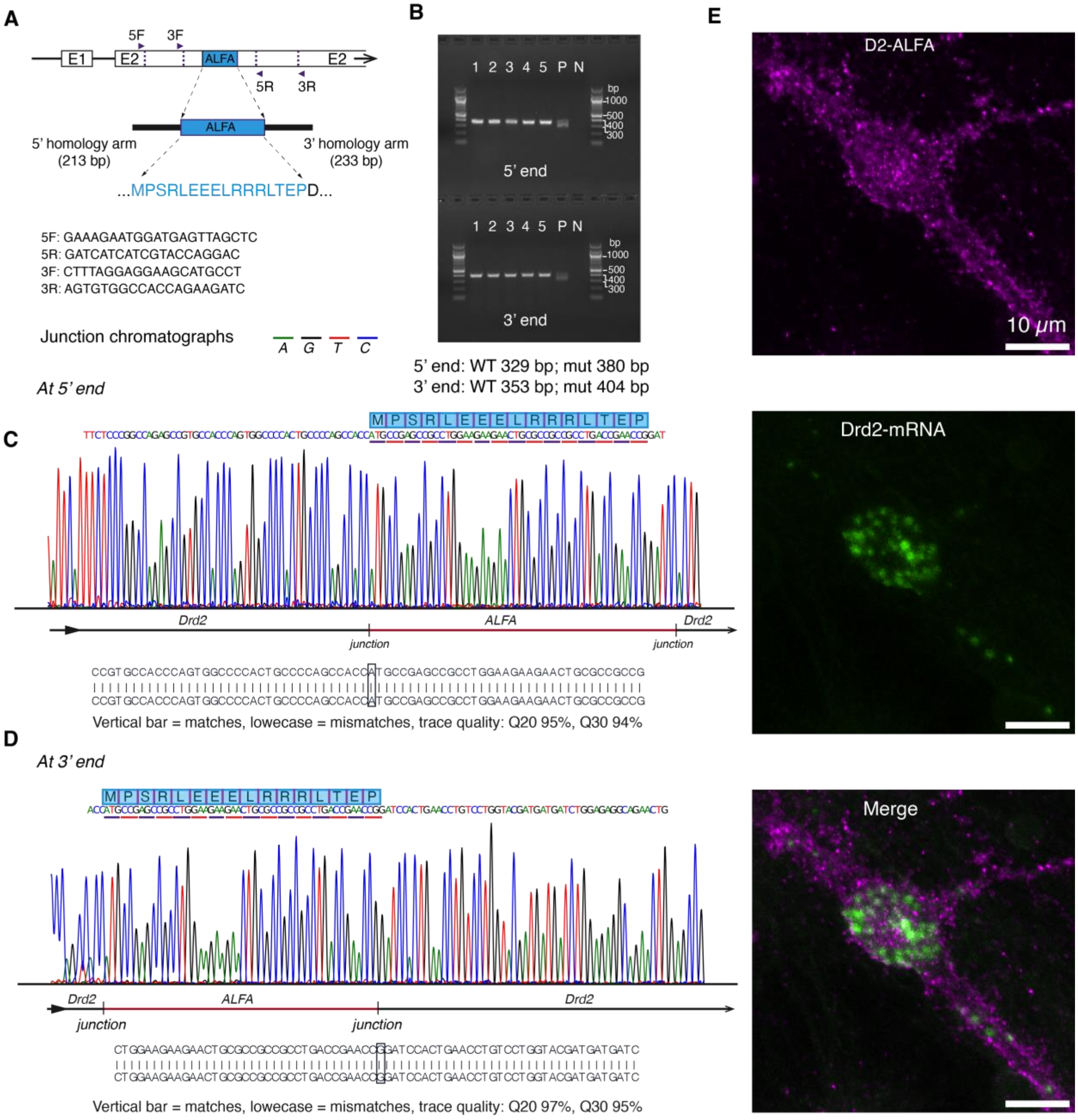


**Figure S7. Design and validation of ALFA-tag knock in into the Drd2 gene locus.**

(A) The design schematic of ALFA-tag insertion into the second exon (E2) of Drd2 gene, with 213 bp and 233 bp homology arms on the 5' and 3' ends respectively. 5' and 3' genotyping primers are shown.

(B) Gel electrophoresis of PCR products confirming correct integration at both the 5′ and 3′ junctions. Five biological replicates are shown. The positive control (P), containing a mixture of wild-type (WT) and knock-in alleles (mut), displays two closely migrating bands corresponding to the WT and mut products. The negative control (N) lane, lacking template DNA, shows no detectable amplification, confirming specificity and absence of contamination.

(C–D) Representative Sanger sequencing chromatograms spanning the 5′ and 3' junctions of the ALFA-tag knock-in at the Drd2 gene locus. Traces show high-quality, well-resolved base calls across the genomic sequence upstream of the insertion site, the ALFA-tag coding sequence, and the downstream genomic sequence. Individual base calls are aligned below the chromatograms, with vertical bars indicating exact matches to the reference sequence. No insertions, deletions, or sequence ambiguities were detected at the junctions. Phred quality metrics are shown for the displayed window (Q-scores) confirming high-confidence base calling.

(E) Representative RNAScope readout of Drd2 mRNA transcripts shows ALFA-tag expression is observed in neurons that are enriched in Drd2 mRNA puncta. Drd2 mRNA puncta were detected in n=34 of 35 cells that expressed the ALFA-tag.

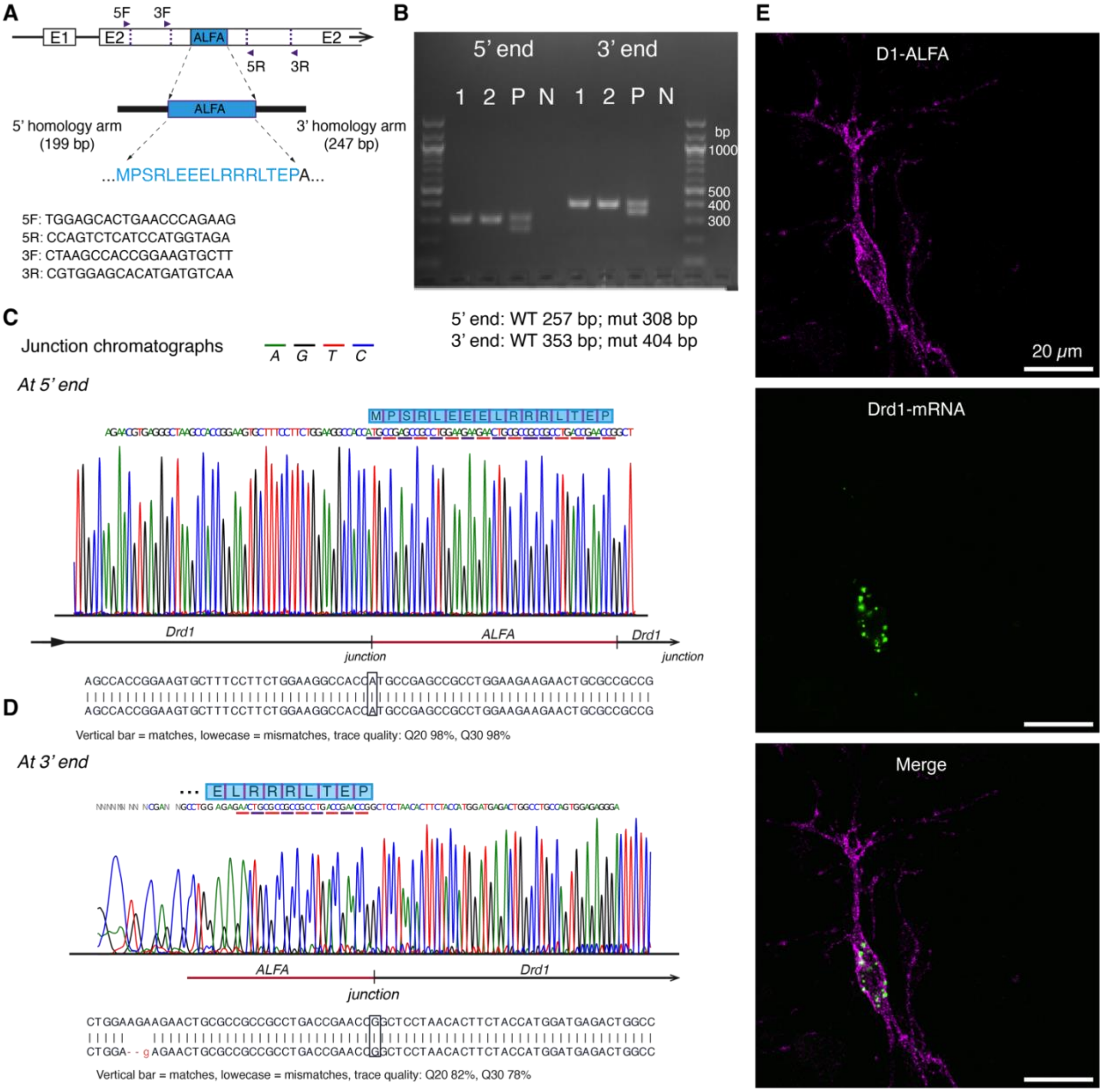


**Figure S8. Design and validation of ALFA-tag knock in into the Drd1 gene locus.**

(A) The design schematic of ALFA-tag insertion into the second exon (E2) of Drd1 gene, with 199 bp and 247 bp homology arms on the 5' and 3' ends respectively. 5' and 3' genotyping primers are shown.

(B) Gel electrophoresis of PCR products confirming correct integration at both the 5′ and 3′ junctions. Two biological replicates are shown. The positive control (P), containing a mixture of wild-type (WT) and knock-in alleles (mut), displays two closely migrating bands corresponding to the WT and mut products. The negative control (N) lane, lacking template DNA, shows no detectable amplification, confirming specificity and absence of contamination.

(C–D) Representative Sanger sequencing chromatograms spanning the 5′ and 3' junctions of the ALFA-tag knock-in at the *Drd1* gene locus. Traces show high-quality, well-resolved base calls across the genomic sequence upstream of the insertion site, the ALFA-tag coding sequence, and the downstream genomic sequence. Individual base calls are aligned below the chromatograms, with vertical bars indicating exact matches to the reference sequence. No insertions, deletions, or sequence ambiguities were detected at the junctions. Phred quality metrics are shown for the displayed window (Q-scores) confirming high-confidence base calling. Low read quality upstream of the ALFA sequence in (D) is compensated by high quality reads on upstream of the ALFA-tag sequence in (C).

(E) Representative RNAScope readout of Drd1 mRNA transcripts shows ALFA-tag expression is observed in neurons that are enriched in Drd1 mRNA puncta. Drd1 mRNA puncta were detected in n = 5 of 5 cells that expressed the ALFA-tag.

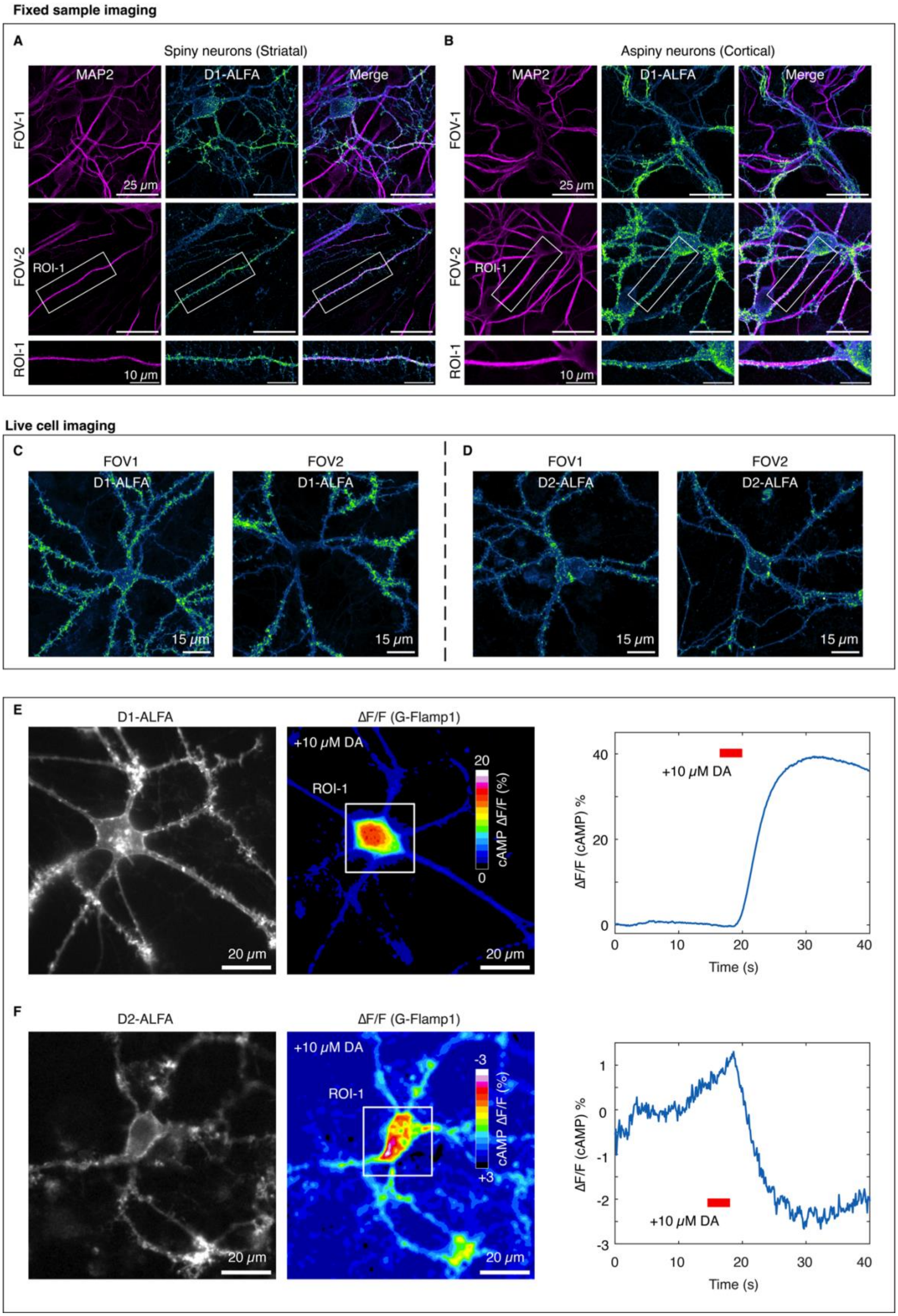
Fixed sample imaging
A
Spiny neurons (Striatal)
MAP2
D1-ALFA
Merge
FOV-1
FOV-2
ROI-1
25 μm
10 μm
B
Aspiny neurons (Cortical)
Live cell imaging
C
FOV1
FOV2
D1-ALFA
15 μm
D
D2-ALFA
E
ΔF/F (G-Flamp1)
+10 μM DA
cAMP ΔF/F (%)
20 μm
ΔF/F (cAMP) %
Time (s)
F

**Figure S9. Visualizing D1-ALFA and D2-ALFA expression in fixed samples and live cells.**

(A–B) D1-ALFA expression in spiny and aspiny fixed neurons, co-labeled with MAP2 to visualize soma and dendrites. D1-ALFA is present in soma, dendritic shafts and spines. Expression shows pronounced local non-homogeneity, appearing as a punctate distribution, as well as global inhomogeneity, with some dendritic segments exhibiting higher D1-ALFA levels than others.

(C–D) Maximum intensity projections of live-cell images of D1-ALFA and D2-ALFA expression using an anti-ALFA nanobody conjugated to Atto643 reveals punctate receptor localization, with differential enrichment in specific neuronal compartments (two representative examples are shown for D1-ALFA and D2-ALFA).

(E–F) Bath application of 10 μM dopamine (DA) to a G-Flamp1–expressing Drd1+ and Drd2+ neurons. Live-cell receptor imaging and cAMP activity imaging are shown. Note the non-uniform segregation of local cAMP modulations. Right side panels represent temporal traces of cAMP modulations.

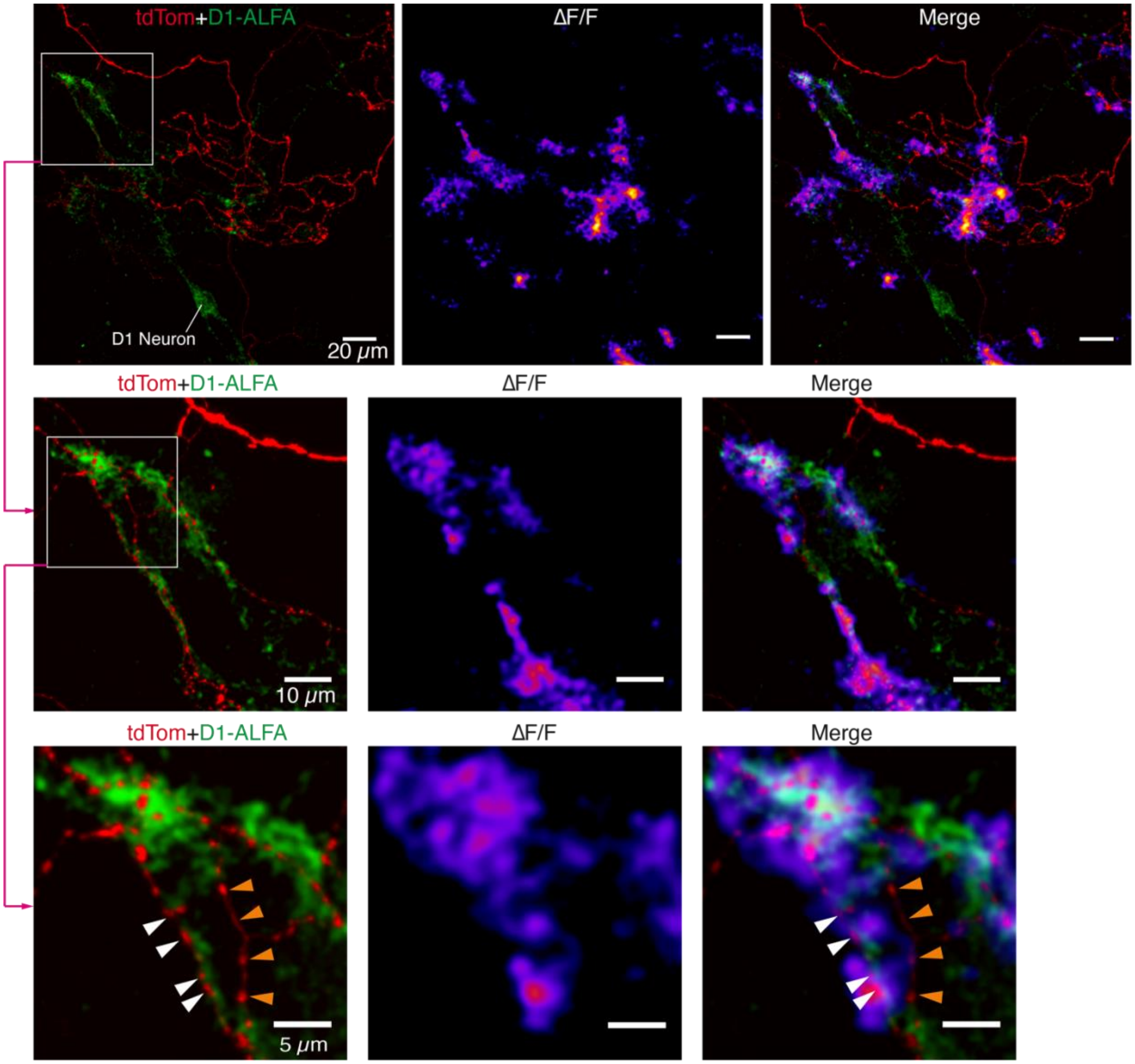


**Figure S10. DopaFilm activity imaging in a dopaminergic axonal arbor arborizing in the vicinity of a D1-ALFA–tag expressing neuron.**

DopaFilm signals are shown in a dopaminergic axonal arbor (tdTom, red) arborizing in the vicinity of a D1-ALFA–expressing neuron (green). Each row presents a progressively magnified view of a smaller region, as indicated by arrows and white boxes. Dopaminergic processes apposed to D1-ALFA–positive structures exhibit robust dopamine release, whereas non-apposed processes remain largely quiescent. Last row: white arrows indicate active varicosities, whereas orange arrows show quiescent ones.

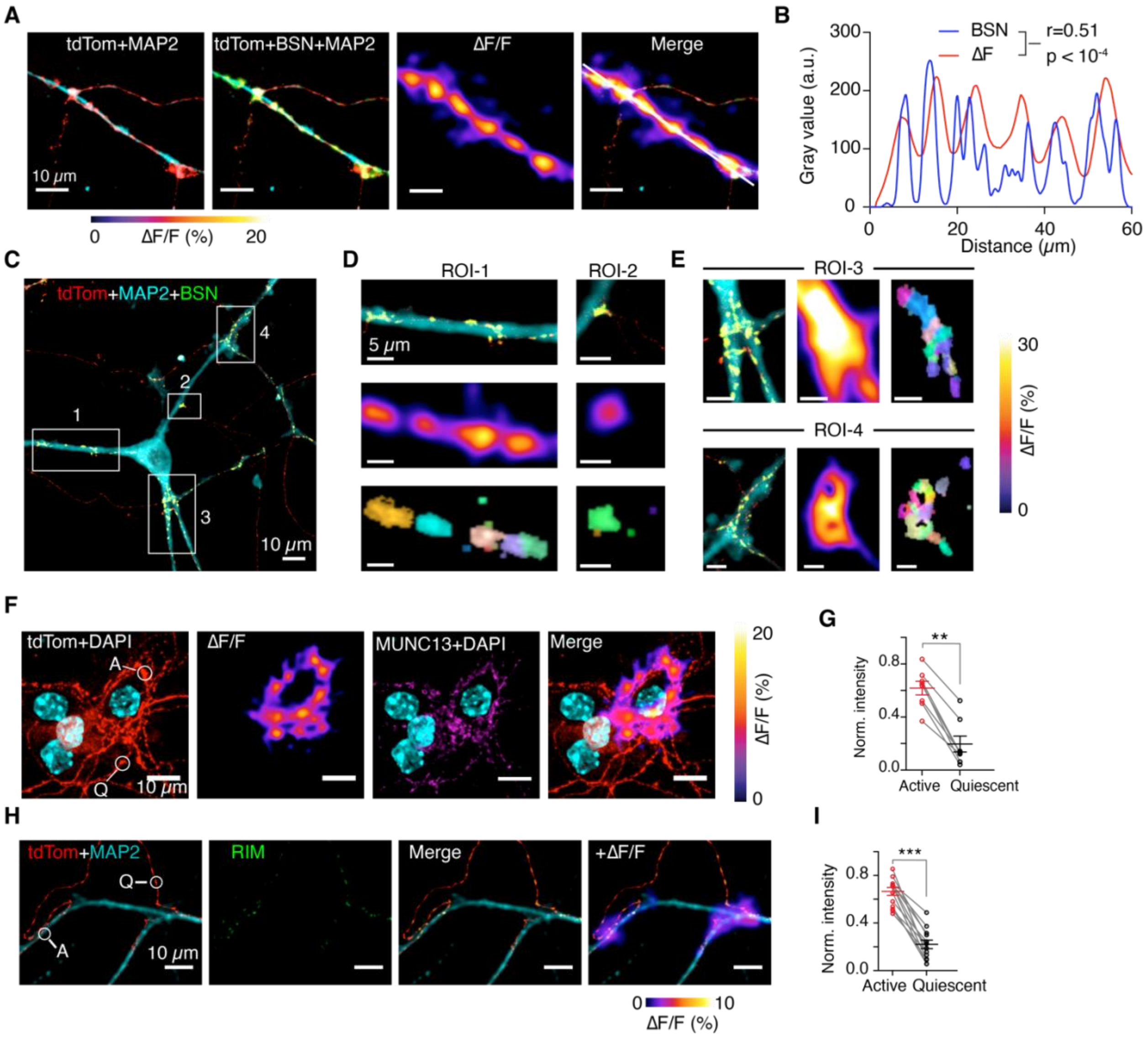


**Figure S11. Synapse-like molecular enrichment at active dopaminergic varicosities.**

(A) Dopaminergic axons apposed to dendrites (tdTomato, red; MAP2, cyan) exhibit robust dopamine hotspot formation (ΔF/F), whereas non-apposed axons remain largely quiescent. Bassoon (BSN; green) is enriched at active varicosities, appearing as yellow where BSN overlaps with tdTomato. In contrast, quiescent varicosities show low BSN levels and appear predominantly red.

(B) Line profile of dopamine release (ΔF/F) and BSN expression demonstrates a strong correlation (Pearson correlation r = 0.51). The line used for the profile is indicated in the rightmost panel of (A).

(C) MAP2-stained cortical neuron (cyan) innervated by dopaminergic axons (tdTomato, red). BSN puncta are visible as yellow overlap.

(D) Magnified views of ROI1 and ROI2 (white boxes in C). BSN-enriched active varicosities are distributed along the dendritic process. Bottom panels show NNMF components, which group highly correlated pixels into distinct color-coded clusters, each corresponding to an individual dopamine hotspot detected by DopaFilm.

(E) Magnified views of ROI3 and ROI4 (white boxes in C). Multiple Bsn puncta are closely clustered. These “bunched” puncta give rise to clustered release sites and appear as merged signals (as ‘blobs’ instead of distinct hotspots) in DopaFilm; however, NNMF decomposition resolves spatially distinct components corresponding to independent putative release sites (color codes, right panels).

(F–I) Enrichment of Munc13-1 and RIM at soma- or dendrite-apposed varicosities. Representative active (A) and quiescent (Q) varicosities are shown. Statistical significance was assessed using a Kolmogorov–Smirnov (KS) test (** $p < 10^{-2}$; *** $p < 10^{-3}$). Each point is an average over a biological replicate (typically a FOV containing hundreds of active and inactive varicosities). Mean and SEM bars are shown.

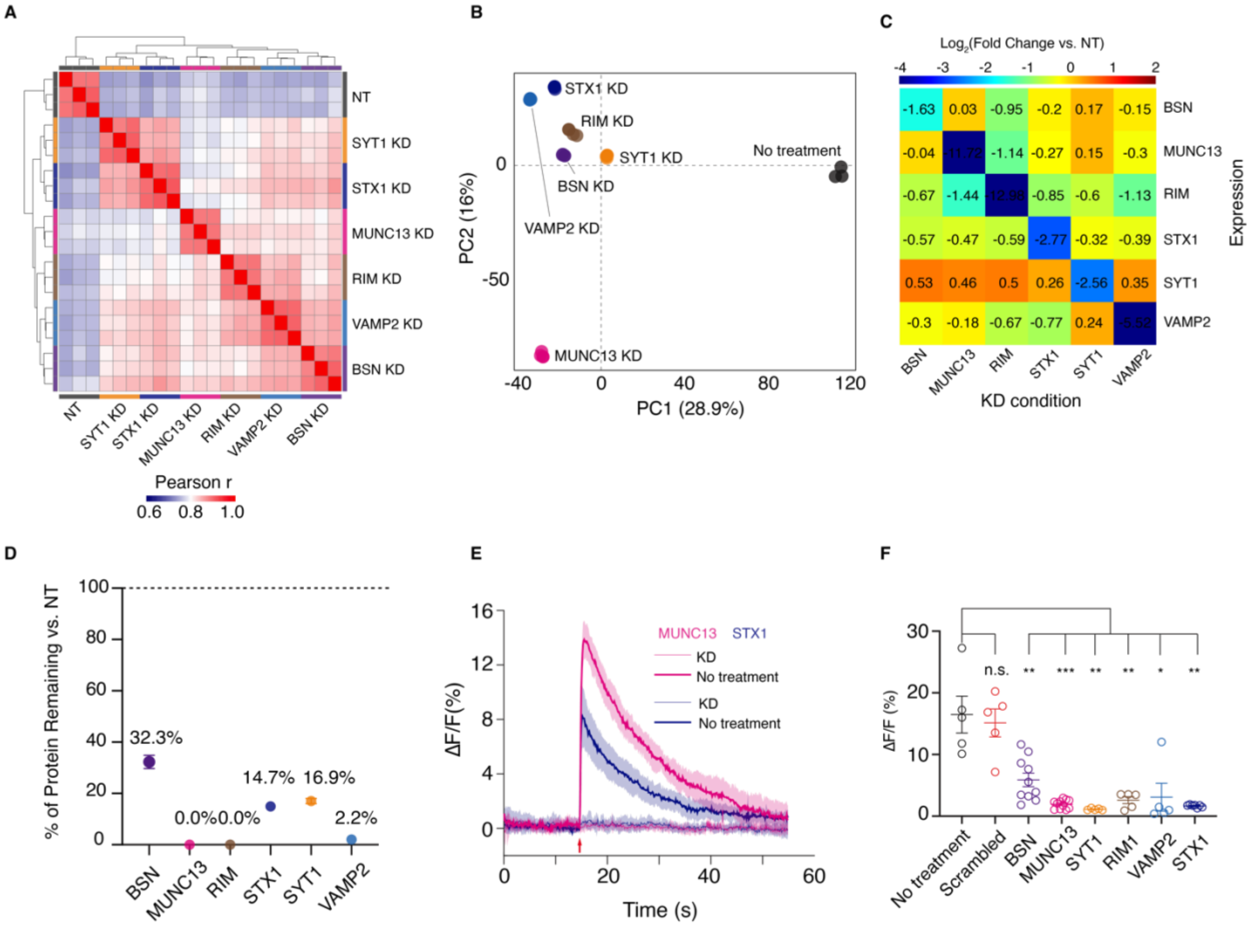

**Figure S12. Assessment of knockdown (KD) efficacy using quantitative proteomics.**

(A) Correlation heatmap (Pearson’s *r*) across six KD conditions and no-treatment (NT) control demonstrates highly reproducible technical replicates (n = 3 per condition) and clear separation between experimental groups.

(B) Principal component analysis (PCA) shows tight clustering within each treatment group and strong spatial separation between the NT control and KD conditions, indicating robust proteomic differences following knockdown.

(C) Fold-change heatmap (KD conditions relative to NT) reveals efficient target protein depletion following treatment, while also highlighting cross-depletion relationships among different KD conditions.

(D) Log2 fold-change values were converted to percent protein remaining, illustrating near-complete depletion for Munc13-1 and RIM knockdowns and overall strong depletion efficiency across most KD conditions.

(E) Representative dopamine activity imaging traces from a KD condition and an NT control. Munc13-1 and Stx1 are shown as an example.

(F) Quantification of peak ΔF/F responses for NT controls, scrambled controls, and KD treatments. Each point represents averaged ΔF/F activity from a biological replicate. Statistical significance was assessed using a Kolmogorov–Smirnov (KS) test (n.s., not significant; p-values: * < 0.05; ** < $10^{-2}$; *** < $10^{-3}$). Mean ± SEM bars are shown.

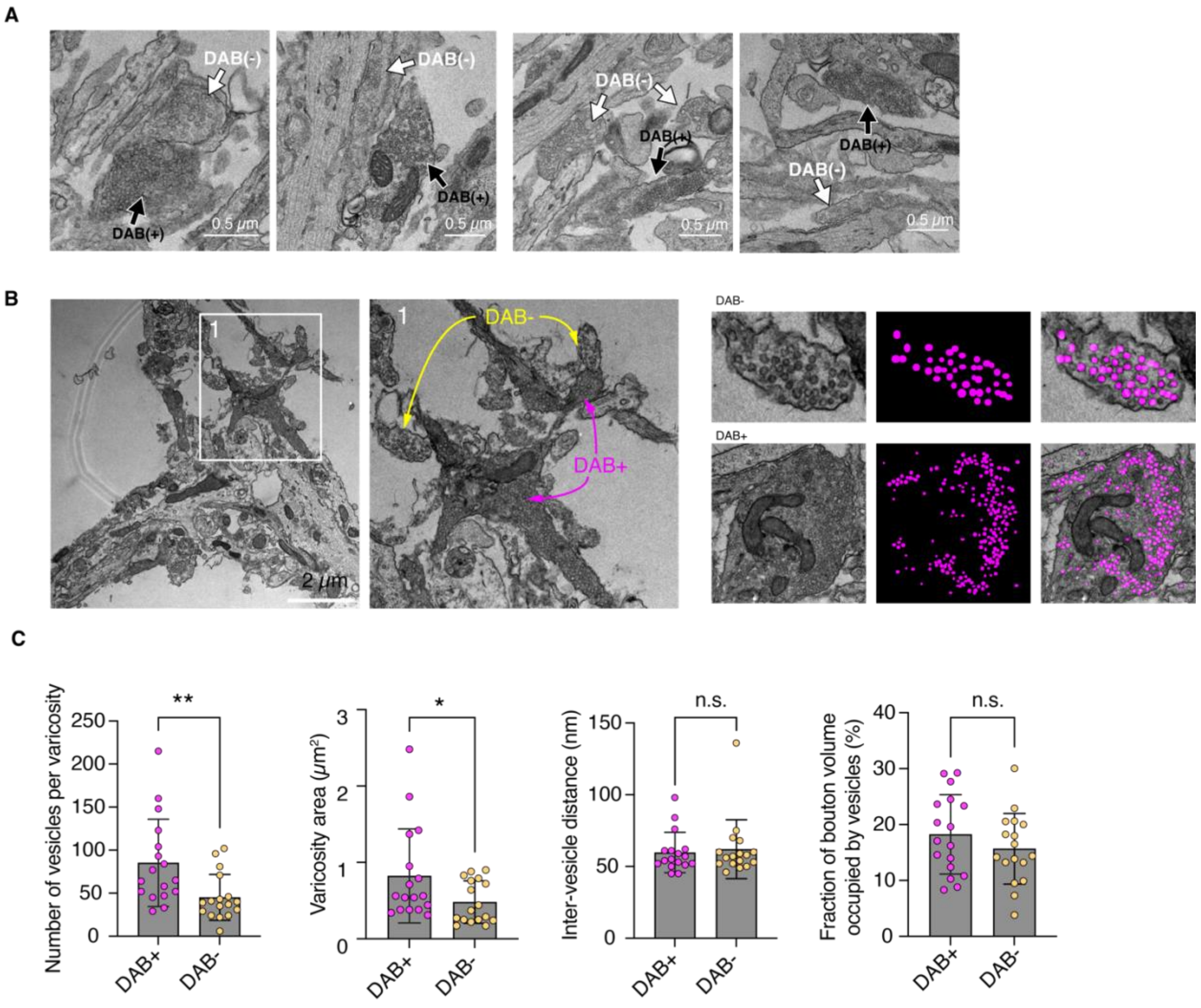


**Figure S13. APEX2-mediated DAB deposition for ultrastructural analysis of dopaminergic (DAB+) and non-dopaminergic varicosities (DAB-).**

(A) Representative transmission electron microscopy (TEM) thin sections showing DAB-positive (DAB+) and DAB-negative (DAB-) varicosities.

(B) A thin section containing DAB+ and DAB- varicosities. ROI1 (white box) is magnified to show DAB+ and DAB- varicosities within the same field of view. Right panels: DAB+ and DAB- varicosities, synaptic vesicle masks (magenta) and overlaid images are presented.

(C) Quantitative comparison of vesicle density and vesicle morphology between DAB+ and DAB- varicosities. Statistical significance was assessed using a Kolmogorov–Smirnov (KS) test (n.s., not significant; p-values: $* < 0.05$; $** < 10^{-2}$). Mean ± SD are shown.

### References


1. A. G. Beyene *et al.*, Imaging striatal dopamine release using a nongenetically encoded near infrared fluorescent catecholamine nanosensor. *Science Advances* **5**, eaaw3108 (2019).
2. C. Bulumulla *et al.*, Visualizing synaptic dopamine efflux with a 2D composite nanofilm. *eLife* **11**, e78773 (2022).
3. C. M. Bäckman *et al.*, Characterization of a mouse strain expressing Cre recombinase from the 3' untranslated region of the dopamine transporter locus. *Genesis* **44**, 383-390 (2006).
4. X. Zhuang, J. Masson, J. A. Gingrich, S. Rayport, R. Hen, Targeted gene expression in dopamine and serotonin neurons of the mouse brain. *Journal of Neuroscience Methods* **143**, 27-32 (2005).
5. A. D. Edelstein *et al.*, Advanced methods of microscope control using μManager software. *JBM* **1**, (2014).
6. Y. Wang *et al.*, EASI-FISH for thick tissue defines lateral hypothalamus spatio-molecular organization. *Cell* **184**, 6361-6377.e6324 (2021).
7. K. Close, Y. He, J. Jeter, G. Ihrke, M. Eddison, Multiplex Detection of Gene Expression in the Intact Drosophila Brain Using Expansion-Assisted Iterative Fluorescence In Situ Hybridization. *JoVE*, e67656 (2025).
8. M. R. Tavakoli *et al.*, Light-microscopy-based connectomic reconstruction of mammalian brain tissue. *Nature* **642**, 398-410 (2025).
